# A general method for the development of multicolor biosensors with large dynamic ranges

**DOI:** 10.1101/2022.11.29.518186

**Authors:** Lars Hellweg, Anna Edenhofer, Lucas Barck, Magnus-Carsten Huppertz, Michelle. S. Frei, Miroslaw Tarnawski, Andrea Bergner, Birgit Koch, Kai Johnsson, Julien Hiblot

## Abstract

Fluorescent biosensors enable to study cell physiology with spatiotemporal resolution, yet most biosensors suffer from relatively low dynamic ranges. Here, we introduce a family of designed Förster Resonance Energy Transfer (FRET) pairs with near quantitative FRET efficiencies based on the reversible interaction of fluorescent proteins with a fluorescently labeled HaloTag. These FRET pairs enabled the straightforward design of biosensors for calcium, ATP and NAD^+^ with unprecedented dynamic ranges. The color of each of these biosensors can be readily tuned by either changing the fluorescent protein or the synthetic fluorophore, which enabled to monitor simultaneously free NAD^+^ in different subcellular compartments upon genotoxic stress. Minimal modifications furthermore allow the readout of these biosensors to be switched to fluorescence intensity, lifetime or bioluminescence. These FRET pairs thus establish a new concept for the development of highly sensitive and tunable biosensors.

**Graphical abstract:** 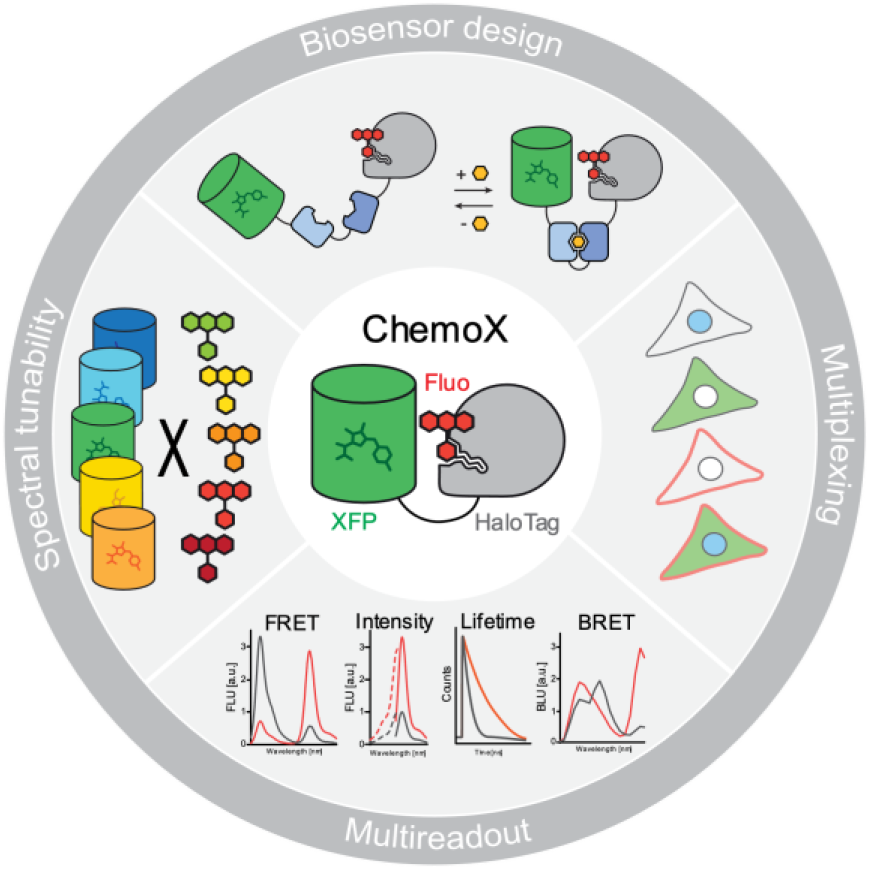

## Introduction

Fluorescent biosensors are powerful tools for the investigation of cellular processes. They allow the real-time monitoring of biological activities, such as changes in metabolite concentration, with subcellular resolution^1^. Biosensors generally consist of two domains: one capable of sensing a biological activity or analyte and a second one translating it into a measurable readout. The readout of current fluorescent biosensors is mostly based on fluorescent proteins (FPs). Biosensors are engineered either by rendering the fluorescence intensity of the FP dependent on the presence of a biological activity^2^ or exploiting the Förster resonance energy transfer (FRET) between two spectrally compatible FPs^1^. In both cases, engineering sensors with large changes of their spectral properties (*i*.*e*. dynamic range) in response to changes of the biological activity of interest often requires laborious optimization, screening a large number of variants^2-5^.

One popular method to develop FRET biosensors is to sandwich a sensing domain between a cyan (CFP) and yellow FP (YFP). These FPs exhibit a large spectral overlap that favor efficient FRET but result in spectral crosstalk which limits their dynamic range while occupying a large part of the visible spectrum. To increase their dynamic range and spectral compatibility with other fluorescent tools, sensors based on green/red (GFP/RFP) or orange/red (OFP/RFP) FRET pairs have been developed but their low FRET efficiencies result too in relatively small dynamic ranges^6-9^. In comparison to FPs, synthetic fluorophores exhibit overall superior photophysical properties, in particular for red-shifted wavelengths^10^. Rhodamines represent the largest class of synthetic fluorophores whose properties were highly optimized for molecular brightness, photostability and fluorogenicity, covering the visible and near-infrared spectrum^10^. On the other hand, self-labeling proteins (SLPs) enable specific labeling in live cells using cell-permeable bio-orthogonal fluorophore substrates in an analogous manner to FPs^11^. SLPs in combination with rhodamines therefore represent appealing candidates as FRET pairs in biosensor design^12^. However, the sole implementation of synthetic fluorophores into the design of biosensors does not give access to large dynamic ranges. In a recent example, the CFP/YFP FRET pair of multiple biosensors was replaced by two SLPs labeled with different rhodamine fluorophores but reached relatively low dynamic ranges^13^.

We hypothesized that engineering a reversible interaction between a FP and fluorescently labeled SLP should enable the straightforward development of FRET biosensors with large dynamic ranges. We thus engineered an interface between a FP FRET donor and a rhodamine-labeled HaloTag^14, 15^ FRET acceptor to reach near quantitative FRET efficiency. By implementing our chemogenetic FRET pairs into the design of biosensors and fine-tuning the FP-HaloTag interface, we developed FRET biosensors for calcium (Ca^2+^), adenosine triphosphate (ATP) and nicotinamide adenine dinucleotide (NAD^+^) with unprecedented dynamic ranges in a straightforward manner. The spectral tunability provided by the HaloTag labeling and by the common beta-barrel architecture of FPs further enabled to readily choose the spectral properties of the sensors such that they could be multiplexed in fluorescence microscopy. Finally, we complete this chemogenetic toolbox by providing simple means to convert FRET biosensors into intensiometric and fluorescence lifetime-based sensors in the far-red range as well as bioluminescent sensors.

## Results

### Chemogenetic FRET pair engineering

To test the FRET between EGFP and fluorescently labeled HaloTag7 (HT7), EGFP was fused directly to the N- or C-terminus of HT7, labeled with the far-red fluorophore silicon rhodamine (SiR) and the fluorescence emission profile was measured in order to evaluate the intra-molecular FRET efficiency of the chemogenetic design (**Fig. S1a**). Fusing EGFP to the N-terminus of HT7 (EGFP-HT7) revealed the highest FRET ratio (2.2 ± 0.1 [mean ± s.d.], **Fig. S1b**). Due to its <u>chemo</u>genetic nature and involving E<u>G</u>FP, this design was named ChemoG1. As EGFP and SiR show only limited spectral overlap (**Fig. S1c**), this high FRET ratio suggested that the two chromophores are in very close proximity. The X-ray structure of ChemoG1 labeled with the fluorophore tetramethylrhodamine (TMR, structurally similar to SiR) (PDB ID: 8B6S, 1.8 Å resolution) confirmed the fluorophore location at the interface between HT7 and EGFP, in close proximity to the EGFP chromophore (15.2 Å distance) (**Fig. 1a & S2a**). The EGFP surface residues Y39, K41 and F223 form a salt bridge with the carboxylate of the fluorophore (K41) and π-stacking interactions with the fluorophore benzyl (Y39) and xanthene (F223) moieties (**Fig. S2a**). Their modification led to drastic loss in FRET (**Fig. S2b**).

**Figure 1.**
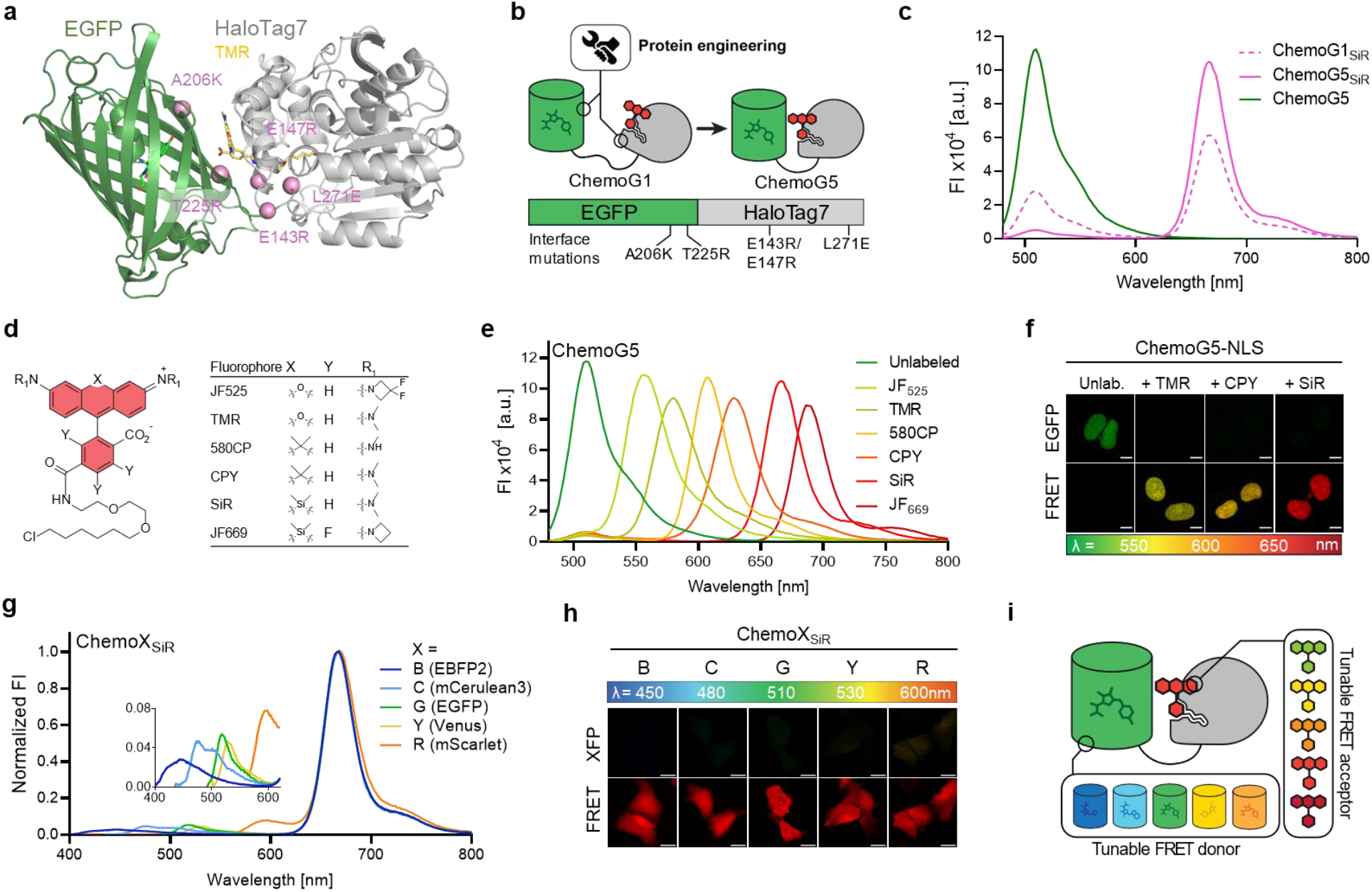
Development of chemogenetic FRET pairs with tunable wavelengths (ChemoX). **a**. Crystal structure of TMR-labeled ChemoG1. ChemoG1 = EGFP-HaloTag7 fusion construct. PDB ID 8B6S, 1.8 Å resolution. Proteins are represented as cartoon, the EGFP chromophore and TMR are shown as sticks. Pink spheres represent the engineered positions at the EGFP and HT7 interface. **b**. Schematic representation of ChemoG1 interface engineering. **c**. Fluorescence intensity (FI) emission spectra of SiR-labeled ChemoG1 (ChemoG1_SiR_) and ChemoG5 (ChemoG5_SiR_) with unlabeled ChemoG5. Represented are the means of 3 technical replicates. **d**. General chemical structure of rhodamine fluorophores. **e**. Fluorescence intensity (FI) emission spectra of ChemoG5 labelled with spectrally distinct rhodamine fluorophores listed in **d**. Represented are the means of 3 technical replicates. **f**. Confocal images of U-2 OS cells expressing ChemoG5 in the nucleus (ChemoG5-NLS) labeled with TMR, CPY, SiR or unlabeled. Shown are the EGFP and FRET channels corresponding to the maximal emission of the respective fluorophores. LUT of EGFP and FRET channels are adjusted to the same values for each condition. Scale bars = 10 µm. **g**. Fluorescence intensity (FI) emission spectra of ChemoX constructs labeled with SiR. Spectra were normalized to the maximum FRET emission. Inlay shows a zoom-in of the FRET donor fluorescence emission. Represented are the means of 3 technical replicates. **h**. Confocal images of ChemoX constructs expressed in U-2 OS cells and labeled with SiR. Shown are the corresponding FP and FRET emission channels. LUT of XFP and FRET channels are adjusted to the same values for each construct. Scale bars = 25 µm. **i**. Schematic representation of the spectral tunability of the ChemoX approach.

We hypothesized that the ChemoG1 conformation observed in the crystal structure exists only transiently in solution and identified interface mutations in order to stabilize it (*i*.*e*. EGFP: A206K, T225R; HT7: E143R, E147R, L271E; **Fig. 1b**). These mutations improved the FRET efficiency in comparison to ChemoG1 in a stepwise manner, generating ChemoG2 to ChemoG5 (**Fig. S2c-d, Table S1**). SiR-labeled ChemoG5 (*i*.*e*. ChemoG5_SiR_) carries all interface mutations and exhibits a near quantitative FRET efficiency (95.8 ± 0.1 %, **Fig. 1c**). The fluorescence intensity of EGFP was not affected by the surface mutations (**Fig. S2e**). The X-ray structure of ChemoG5_TMR_ (PDB ID: 8B6T, 2.0 Å resolution) revealed additional hydrogen bonds (T225R^EGFP^-P170/V173^HT7^) and electrostatic surface modifications (A206K^EGFP^, E143R-E147R^HT7^, L271E^HT7^-R72^EGFP^), likely responsible for the near quantitative FRET (**Fig. S2f-k**). Despite the ionic nature of some of the interactions, the FRET was minimally affected by changes in pH or salt concentration (**Fig. S3a-b**). In U-2 OS cells, the stepwise improvement in FRET of SiR-labeled ChemoG1 to ChemoG5 was confirmed by fluorescence microscopy with a maximum FRET/EGFP ratio of 16.4 ± 2.7 for ChemoG5 that, in turn, was not noticeably affected by changes in the local environment upon expression at different subcellular localizations (**Fig. S4, Table S2**).

HT7 enables tuning of the spectral properties of the FRET acceptor on demand using different fluorophore substrates. Labeling ChemoG5 with different rhodamine fluorophores (**Fig. 1d**) yielded efficient FRET pairs with acceptor maximum fluorescence emission wavelengths ranging from 556 nm (JF_525_) to 686 nm (JF_669_) (FRET efficiencies ≥ 94 %, **Fig. 1e, Table S3**). By expressing ChemoG5 in the nucleus of U-2 OS cells (ChemoG5-NLS), it was possible to confirm the spectral tunability of the FRET acceptor by fluorescence microscopy upon labeling with different rhodamines (**Fig. 1f**). Labeling ChemoG5 with the structurally distinct cyanine fluorophores Cy3 or Cy5, however, resulted in lower FRET compared to the spectrally similar rhodamine fluorophores TMR and SiR, respectively (**Fig. S5a**-**b**). The X-ray structure of Cy3-labeled HT7 (PDB ID: 8B6R, 1.5 Å resolution, **Fig. S5c-d**) revealed a Cy3 conformation at the HT7 surface incompatible with the interactions observed between TMR and EGFP in ChemoG5_TMR_, potentially explaining the weaker FRET.

To further expand the spectral tunability of the ChemoG design, EGFP was exchanged with a blue (EBFP2), cyan (mCerulean3), yellow (Venus) or red (mScarlet) FP creating the chemogenetic FRET constructs ChemoB, ChemoC, ChemoY and ChemoR, respectively. The design is thus named ChemoX, where ‘X’ refers to the color of the respective FP. Transposing the structural features of ChemoG5 to the ChemoX constructs led to optimized ChemoB, ChemoC and ChemoY variants all exhibiting near quantitative FRET efficiencies (≥94 %) upon SiR labeling (**Fig. S6a-c, Table S4**). Attempts to transpose the structural features of ChemoG5 to ChemoR revealed challenging (**Fig. S6d**) probably because of the different phylogenetic origin of mScarlet^16^. Favored by the large spectral overlap between mScarlet and SiR, the initial ChemoR construct nevertheless showed a high FRET efficiency that was further increased by the mutation D201K^mSca^ (91.3 ± 0.3 %, **Fig. S6d-e, Table S4**). The ChemoX palette offers multiple options throughout the visible spectrum (**Fig. 1g**) and displays efficient FRET in cells (FRET ratio > 14) such that the fluorescence emission was almost exclusively observed in the FRET channel (**Fig. 1h, Fig. S6f**). ChemoX thus constitutes a platform of FRET pairs whose colors can be readily chosen by exchanging the fluorescent protein or the synthetic fluorophore (**Fig. 1i**).

### ChemoX-based calcium FRET sensors

We hypothesized that a reversible interaction of FPs with the fluorescently labeled HT7 in ChemoX could enable the development of a new family of fluorescent biosensors. As initial proof of principle and analogously to the calcium (Ca^2+^) sensor yellow cameleon 3.6 (YC 3.6)^17^, we designed a Ca^2+^ sensor in which EGFP and HT7 sandwich the Ca^2+^ binding protein calmodulin (CaM) and its cognate binding peptide M13 (EGFP-CaM/M13-HT7, **Fig. 2a**). CaM and M13 were connected via a poly-proline linker (P30) while short GGS linkers connected FP/HT7 with the sensing domains. The SiR-labeled construct EGFP-CaM/M13-HT7 displayed a large change of fluorescence emission spectrum with increasing concentrations of free Ca^2+^ (**Fig. S7a**), displaying a maximal FRET/EGFP ratio (R) change (^max^ΔR/R_0_) of 22.8 ±0.3, also referred to as dynamic range. The introduction of the previously identified EGFP/HT7 interface mutations (**Table S1**) into the sensor led to a stepwise FRET increase both in absence and in presence of free Ca^2+^ (**Fig. S7b**). This approach revealed key in tuning the dynamic range of the sensor (**Fig. S7c**), allowing us to identify ChemoG-CaM, a Ca^2+^ sensor with the interface mutations A206K^EGFP^ and L271E^HT7^. ChemoG-CaM_SiR_ showed a large dynamic range (^max^ΔR/R_0_ = 36.1 ±1.0 fold) (**Fig. 2b, Table S5**) with a half maximal response (C50) at 179 nM free Ca^2+^ concentration (95% confidence interval [CI]: 173-185 nM), similar to YC 3.6 (243 nM, 95% CI: 233-251 nM). Furthermore, the C50 of the sensors was independent of the number of interface mutations (**Fig. S7d & Table S5**). The ChemoX-CaM sensor color can be readily tuned by either changing the synthetic fluorophore or the FP. All generated sensors exhibited large dynamic ranges, outperforming YC 3.6 with the exception of the red version of the sensor (**Fig. 2c-d, Fig. S7e-j, Table S5 & S6**). Replacing mScarlet by mRuby2 led to a sensor with increased dynamic range (^max^ΔR/R_0_ = 3.4. ±0.1 fold), named ChemoR-CaM_SiR_ (**Fig. S8, Table S5**). From here, ChemoR biosensors will always be based on mRuby2 as it revealed more potent in yielding biosensors with larger dynamic ranges. Noteworthy, all sensors revealed similar C50s underlining the straightforwardness of the approach (**Fig. S7g & j**). As previously described for YC3.6^17^, ChemoG-CaM_SiR_ revealed sensitive to pH changes (**Fig. S7k)**. This sensitivity might come from the sensing domain since ChemoG1-5 revealed mostly pH insensitive over the same range (**Fig. S3**).

**Figure 2.**
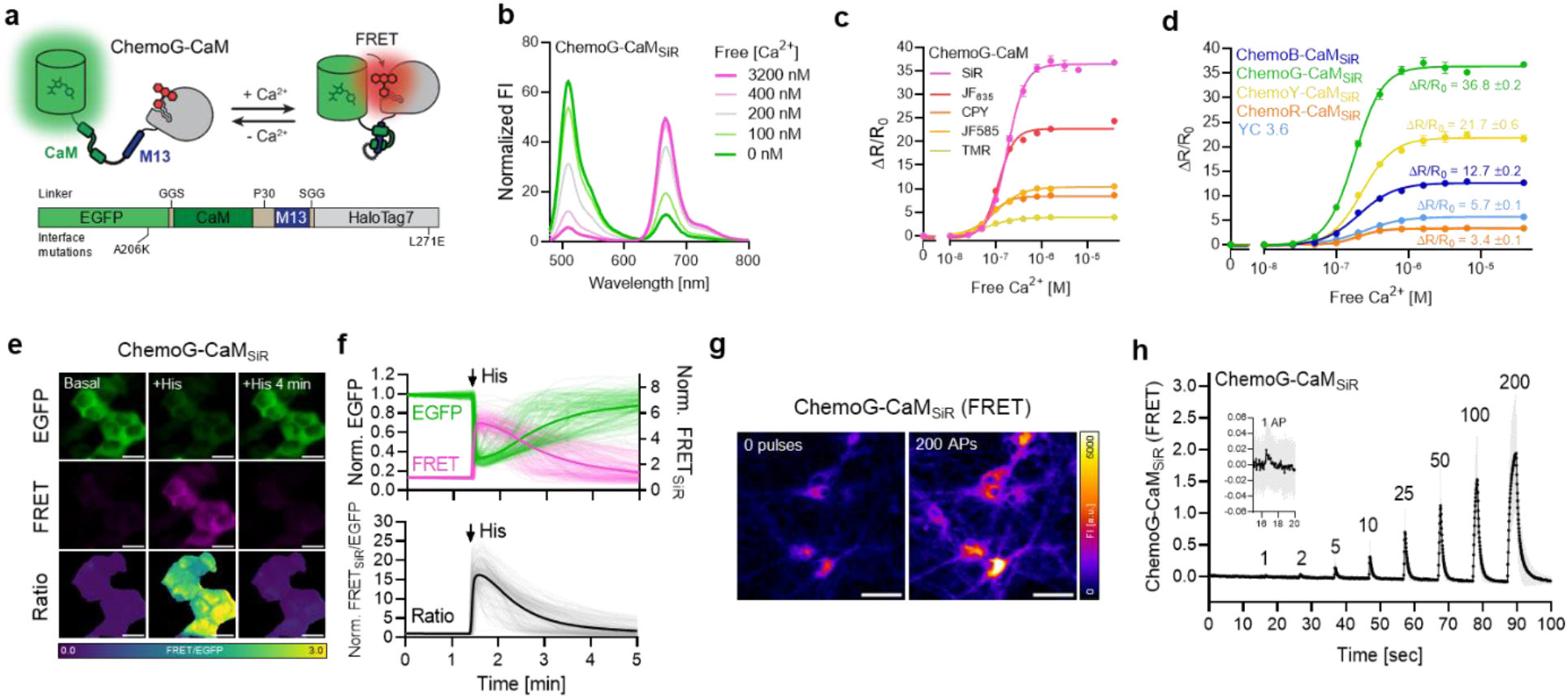
Development of ratiometric calcium sensors based on ChemoX. **a**. Schematic representation of ChemoG-CaM. **b**. Normalized fluorescence intensity (FI) emission spectra of SiR-labeled ChemoG-CaM at different concentrations of free calcium. Represented are the means of 3 technical replicates **c**. Calcium titration curves of ChemoG-CaM labeled with different fluorophores. Represented are the means ±s.d. of the FRET/EGFP ratio changes (ΔR/R_0_) (n = 3 technical replicates). ΔR/R_0_ and C50 are summarized in **Table S6. d**. Calcium titration curves of ChemoX-CaM_SiR_ and YC 3.6. Data represented as in **c** (n = 3 technical replicates). ΔR/R are indicated and summarized together with the C50 in **Table S5. e**. Widefield images of HeLa Kyoto cells transiently expressing ChemoG-CaM labeled with SiR. Shown are the EGFP channel, the FRET channel and the ratio image of both channels (FRET/EGFP) in pseudocolor (LUT = mpl-viridis). Cells were treated with 10 µM histamine. The images represent cells at basal conditions before the addition of histamine (basal), 15 seconds after the addition of histamine (+his) and 4 min after the addition of histamine (+his 4 min). Scale bars = 25 µm. **f**. Time course measurement of ChemoG-CaM_SiR_ fluorescence intensity (FI) in HeLa Kyoto cells. Represented are the EGFP and FRET channel (*upper panel*) and FRET/EGFP ratio normalized to 1 at t = 0 min (*lower panel*). Cells were treated with 10 µM histamine at the time point indicated with an arrow. Experiments as explained in panel **e**. (n = 161 cells from 3 biological replicates). Represented are the means (solid line) plus traces of the individual cells (dim lines). **g**. Widefield images of hippocampal neurons isolated from rats expressing cytosolic ChemoG-CaM labeled with SiR at different stimulation intensities. Neurons were stimulated with an electric field corresponding to 0 or 200 action potentials (APs). The fluorescence intensity (FI) of the SiR FRET channel is represented in pseudocolor (LUT = Fire). Scale bars = 50 µm. **h**. Time course measurement of ChemoG-CaM_SiR_ fluorescence intensity in hippocampal neurons isolated from rats. Represented is the FRET fluorescence intensity change (ΔFI/FI_0_) upon electric field stimulation. Experiments conducted as explained in **g**. (n = 61 cells from 3 biological replicates). Number of action potentials (AP) are indicated. Represented is the mean (line) ±s.d. (shade area).

Next, we confirmed the performance of ChemoG-CaM_SiR_ in HeLa cells, where the sensor showed a large FRET increase (∆R/R_0_ = 16.8 ±3.9 fold) upon histamine-induced Ca^2+^ influx into the cytosol (**Fig. 2e-f**). Finally, considering the importance of Ca^2+^ transients in neurobiology, AAV-delivered ChemoG-CaM_SiR_ was characterized in rat primary hippocampal neurons. The sensor displayed a maximum FRET_SiR_ increase (∆FI/FI_0_) of 206 ±90 % upon electric field stimulation (200 action potentials (APs)) and was able to detect as low as one action potential (**Fig. 2g-h**). While ChemoG-CaM_SiR_ is inferior compared to the GCamP8 series^18^, it’s maximum ∆FI/FI_0_ is comparable to the recently developed HaloCaMP1a and 1b in combination with JF_63519_.

### ChemoX-based ATP FRET sensors

Adenosine triphosphate (ATP) is essential for cellular energy homeostasis^20^ and plays important roles in signaling processes^21^. However, biosensors currently available to investigate intracellular changes in free ATP show limited dynamic ranges and are spectrally restricted^22-24^. We therefore developed a ChemoG-based ATP biosensor by exchanging the FRET pair mseCFP and Venus of the ATP biosensor ATeam 1.03^22^ with EGFP and HT7, respectively (**Fig. 3a**). The dynamic range of the construct EGFP-F_O_-F_1_-HT7_SiR_ was optimized by the introduction of EGFP/HT7 interface mutations A206K^EGFP^ and L271E^HT7^ obtaining ChemoG-ATP_SiR_ with a ^max^ΔR/R_0_ of 12.1 ±0.4 fold (**Fig. 3b, Fig. S9a-b, Table S7**). The sensor responds to millimolar ATP concentration (C50 = 2.33 mM, 95%CI: 2.18-2.53 mM) with high selectivity over other nucleotides (**Fig. S9c**). Consistent with previous observations for ATeam 1.0322, the sensor is sensitive to changes in temperature and pH (**Fig. S9d-e**). Based on the ChemoX design, a color palette of ATP sensors was developed (**Fig. S9f**-**g**), which exhibits dynamic ranges larger or similar to the one of ATeam 1.03 (**Fig. 3c, Table S7**).

**Figure 3.**
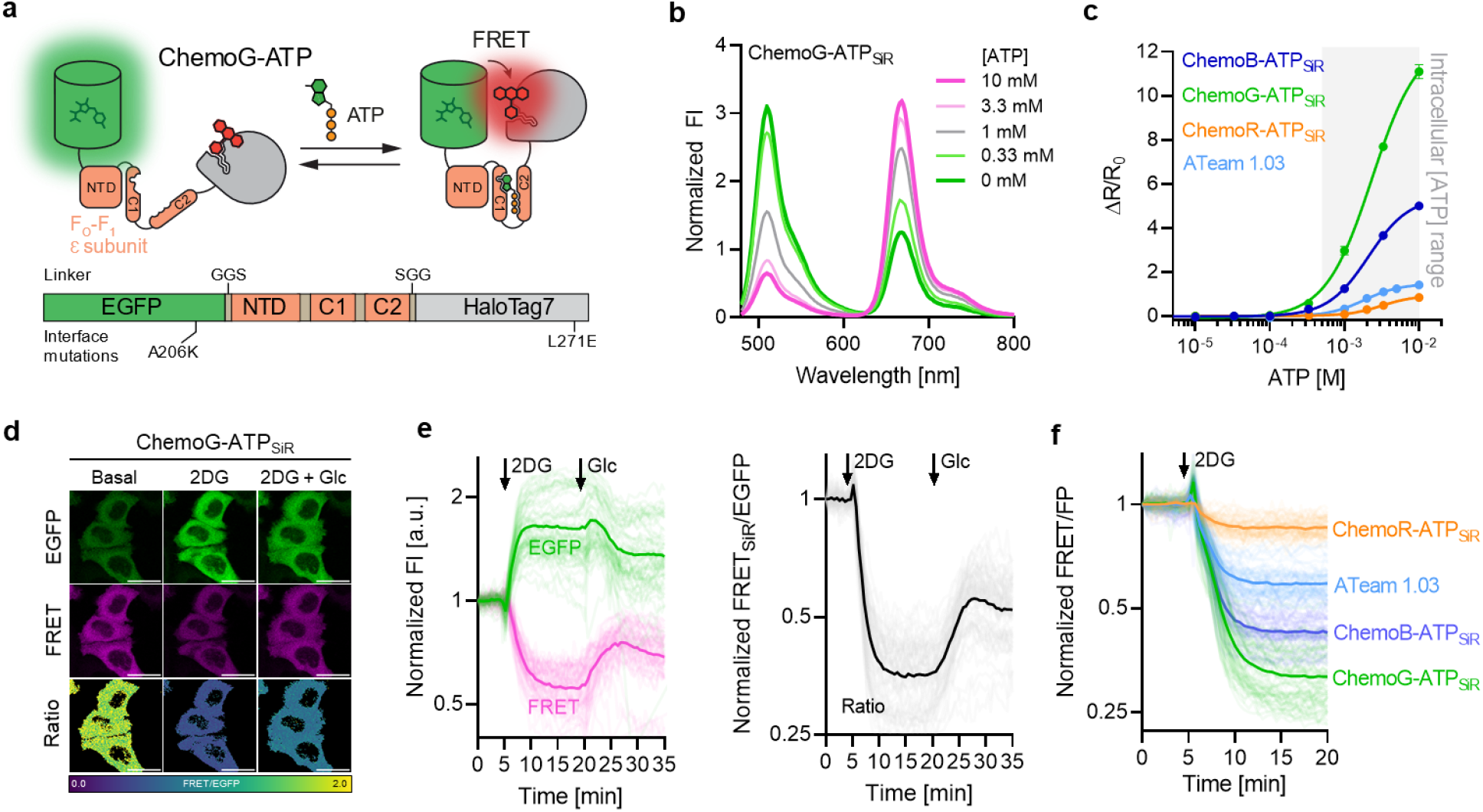
Development of ratiometric ATP sensors based on ChemoX. **a**. Schematic representation of ChemoG-ATP. **b**. Fluorescence intensity (FI) emission spectra of SiR-labeled ChemoG-ATP at different ATP concentrations. Represented are the means of 3 technical replicates. **c**. ATP titration curves of ChemoX-ATP_SiR_ sensors. Represented are the means ±s.d. of the FRET/EGFP ratio changes (ΔR/R_0_) (n = 3 technical replicates). The intracellular ATP concentration range is indicated with a grey box. ΔR/R_0_ and C50 values are summarized in **Table S7. d**. Confocal images of HeLa Kyoto cells expressing ChemoG-ATP labeled with SiR. Shown are the EGFP channel, the FRET channel and the ratio image of both channels (FRET/EGFP) in pseudocolor (LUT = mpl-viridis). Cells were treated at t = 5 min with 10 mM 2-deoxy-D-glucose (2DG). At t = 20 min, 20 mM glucose (Glc) was added to the cells until the end of the experiment (t = 35 min, 2DG + Glc). Scale bars = 25 µm. **e**. Time course measurement of ChemoG-ATP_SiR_ fluorescence intensity (FI) in HeLa Kyoto cells. Represented are the EGFP and FRET channel (*left panel*) and FRET/EGFP ratio (*right panel*) normalized to 1 at t = 0 min. Cells were treated with 10 mM 2DG and subsequently with 20 mM Glc at time points indicated with arrows. Experiments as explained in panel **d**. (n = 59 cells from 3 biological replicates). Represented are the means (solid lines) and traces of the individual cells (dim lines). **f**. Time course measurement of ChemoB-ATP_SiR_ (n = 58 cells), ChemoG-ATP_SiR_ (n = 63 cells), ChemoR-ATP_SiR_ (n = 52 cells) and ATeam 1.03 (n = 59 cells) fluorescence intensity in HeLa Kyoto cells. Represented is the FRET/FP ratio upon treatment with 10 mM 2DG. Ratios are normalized to 1 at t = 0 min. Addition of 2DG is indicated with an arrow. Represented are the means (line) and single cell traces (dim lines) from 3 biological replicates.

ChemoG-ATP_SiR_ was expressed in the cytosol of HeLa Kyoto cells where treatment with the glycolysis inhibitor 2-deoxy-D-glucose (2DG) led to a strong FRET/EGFP decrease (ΔR/R_0_ = -67.9 ±6.3 %) that could be partially compensated by perfusing high concentrations of glucose (**Fig. 3d-e**). The FRET/EGFP ratio briefly increased immediately after addition of 2DG, which we attribute to the transient change in temperature. Similarly, ChemoB-ATP_SiR_ and ChemoR-ATP_SiR_ reported a 2DG-induced decrease in cytosolic ATP (ChemoB-ATP_SiR_ ΔR/R_0_ = -57.5 ±4.7 %, ChemoR-ATP_SiR_ ΔR/R_0_ = -14.5 ±4.9 %), which we compared to ATeam 1.03 (ΔR/R_0_ = -41.1 ± 5.1 %) (**Fig. 3f**). While ChemoB-ATP_SiR_ revealed more potent than ATeam 1.03 in translating intracellular ATP concentration changes into FRET changes, this was not the case for ChemoR-ATP_SiR,_ consistent with in vitro titrations (**Fig. 3c**).

### ChemoX-based NAD^+^ FRET sensors

Nicotinamide dinucleotide (NAD^+^) is highly compartmentalized within cells^25^ and the regulation of its subcellular pools plays an important role in many biological processes^26, 27^. However, monitoring changes in intracellular NAD^+^ at multiple subcellular locations is limited by the spectral incompatibility and selectivity of current biosensors^28-30^. Inspired by the recently developed NAD^+^ sensor based on DNA ligase A (LigA)^29, 31^, we developed a sensor based on the catalytically inactive LigA from *Thermus thermophilus* (*tt*LigA^D^ = ttLigA^K118L-D289N^), sandwiching the sensing domain between EGFP and HT7 (**Fig. 4a**). The sensor’s C50 was optimized by implementing the mutations Y226W and V292A in *tt*LigA^D^ (**Fig. S10a-b**) and the dynamic range optimized by implementing selected EGFP/HT7 interface mutations (*i*.*e*. A206K-T225R^EGFP^ and L271E^HT7^, **Fig. S10c-d**). The resulting ChemoG-NAD_SiR_ sensor presents a high dynamic range (^max^ΔR/R_0_ = 34.7 ± 0.4 fold) and a C50 (200 µM, 95% CI: 182-220 µM) in the range of expected intracellular NAD+ concentration (50-400 µM depending on the compartment^28, 29^) (**Fig. 4b, Table S8**). Furthermore, the sensor did not respond to increasing concentrations of NAD^+^ precursors or structurally related molecules alone (**Fig. S10e**). However, NAD^+^ titrations in presence of certain NAD^+^ precursors or adenine nucleotides did affect the FRET ratio of the sensor (**Fig. S10f-g**). Since AMP, ADP and ATP affect the sensor response for NAD^+^ to the same extent; a relative change in the ATP/ADP/AMP ratio should not affect the response of the sensor in cells, but only a change in the total concentration of the adenosine nucleotide pool. Finally, temperature and pH moderately influenced the sensor’s response (**Fig. S10h-i**). The spectral properties of the sensor can be readily tuned based on the ChemoX design yielding a color palette of NAD^+^ sensors including ChemoB-NAD_SiR_ (^max^ΔR/R_0_ = 11.2 ±0.1 fold) and ChemoR-NAD_SiR_ (^max^ΔR/R_0_ = 3.0 ±0.1 fold) (**Fig. 4c-d, Fig. S11, Table S8**).

**Figure 4.**
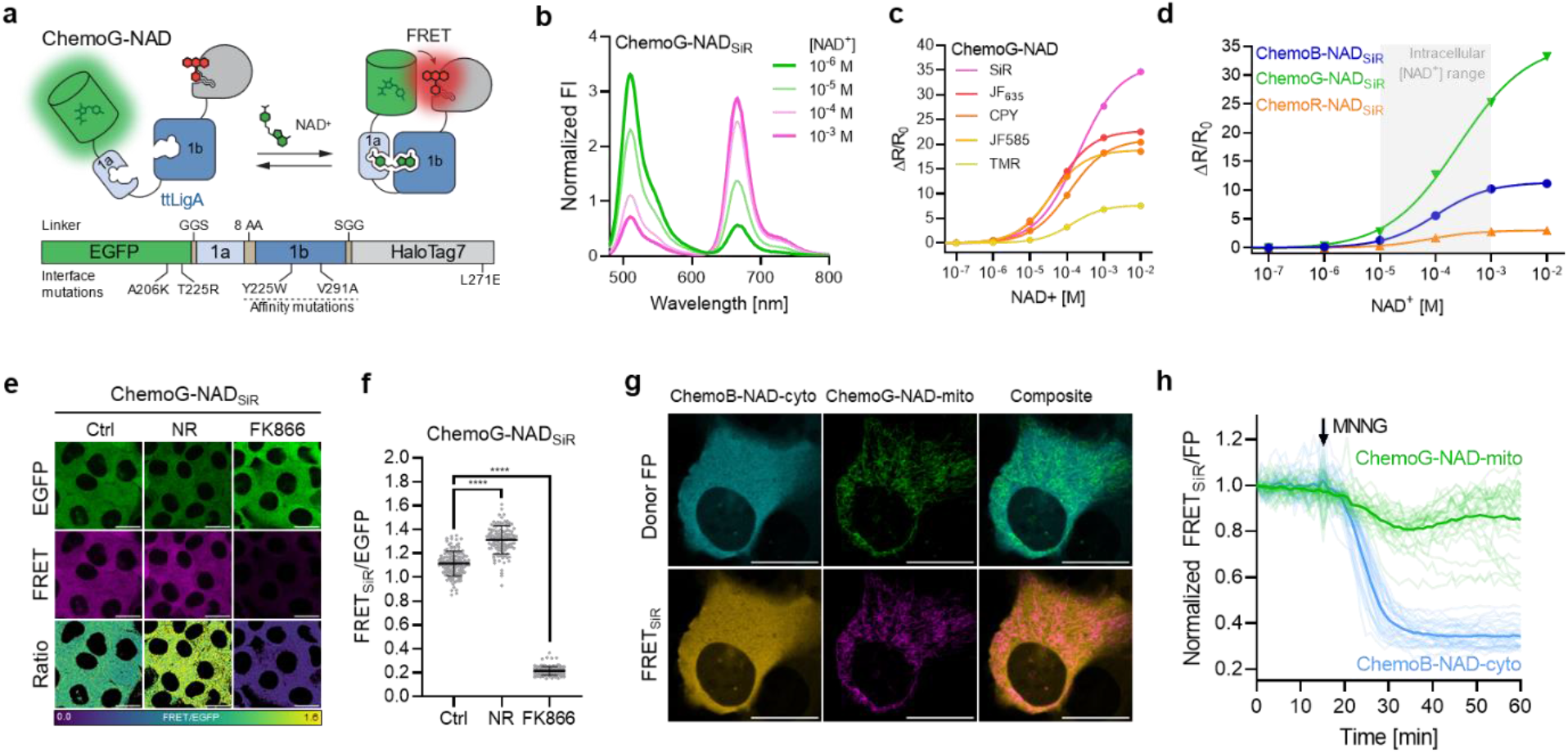
Multiplexing subcellular NAD^+^ pools using ChemoX-NAD biosensors. **a**. Schematic representation of ChemoG-NAD. **b**. Normalized fluorescence intensity (FI) emission spectra of SiR-labeled ChemoG-NAD at different NAD^+^ concentrations. Represented are the means of 3 technical replicates. **c**. NAD^+^ titration curves of ChemoG-NAD labeled with different fluorophores. Represented are the means ±s.d. of the FRET/EGFP ratio changes (ΔR/R_0_) (n = technical 3 replicates). ΔR/R_0_ and C50 values are summarized in **Table S8. d**. NAD^+^ titration curves of ChemoX-NAD_SiR_ biosensors. Represented are the means ±s.d. of the FRET/EGFP ratio changes (ΔR/R_0_) (n = 3 technical replicates). The intracellular free NAD^+^ concentration range is indicated with a grey box. ΔR/R_0_ and C50 values are summarized in **Table S8. e**. Confocal images of U-2 OS cells expressing ChemoG-NAD labeled with SiR. Shown are the EGFP channel, the FRET channel and the ratio image of both channels (FRET/EGFP) in pseudocolor (LUT = mpl-viridis). Prior to imaging, cells were treated for 24h with either DMSO (Ctrl), 100 nM FK866 or 1 mM NR. Scale bars = 25 µm. **f**. Dot plots representing the FRET/EGFP ratios of ChemoG-NAD_SiR_ expressed in U-2 OS cells treated as described in **d**. (n ≥ 117 cells for each condition from 3 biological replicates). p-values are given based on unpaired t-test with Welch’s correction (**** p < 0.0001). Represented are the means ±s.d. **g**. Confocal image of U-2 OS cell co-expressing ChemoB-NAD-cyto and ChemoG-NAD-mito labeled with SiR. Shown are the FRET donor FP channels, the FRET channels and the composites of FP or FRET channels of both sensors pseudocolored (EBFP2 (cyan), EGFP (green), EBFP2-FRET (orange), EGFP-FRET (magenta)). The brightness of the donor and FRET channels were adjusted individually to show potential crosstalk between the channels. Scale bars = 25 µm. **h**. Time course measurement of ChemoB-NAD-cyto (cytosol) and ChemoG-NAD-mito (mitochondria) fluorescence intensity co-expressed in U-2 OS cells and labeled with SiR. Represented are the means of the FRET/FP ratios (line) and single cell traces (dim lines) normalized to 1 at t = 0 min. Addition of 100 µM MNNG is indicated with an arrow (n = 28 cells from 4 biological replicates).

The impact of the nicotinamide phosphoribosyltransferase (NAMPT) inhibitor FK866 and the NAD^+^ precursor nicotinamide riboside (NR) on cellular free NAD^+^ concentration was next studied in U-2 OS cells using cytosolically expressed ChemoG-NAD_SiR_. In fluorescence microscopy experiments, the FK866-induced NAD^+^ depletion led to a strong FRET ratio decrease (ΔR/R_0_ = -78.9 ±3.6 %), whereas treatment with NR increased the intracellular NAD^+^ as indicated by a significant FRET ratio increase (ΔR/R_0_ = 28.4 ±11.4 %) (**Fig. 4e-f**). The sensor was able to translate NAD^+^ changes with similar trends in the nucleus and mitochondria of U-2 OS cells (**Fig. S12**). While ChemoB-NAD_SiR_ showed similar performances to ChemoG-NAD_SiR_ in cells (FK866: ΔR/R_0_ = -66.2 ±8.5 %; NR: ΔR/R_0_ = 17.3 ±21.1 %), ChemoR-NAD_SiR_ presents a reduced sensitivity (FK866: ΔR/R_0_ = -21.3 ±3.2 %; NR: ΔR/R_0_ = 2.5 ±4.5 %) (**Fig. S13**), which is nevertheless comparable to previously published NAD^+^ biosensors ^28, 29^ and of interest for multiplexing purposes.

By using two spectrally compatible biosensors of the ChemoX-NAD palette, we were able to monitor for the first time the fluctuation of free NAD^+^ in real-time in two subcellular compartments of U-2 OS cells, *i*.*e*. ChemoB-NAD_SiR_ and ChemoG-NAD_SiR_ located in the cytosol and mitochondria, respectively. The sensors showed negligible cross talk between the different emission channels (**Fig. 4g**). Upon treatment of the cells with *N*-methyl-*N*’-nitro-*N*-nitrosoguanidine (MNNG), an alkylating agent that is known to lead to hyperactivation of the NAD^+^-consuming enzyme PARP1^32^, we observed a rapid depletion of cytosolic NAD^+^while mitochondrial NAD^+^ showed a slower, less pronounced decrease (**Fig. 4h, Fig S13a**). A similar trend was observed using ChemoB-NAD_SiR_ and ChemoG-NAD_SiR_ co-expressed in the nucleus and mitochondria, respectively (**Fig. S14b-d**). In cytosol and nucleus, NAD^+^ depletion was observed only 5 minutes after addition of MNNG and completed in less than 30 minutes. Previous observations already showed that, during genotoxic stress induced by alkylating agents, mitochondrial NAD^+^ is maintained longer than cytosolic NAD^+ 33^. However, the multiplexing of biosensors here revealed that some cells experienced a decrease in mitochondrial NAD^+^ comparable to the decrease observed in the cytosol and nucleus, indicating a cell-to-cell heterogeneity in response to genotoxic stress.

### ChemoX-based intensiometric and lifetime sensors for NAD^+^

ChemoX FRET sensors involve rhodamine fluorophores whose photophysical properties can be influenced by the environment^34^. We therefore hypothesized that ChemoX sensors could be converted into intensiometric and fluorescence lifetime-based sensors as the sensor’s conformational change would likely affect the environment of the rhodamine and thereby its photophysical properties (**Fig. S15a**). The fluorescence intensity (FI) of SiR in the context of ChemoG-NAD_SiR_ already showed a 28.0 ± 1.9 % increase upon NAD^+^ addition (**Fig. S15b**) while the FI of isolated HT7_SiR_ remained mostly unaffected by NAD^+^ (**Fig. S15c**). The sensor’s dynamic range (^max^ΔFI/FI_0_) was further improved by introducing the mutation P174W^HT7^, which is known to reduce the FI of SiR on HT7^34^. EGFP was additionally replaced by the non-fluorescent ShadowG^35^ since it serves only as scaffolding element, creating the sensor ChemoD-NAD (D standing for dark) (**Fig. 5a, Fig. S15d-e**). ChemoD-NAD_SiR_ exhibits a ^max^ΔFI/FI_0_ of 161 ± 5.0 %, reaching a maximum FI comparable to isolated HT7_SiR_ with a C50 at 32.7 µM (95% CI: 24.3-42.1 µM) (**Fig. 5b, Fig. S15d**). The spectral properties of ChemoD-NAD can be adapted using different fluorophore substrates among which JF_635_ yielded the highest ^max^ΔFI/FI_0_ with 226.6 ± 4.3 % (**Fig. 5c, Table S9**). The sensor FI intensity change probably occurs through dequenching of the fluorophore induced by conformational change and interaction with ShadowG. The mutation P174W^HT7^ was previously reported to mostly affect the quantum yield and therefore the fluorescence lifetime of rhodamines at the HT7 surface^34^. We hypothesized that the sensor might also present changes in fluorescence lifetime and could be used for fluorescence lifetime imaging microscopy (FLIM). Indeed, in vitro, the fluorescence lifetime (τ) of ChemoD-NAD_SiR_ increased in the presence of NAD^+^ from 2.21 ± 0.01 ns to 3.37 ± 0.01 ns (**Fig. 5d, Fig. S15f-g**) offering a dynamic range (^max^Δτ) of 1.16 ± 0.01 ns and a C50 at 22.4 µM (95% CI: 20.6-24.4 µM). ChemoD-NAD can also be combined with other fluorophores, notably CPY that showed the largest ^max^Δτ with 1.18 ± 0.01 ns (**Fig. 5e-f, Fig. S15h-i, Table S10**).

**Figure 5.**
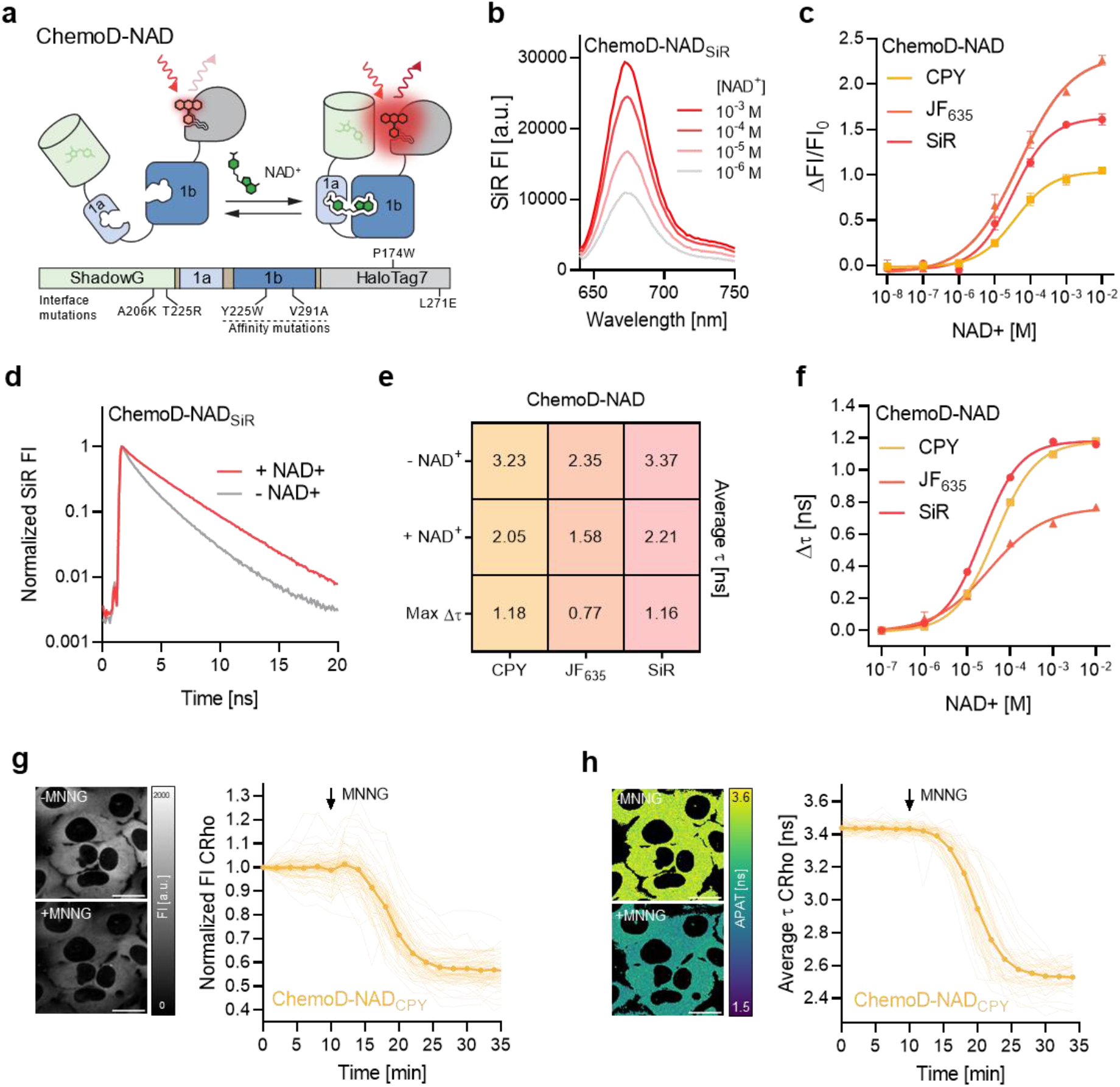
Development of far-red NAD^+^ biosensors based on fluorescence intensity and fluorescence lifetime. **a**. Schematic representation of ChemoD-NAD. **b**. Fluorescence intensity (FI) emission spectra of SiR-labeled ChemoD-NAD at different NAD^+^ concentrations. Represented are the means of 3 technical replicates **c**. NAD^+^ titration curves of ChemoD-NAD labeled with CPY, JF_635_ or SiR. Represented are the means ±s.d. of the fluorescence intensity changes (ΔF/F_0_) (n = 3 technical replicates). ΔF/F_0_ and C50 values are summarized in **Table S9. d**. Fluorescence lifetime decay curves of ChemoD-NAD_SiR_ in presence of 1 mM (+ NAD^+^) or absence of NAD^+^ (-NAD^+^). **e**. Intensity-weighted average fluorescence lifetimes (τ) of ChemoD-NAD labeled with different fluorophores. Represented are the mean intensity-weighted average fluorescence lifetimes in presence of 1 mM (+ NAD^+^) or absence of NAD^+^ (-NAD^+^) and the change in lifetime (Δτ) (n = 3 technical replicates). **f**. NAD^+^ titration curves of ChemoD-NAD labeled with CPY, JF_635_ or SiR. Represented are the means ±s.d. of the intensity-weighted average fluorescence lifetime changes (Δτ) (n = 3 technical replicates). Δτ and C50 values are summarized in **Table S10. g-h**. Confocal images of U-2 OS cells expressing ChemoD-NAD labeled with CPY. Images are representative snapshots of the CPY fluorescence intensity (FI) channel (**g**) or average photon arrival time (APAT) (**h**) before (-MNNG) and after 100 µM MNNG (+ MNNG) treatment. Time course measurements of ChemoD-NAD_CPY_ fluorescence intensity normalized to 1 at t= 0 min (**g**, n = 86 cells from 3 biological replicates) and intensity-weighted average fluorescence lifetime (**h**, n = 55 cells from 3 biological replicates) in U-2 OS cells are shown next to the confocal images corresponding to the same treatments. Represented are the means (line) and traces of single cells (dim lines). Addition of 100 µM MNNG is indicated with an arrow. Scale bars = 25 µm.

Upon treatment of U-2 OS cells with MNNG, ChemoD-NAD_CPY_ was able to monitor in real-time the depletion of intracellular NAD^+^ (**Fig. 5g-h**), resulting in a FI decrease of 43.0 ± 6.1 % and a τ decrease of 0.91 ±0.08 ns. Similar trends could be observed with ChemoD-NAD_SiR_ and Chemo-NAD_JF635_ (**Fig. S15j-m**). Consistent with the *in vitro* titrations (**Fig. 5c**+**f**), ChemoD-NAD_JF635_ showed the largest change in FI (ΔFI/FI_0_ = -54.2 ± 5.1 %) but the smallest change in τ (Δτ = -0.20 ± 0.08 ns) (**Fig. S15l-m**).

### ChemoX-based bioluminescent sensors

For high-throughput screening purposes, bioluminescent sensors are particularly interesting which motivated us to further develop bioluminescent ChemoX-based sensors. Inspired by the Nano-lantern design^36, 37^, we fused a circularly permuted NanoLuc to the N-terminus of ChemoG-NAD, which gave rise to ChemoL-NAD (L for luminescent), a luminescent BRET-FRET-based sensor for NAD^+^ (**Fig. 6a**). ChemoL biosensors were combined with CPY, since the luminescence reader used could not detect emission wavelengths >650 nm. ChemoL-NAD_CPY_ exhibits a large dynamic range (^max^ΔR/R_0_, R = BRET-FRET_CPY_/EGFP) of 7.5 ± 0.1 fold and a C50 of 60.5 µM (95% CI: 57.3-63.9 µM) (**Fig. 6b-c**). In U-2 OS, ChemoL-NAD_CPY_ was able to detect free NAD^+^ concentration changes induced by treatments with FK866 and NR in the cytosol as well as in the nucleus and mitochondria (**Fig. 6d & S16a-b**). Based on the large ratio changes and excellent Z’ factors of ChemoL-NAD_CPY_ in different subcellular compartments upon treatment with FK866 (cytosol Z’ = 0.76, nucleus Z’ = 0.79, mitochondria Z’ = 0.52), the sensor represents a powerful tool to be used in high-throughput screenings for compounds influencing free intracellular NAD^+^. Furthermore, the sensor was able to report the compensation of FK866-induced NAD^+^ depletion by simultaneous addition of NR and might therefore be useful for screening of compounds replenishing intracellular NAD^+^.

**Figure 6.**
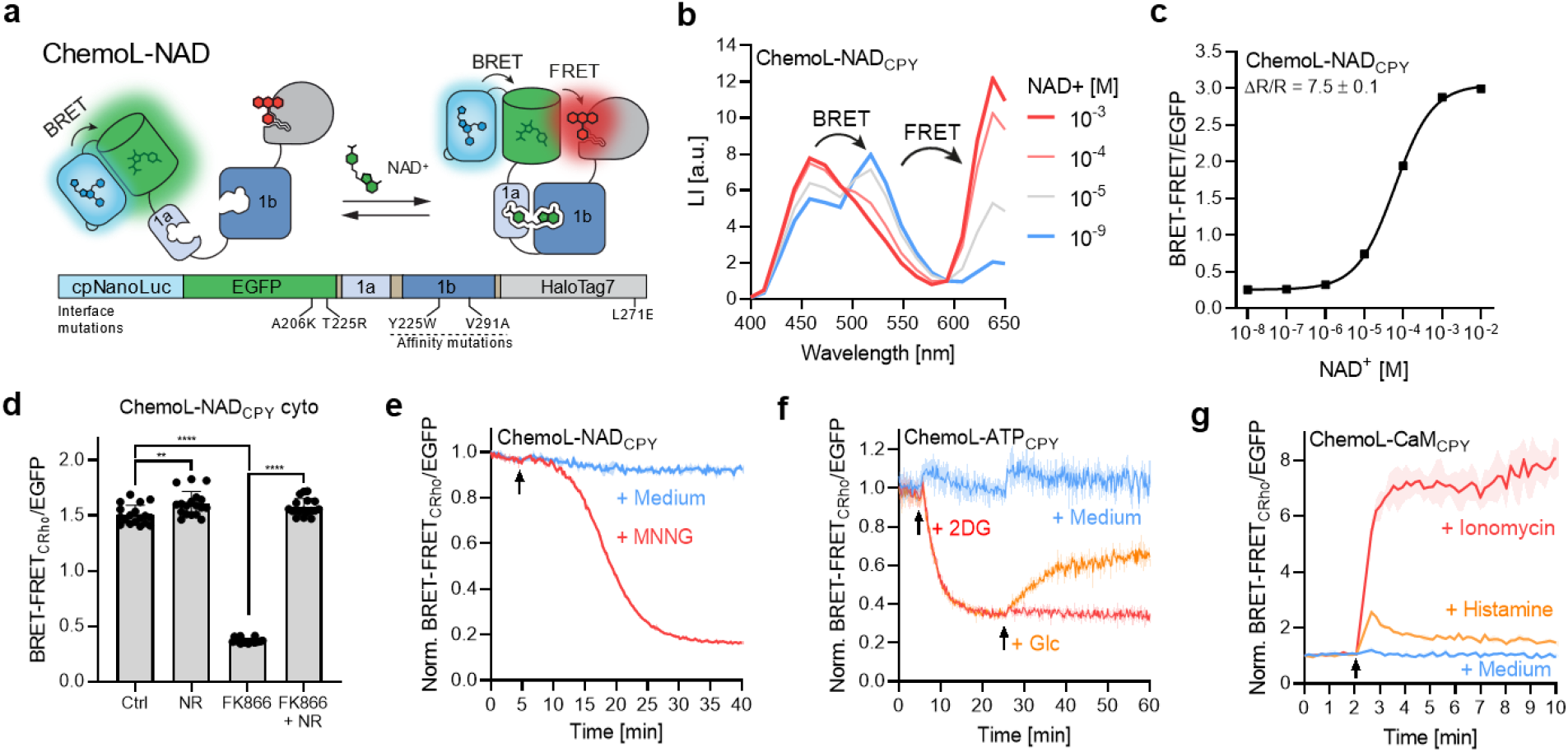
Conversion of ChemoG-based biosensors into luminescent ChemoL biosensors. **a**. Schematic representation of ChemoL-NAD. **b**. Luminescent intensity emission (LI) spectra of CPY-labeled ChemoL-NAD at different NAD^+^ concentrations. Represented are the means of 3 technical replicates. **c**. NAD^+^ titration curve of ChemoL-NAD_CPY_. Represented are the mean BRET-FRET/EGFP luminescence ratios (*i*.*e*. BRET-FRET_CPY_/EGFP) ±s.d of 3 technical replicates. **d**. ChemoL-NAD_CPY_ BRET-FRET/EGFP ratios in U-2 OS cells upon treatment for 24 h with DMSO (Ctrl), 1 mM NR, 100 nM FK866 or 100nM FK866 and 1 mM NR. Represented are the means ±s.d. and single well ratios (circles) (n = 18 wells per condition from 3 biological replicates). p-values are given based on unpaired t-test with Welch’s correction (** p < 0.01, **** p<0.0001). **e**. Time course measurement of ChemoL-NAD_CPY_ expressed in U-2 OS cells. Represented are the means of the BRET-FRET/EGFP ratios (line) ±s.d. (shade areas) normalized to 1 at t = 0 min. Cells were untreated (+ medium) or treated with 100 µM MNNG (+ MNNG) at t = 5 min indicated with an arrow (n = 3 wells from one representative biological replicate; two additional biological replicates can be found in **Fig. S17**). **f**. Time course measurement of ChemoL-ATP_CPY_ expressed HeLa Kyoto cells. Represented are the BRET-FRET/EGFP ratios normalized to 1 at t = 0 min. Cells were untreated (medium), treated with 10 mM 2DG at t = 5 min (red and orange) and additionally treated with 20 mM glucose at t = 25 min (orange). Addition of medium, 2DG and Glc is indicated with an arrow (n = 3 wells from one representative biological replicate; two additional biological replicates can be found in **Fig. S17**). **g**. Time course measurement of ChemoL-CaM_CPY_ expressed HeLa Kyoto cells. Represented are the BRET-FRET/EGFP ratios normalized to 1 at t = 0 min. Cells were untreated, treated with 10 µM histamine or 1 µM ionomycin at t = 2 min (n = 3 wells from one representative biological replicate; two additional biological replicates can be found in **Fig. S17**). Addition of drugs is indicated with an arrow.

Similar to ChemoL-NAD, luminescent sensors for Ca^2+^ and ATP were developed, namely ChemoL-CaM_CPY_ (^max^ΔR/R_0_ = 9.1 ± 0.1 fold; C50 = 96.6 nM, 95% CI: 91.0-102.7 nM, **Fig. S16c-d**) and ChemoL-ATP_CPY_ (^max^ΔR/R_0_ = 6.1 ± 0.1 fold; C50 = 1.00 mM, 95% CI: 0.95-1.05 mM, **Fig. S16e-f**). Using the ChemoL-based biosensors, it was possible to monitor in real-time intracellular changes of NAD^+^, ATP and Ca^2+^ upon different drug treatments (**Fig. 6e-g, Fig. S17a-i**). All sensors showed large ratio changes for the respective drug treatments comparable to the responses observed with FRET-based sensors (ChemoL-NAD_CPY_ ΔR/R_0_ = -74.0 ±4.4 % [MNNG]; ChemoL-ATP_CPY_ ΔR/R_0_ = -70.2 ±6.0 % [2DG]; ChemoL-CaM_CPY_ ΔR/R_0_ = 7.4 ±0.5 times [Ionomycin]) (**Fig. 2f, Fig. 3e, Fig. 4h**).

## Discussion

FP-based FRET biosensors often suffer from small dynamic ranges. Previous strategies to improve the dynamic range of FRET biosensors involved the use of dimerizing domains^38^, screening for optimized linkers^3, 39^ or changing the orientation of the FRET pair^22, 40^. Despite some success, these approaches often require a large number of variants to be screened, which hinders the straightforward development of effective sensors. By engineering an interaction interface between a FP and the rhodamine-labeled HaloTag, we developed FRET pairs (ChemoX) with near-quantitative FRET efficiency. Testing not more than 5 combinations of interface mutations between FP and HaloTag, biosensors with unprecedented dynamic ranges for Ca^2+^ (36.1 fold), ATP (12.1 fold) and NAD^+^ (34.7 fold) were obtained, as compared to similar FRET-based biosensors^3, 17, 22, 28, 40, 41^. Specifically, it was possible to increase the dynamic range of existing biosensors for Ca^2+^ (YC 3.6) and ATP (ATeam 1.03) by 6.3 and 8.6 fold, respectively, by exchanging the CFP/YFP FRET pair with the ChemoG_SiR_ FRET pair, in addition to shifting the fluorescence excitation and emission to longer wavelengths. To facilitate the development of ChemoX-based biosensors by others, we provide a guideline in Supplementary information (**Fig. S18-20, Extended notes 1-3**). Interface mutations were previously used to improve CFP/YFP-based FRET biosensors but with a comparatively moderate success^42^. Semi-synthetic FRET biosensors such as Snifits also reach large dynamic ranges but require engineering of chemicals whose synthesis might not be accessible to non-chemists^12, 28, 43^. In contrast, ChemoX biosensors only require standard HaloTag fluorophore substrates, broadly used by the microscopy community^19, 44-49^. Current ChemoX-based sensors involve sensing domains with large conformational changes and future efforts will focus on developing sensors able to translate smaller conformational changes into similar dynamic ranges. In light of recent progresses in engineering new analyte binding domains^4, 5^, we foresee that ChemoX could support their conversion into potent biosensors for new biological activities.

Another key feature of the ChemoX resides in its versatility. First, ChemoX offers an extensive spectral tunability by exchanging the FRET FP donor and using different HaloTag fluorescent substrates as FRET acceptor. The color of the FRET sensors can thus be readily tuned, offering fluorescence excitation and emission maxima ranging from 386 nm to 558 nm and 448 to 668 nm, respectively. For example, the fluorescence maximum emission peaks of ChemoB_SiR_ sensors are separated by more than 200 nm (448 nm for EBFP2, 668 nm for SiR) and still offer dynamic ranges of up to 12.7 fold. Such sensors can theoretically be spectrally combined with GFP-, YFP- or even RFP-based intensiometric biosensors. We demonstrated this multiplexing ability by monitoring for the first time free NAD^+^ in two different cell compartments simultaneously, combining the two sensors ChemoB_SiR_-NAD and ChemoG_SiR_-NAD in fluorescence microscopy. This enabled to track the real-time co-regulation of subcellular free NAD^+^ pools in live cells, revealing a heterogeneous mitochondrial NAD^+^ regulation upon genotoxic stress that could not have been observed in previous studies based on lysates of subcellular fractionation^50, 51^. In the future, the multiplexing of biosensors will enable to further study this heterogeneity at the single cell level. Since the regulation of NAD^+^ inside and between different subcellular compartments plays important roles for key biological processes^25-27, 29^, the ChemoX-NAD biosensor palette represents a promising toolbox for the investigation of metabolic and signaling pathways.

Additionally, ChemoX FRET biosensors can be converted into single channel intensiometric and fluorescence lifetime biosensors as exemplified for ChemoD-NAD that showed ^max^ΔFI/FI_0_ of 227 % (with JF_635_) and ^max^Δτ of 1.18 ns (with CPY), respectively. ChemoD-NAD showed superior fluorescence intensity changes than some established intensiometric NAD^+^ biosensors^29^. Other intensiometric NAD^+^ biosensors^30^ or biosensors for different biological activities^18, 19, 52^ show superior dynamic ranges. In the future, it will be therefore interesting to explore whether ChemoD-based biosensors can be improved to reach comparable dynamic ranges. On the other hand, ChemoD-NAD showed fluorescence lifetime changes of at least similar magnitude than current state of the art fluorescence lifetime biosensors^53-55^. Based on synthetic far-red fluorophores (maximum emission wavelengths ≥ 628 nm), the intensiometric and FLIM-based modalities of ChemoD biosensors bring multiple advantages in term of brightness, photostability, phototoxicity and auto-fluorescence compared to FP-based biosensors. Future applications for deep tissue imaging of biological activities can thus be foreseen as HaloTag can be labeled in various animal models^49, 56^. Furthermore, extending the ChemoD biosensor design to additional, orthogonal SLPs should enable multiplexing of such far-red biosensors. Finally, using a BRET-FRET approach, luminescent ChemoL biosensors with dynamic ranges of up to 9.1 fold were developed, offering applications in high-throughput screening. The dynamic ranges of ChemoL biosensors revealed to be comparable^57^ or even superior^58, 59^ to recently developed BRET sensors.

In conclusion, ChemoX is a chemogenetic platform of FRET pairs enabling the development of biosensors with high dynamic ranges attributed to the reversible interaction of FPs with a fluorescently labeled HaloTag. Their color can be readily tuned by either changing the FP or the rhodamine fluorophore substrate, offering multiple options throughout the visible spectrum with the ability for multiplexing. As we demonstrated, the conversion of ChemoX FRET biosensors into intensiometric, fluorescence lifetime-based and bioluminescent biosensors is possible with little effort and through small modifications. ChemoX represents thus an extremely versatile platform enabling the rapid engineering of potent biosensors that should find broad applications in cell biology.

## Methods

Experimental procedures are described in supplementary information.

## Data availability

The X-ray crystal structures of ChemoG1_TMR_, ChemoG5_TMR_ and HaloTag7_Cy3_ were deposited to the PDB with accession codes 8B6S, 8B6T and 8B6R, respectively. Plasmids of interest from the study will be deposited at Addgene. Data is available by inquiry to the corresponding author upon reasonable request.

## Acknowledgements

This work was supported by the Max Planck Society, the Ecole Polytechnique Federale de Lausanne (EPFL) and the Deutsche Forschungsgemeinschaft (DFG, German Research Foundation) TRR 186. L.H. & M-C.H. were supported by the Heidelberg Biosciences International Graduate School (HBIGS). M-C.H was supported by the Boehringer Ingelheim found. The authors thank Ilme Schlichting for X-ray data collection. Diffraction data were collected at the Swiss Light Source, beamline X10SA, of the Paul Scherrer Institute, Villigen, Switzerland. The authors thank Luke Lavis for providing Janelia Fluor HaloTag substrates and Alexey N. Butkevich for providing CP580-CA, A. Herold (rAAVs), J. Kress (cell lines), B. Réssy (chemicals) and D. Schmidt (chemicals) for provision of reagents or material, and the members of the Chemical Biology Department (MPI, Heidelberg) for critical proofreading of the manuscript.

## Author contributions

L.H., L.B., A.B. and J.H. performed the in vitro experiments. L.H., A.E., M.H. and M.S.F. performed cell and microscopy experiments. L.H. processed and analyzed the data. M.T. solved the crystal structures. L.H. and J.H. analyzed the crystal structures. B.K. supervised and contributed to stable cell lines generation. J.H. and K.J. supervised the work. L.H. and J.H. wrote the manuscript with input from all authors.

