## Supplementary information for "A general method for the development of multicolor biosensors with large dynamic ranges"

### Methods

#### Reagents, chemicals and fluorophores

Reagents and chemicals were obtained from different manufacturers listed in **Methods Table M1.** Fluorophore chloroalkane (CA) substrates were synthesized according to literature by B. Réssy and D. Schmidt (MPI-MR, Heidelberg, Germany), purchased from commercial vendors (Promega) or kindly provided by Dr. Luke Lavis (HHMI, Ashburn, VA, USA) or Dr. Alexey N. Butkevich (MPIMR, Heidelberg, Germany). See **Methods Table M2** for details. The fluorophore substrates were prepared as 1 mM DMSO stocks, stored at -20 °C and used for all experiments. Milli-Q® water was used for all buffers and solutions.

#### Plasmids and cloning

Primers for cloning were obtained from Sigma-Aldrich, synthetic genes from Eurofins and PCRs were performed using the KOD Hot Start Master Mix (Sigma-Aldrich, Merck) according to the manufacturer's protocol. DNA size of PCR products was verified using standard agarose gel electrophoresis. Gibson assembly[1] was used as standard method for cloning. Transformations were performed using standard electroporation. Site-directed mutagenesis was performed using the Q5 Site-Directed Mutagenesis Kit (NEB) according to the manufacturer's protocol (transformation via heat-shock, *Escherichica coli* (*E. coli*) strain NEB 5-alpha) together with the online tool for primer design (NEBaseChanger, https://nebasechanger.neb.com/). All plasmids were amplified in the *E. coli* strain *E. cloni* 10G (Novagen), grown at 37 °C, and extracted using the QIAprep Spin Miniprep Kit (Qiagen) according to the manufacturer's protocol with the exception of the pAAV plasmids (see below). DNA sequences were validated by Sanger sequencing (Eurofins) and DNA was stored at -20 °C until further use. Plasmids are listed in **Methods Table M3** together with accession codes of plasmids deposited to Addgene and plasmids from Addgene used as template.

The pET-51b(+) vector (Novagen) was used as backbone for protein expression in *E. coli* (see below). Genes of interest (GOI) were flanked by an N-terminal Strep-tag®II and enterokinase cleavage sequence and a C-terminal poly-histidine tag (6x His) for protein purification (for details see protein sequences in Supplementary Methods below). For protein crystallization purposes, genes were flanked only by a N-terminal poly-histidine tag (10x His) followed by a tobacco etch virus (TEV) cleavage sequence.

The pCDNA5/FRT or pCDNA5/FRT/TO vectors (ThermoScientific) were used as backbone for protein expression in mammalian cells. For targeted expression at subcellular localizations, the GOI was flanked by N- and/or C-terminal localization sequences (for details see protein sequences in Supplementary Methods below). Localization sequences were amplified from Addgene plasmids as indicated:

- cytosol, nuclear export signal (NES, Addgene #101061[2], kind gift from E. Schreiter),
- nucleus, nuclear localization signal (NLS, Addgene #113931[3]),
- mitochondria matrix, Cox8 repetitions (Addgene #113916[4]),
- outer plasma membrane, IgKchL-PDGFR$_{tm}$ (Addgene #182009[5]),
- inner nuclear membrane, LaminB1 (Addgene #55069, kind gift from M. Davidson).

The NES sequence for cytosolic expression of constructs using pCDNA5/FRT plasmids (LPPLERLTL) was created in the design of the overhangs of PCR primers used for the cloning. For co-expression of similar genes (multiplex experiments), codon-optimized sequences for human (*Homo sapiens,* HsOpt) or zebrafish (*Danio rerio*, ZfOtp) were synthesized (Eurofins) to limit the probability of recombination events between the genes. Co-translational expression was performed using the T2A self-cleaving peptide sequence[6]. The sensor YC 3.6[7] was reconstructed by molecular cloning from Yellow Cameleon-Nano140 (Addgene #51966[8], kind gift of Takeharu Nagai).

The pAAV2-hSyn vector (Addgene #101061[2], kind gift from E. Schreiter) was used as backbone for cloning and downstream production of recombinant adeno-associated virus (rAAV) particles. The GOI was flanked by an N-terminal nuclear export signal (NES) sequence (for details see protein sequences in Supplementary Methods below). pAAV2-hSyn plasmids were cloned and amplified in *E. coli* strain NEB stable (NEB) and grown at 30 °C in Erlenmeyer flasks.

**Protein expression and purification**

*E. coli* strain BL21 (DE3)-pLysS (Sigma-Aldrich) was used for the production of proteins. After electroporation with plasmid DNA, colonies grown overnight on Lysogen broth (LB) agar plates supplemented with 100 µg/mL Ampicillin (Amp) at 37 °C were picked to inoculate 5 mL liquid LB-Amp and grown over night (o.n.) at 37 °C and 220 rotations per minute (rpm). O.n. cultures were diluted 1:200 in 0.1-1 L LB-Amp and grown at 37 °C and 220 rpm until reaching an optical density at 600 nm ($OD_{600}$) of 0.6-0.8. Protein expression was induced by the addition of 0.5 mM isopropyl-β-D-thiogalactopyranoside (IPTG) and the cultures grown at 16 °C for 20-24 h. Cells were harvested by centrifugation at 4,500 g and 4 °C for 10 min. Cell pellets were resuspended in 30 mL ice-cold lysis buffer (50 mM $KH_2PO_4$, 300 mM NaCl, 5 mM imidazole, pH 8.0) supplemented with 1 mM phenylmethylsulfonyl fluoride (PMSF) and 250 µg/mL lysozyme. Cells were lysed by sonication (7 min with 50% on/off cycles and 70% amplitude, SONOPULS Bandelin) and cell debris cleared by centrifugation at 10,000 g and 4 °C for 15 min. The supernatant was incubated with 0.5-1 mL Ni-NTA resin (HisPur™ Ni-NTA Superflow Agarose, ThermoScientific) for 1 h at 4 °C on a roller shaker. The Ni-NTA beads were poured into a 5 mL polypropylene column and washed with 10 column volumes of wash buffer (50 mM $KH_2PO_4$, 300 mM NaCl, 10 mM imidazole, pH 7.5). His-tagged proteins were eluted with 1.5 mL elution buffer (50 mM $KH_2PO_4$, 300 mM NaCl, 500 mM imidazole, pH 7.5). Subsequently, the buffer was exchanged with Activity buffer (50 mM HEPES, 50 mM NaCl, pH 7.3) during protein concentration using Amicon Ultra Centrifugal Filters (Millipore) with an appropriate molecular weight cutoff.

Protein concentration was determined by absorbance measurements at 280 nm using a Nanodrop 2000c spectrophotometer. The extinction coefficient at 280 nm for the different proteins was extrapolated from the amino acid sequence using the Geneious software. Protein purity was confirmed by standard SDS-PAGE using 4–20% Mini-PROTEAN TGX Stain-free Precast Protein Gels (BioRad) that were imaged using a

GelDoc imager (BioRad). Purified proteins were stored in presence of 45% (w/v) glycerol at -20 °C until further use.

**Protein crystallization**

For protein crystallization, HaloTag7 and ChemoG variants were produced and purified as previously reported[9]. In short, after Ni-NTA resin purification, the buffer was exchanged to TEV-cleavage buffer (25 mM $Na_2HPO_4$, 200 mM NaCl, pH 8.0). TEV protease (weight ratio 30:1, protein of interest(POI):TEV) was added to protein samples and cleavage was performed at 30 °C overnight. After filtering the solution (0.22 µm), the cleaved protein was collected by reverse Ni-NTA resin purification (i.e. uncleaved proteins and His-tagged TEV remained bound to the resin while the cleaved POI was recovered from the flow-through) on a HisTrap FF crude column (Cytiva) using an ÄktaPure FPLC (Cytiva) collecting the flow through using the same buffer as for Ni-NTA resin purification (50 mM $KH_2PO_4$, 300 mM NaCl, 10 mM imidazole, pH 7.5). The proteins were further purified by size exclusion chromatography on a HiLoad 26/600 Superdex 75 pg column (Cytiva) together with exchanging the buffer to Activity buffer. The proteins were concentrated to 5 µM using Amicon® Ultra 4 mL Centrifugal Filters (Merck) and fully labeled with 10 µM fluorophore substrate (ChemoG variants with TMR-CA, HaloTag7 with Cy3-CA) for at least 4 h at room temperature (RT). The labeled proteins were then concentrated to ~250 µL reaching a final concentration between 10 – 16 mg/mL. Proteins were quantified using the absorbance at 280 nm and the extinction coefficient of the protein was corrected for the fluorophore absorbance at 280 nm using the respective correction factor (CF280nm) ($TMR_{CF280nm}$ = 0.16, 13,920 $M^{-1}$ $cm^{-1}$; $Cy3_{CF280nm}$ = 0.08, 12,000 $M^{-1}$ $cm^{-1}$).

Crystallization was performed at 20 °C using the vapor-diffusion method. Crystals of HaloTag7-Cy3 were grown by mixing equal volumes of protein solution and a reservoir solution containing 0.2 M magnesium acetate, 19 % (m/v) PEG 3350. Crystals of ChemoG1-TMR and ChemoG5-TMR were obtained by mixing equal volumes of protein solution and precipitant solution composed of 0.085 M Tris-HCl pH 8.5, 0.17 M sodium acetate, 15 % (v/v) glycerol, 27 % (m/v) PEG 4000 or 0.1 M Tris-HCl pH 8.5, 0.2 M magnesium chloride, 30 % (m/v) PEG 4000, respectively. Crystals of HaloTag7-Cy3 and ChemoG5-TMR were briefly washed in cryoprotectant solution consisting of the reservoir solution supplemented with 20 % (v/v) glycerol before flash-cooling in liquid nitrogen, whereas crystals of ChemoG1-TMR were flash-cooled directly in the mother liquor.

**X-ray diffraction data collection and structure determination**

Single crystal X-ray diffraction data were collected at 100 K on the X10SA beamline at the SLS (PSI, Villigen, Switzerland). All data were processed with XDS[10]. The structure of HaloTag7-Cy3 was determined by molecular replacement (MR) using Phaser[11] and HaloTag7-TMR coordinates (PDB ID: 6Y7A) as a search model. The structure of ChemoG1-TMR was determined using HaloTag7-TMR (PDB ID: 6Y7A) and GFP (PDB ID: 1GFL) coordinates, the ChemoG1-TMR model was used to determine the ChemoG5-TMR structure. Geometrical restraints for Cy3 and TMR ligands were generated using the Grade server[12]. The final models were optimized in iterative cycles of manual rebuilding using Coot[13] and refinement using

Refmac5[14] and phenix.refine[15]. Data collection and refinement statistics are summarized in **Methods Table M4**, model quality was validated with MolProbity[16] as implemented in PHENIX.

Atomic coordinates and structure factors have been deposited in the Protein Data Bank under accession codes: 8B6R (HaloTag7-Cy3), 8B6S (ChemoG1-TMR), 8B6T (ChemoG5-TMR).

**General considerations for fluorescence spectroscopy**

Fluorescence measurements were performed in 100 µL Activity buffer supplemented with 0.5 mg/mL BSA in black non-binding flat bottom 96 well plates (Perkin Elmer) unless stated differently. For FRET measurements, proteins were diluted to 200 nM in Activity buffer and HaloTag7-based proteins were additionally labeled with 400 nM fluorophore-CA substrates for 1 h at RT. For fluorescence intensity measurements of intensiometric biosensors, HaloTag7-based proteins were diluted to 1 µM and labeled with 200 nM CA-fluorophore substrates for 1 h at RT. Emission spectra of the proteins were acquired with a Multimode Spark 20M microplate reader (Tecan) using monochromators and at 37 °C. Pipetted plates were temperature-equilibrated (37 °C) inside the plate reader for 20 min before the measurement. Flash numbers, gain, excitation and emission wavelengths were adjusted depending on the fluorescent proteins and synthetic fluorophores (for detailed settings see **Methods Table M5**). Excitation and emission bandwidths were set to 20 nm and 10 nm, respectively, with an emission step size acquisition of 2 nm.

**Analyte titrations of biosensors**

Analyte solutions were prepared at 10x final concentration in Activity buffer and diluted to 1X final concentration in 100 µL Activity buffer (final volume) in presence of the labeled proteins in a 96 well plate. For titrations, dilution series of the analyte were prepared in Activity buffer. The detailed analyte concentrations are listed in **Methods Table M6**. Activity buffer without analyte was always included as control.

$NAD^+$ titrations in presence of structurally similar analytes were performed as previously explained but in presence of the structurally similar analyte (diluted from a 10x solution) and labeled proteins. Activity buffer without $NAD^+$ but with 1x final concentration of the structurally similar analyte was always included as control.

For calcium titrations, a Calcium Calibration Buffer Kit (Life technologies) was used to precisely control the free $Ca^{2+}$ concentration. Two buffers containing either 0.1 M CaEGTA or 0.1 M $K_2$EGTA were mixed in defined ratios according to the manufacturer's protocol to generate buffers with free $Ca^{2+}$ concentrations ranging from 10 nM to 39 µM. 0.1 M $K_2$EGTA (0 µM free $Ca^{2+}$) was always included as control. The free $Ca^{2+}$ concentrations were calculated using equation (**1**):

$$[Ca^{2+}]_{free} = K_{\mathrm{d}}^{\mathrm{EGTA}} \times \frac{[CaEGTA]}{[K_2EGTA]} \quad (\mathbf{1})$$

where $K_d^{EGTA}$ is the dissociation constant of EGTA for $Ca^{2+}$ in 0.1 M KCl at a given pH and temperature[17], and [CaEGTA] divided by [$K_2$EGTA] is the molar ratio of the two buffers. For the calculation of free $Ca^{2+}$, the $K_d^{EGTA}$ at pH 7.2 and 37 °C is 107.9 nM.

For the titration of calcium sensors using the calcium buffers, HaloTag7-based proteins were diluted to 2 µM and labeled with 4 µM fluorophore-CA substrate for 1 h at RT. The calcium sensors were diluted to a final concentration of 200 nM in 20 µL of calcium buffer in a black low volume flat bottom 384 well plate (Corning).

**Sensitivity assays**

The pH sensitivity of the constructs was evaluated using two sodium phosphate-based buffers (SPG pH 4.0 & 10.0, both 1 M, Jena Bioscience) mixed in defined ratios according to the manufacturer's protocol to yield buffers with different pH ranging from 5.5 to 8.0. The buffers were diluted 10-fold in water (0.1 M final concentration), supplemented with 0.5 mg/mL BSA and 50 mM NaCl. The salt concentration sensitivity of the constructs was evaluated using 50 mM HEPES pH 7.3 buffer supplemented with 0.5 mg/mL BSA and various NaCl concentrations ranging from 50 to 500 mM NaCl. A condition without salt (0mM NaCl) was included as well.

For the sensitivity assays, the proteins were diluted to 2 µM and labeled with 4 µM CA-fluorophore substrate for 1 h at RT. The labeled proteins were then diluted to 200 nM in the different buffers. For the biosensors, 10x analyte solutions were added and diluted to 1x final concentration. Measurements were conducted as previously explained.

**Calculation of the FRET efficiency**

The Förster resonance energy transfer (FRET) efficiency (E) was determined by the fraction of quenched donor fluorescence intensity (FI) using equation (**2**):

$$E = 1 - \frac{FI_{DA}}{FI_{D}} \quad (\mathbf{2})$$

where $FI_{DA}$ and $FI_{D}$ are the maximum FI values of the FRET donor with or without FRET acceptor, respectively.

The Förster radius $R_0$ was calculated using equation (**3**):

$$R_0 = 0.211\sqrt[6]{\kappa^2 \times n^{-4} \times Q_D \times J(\lambda)} \quad (\mathbf{3})$$

where $\kappa^2$ is the orientation factor (set to 0.667), n is the refractive index (set to 1.33), $Q_D$ is the quantum yield of the donor (set to the value according to https://www.fpbase.org/[18]) and J(λ) is the spectral overlap of the donor emission and acceptor excitation spectra which was calculated using equation (**4**):

$$J(\lambda) = \frac{\int_0^\infty FI_D(\lambda) \times \varepsilon_A(\lambda) \times \lambda^4 d\lambda}{\int_0^\infty F_D(\lambda) d\lambda} \quad (\mathbf{4})$$

where $FI_D$ is the donor fluorescence at the wavelength λ and $\varepsilon_A(\lambda)$ is the extinction coefficient of the acceptor at the wavelength λ. Spectral overlaps were determined using the software a|e Fluortools (http://www.fluortools.com/software/ae).

**Fluorescent biosensor characterization**

From acquired fluorescence emission spectra, the maximum FI values of the fluorescent protein and/or fluorophore were extracted at their maximum emission wavelength, defined in **Methods Table M5**. For FRET measurements, the FRET/FP ratios (R) were calculated by dividing the maximum FI of the FRET acceptor by the maximum FI of the FRET donor. The dose-dependent response of the FRET biosensors was determined by plotting the ratio change ($\Delta R/R_0$) over the concentration of the cognate analyte using equation (**5**):

$$\Delta R/R_0 = \frac{R_i}{R_0} - 1 \quad (\mathbf{5})$$

where $R_i$ is the FRET/FP ratio at a given analyte concentration i and $R_0$ is the FRET/FP ratio in absence of analyte. The maximum ratio change ($^{max}\Delta R/R_0$) is calculated with $R_i$ at saturating concentration of analyte.

For fluorescence intensity measurements, the dose-dependent response of the intensiometric sensors was determined by plotting the FI change ($\Delta FI/FI_0$) over the concentration of the cognate analyte using equation (**6**):

$$\Delta F/F_0 = \frac{FI_i}{FI_0} - 1 \quad (\mathbf{6})$$

where $FI_i$ is the FI at a given analyte concentration i and $FI_0$ is the FI in absence of the analyte. The maximum FI change ($^{max}\Delta FI/FI_0$) is calculated with $FI_i$ at saturating concentration of analyte.

For fluorescence lifetime (τ) measurements, experiments were conducted on a confocal microscope (Leica SP8 FALCON, Leica Microsystems). For details about the determination of τ values, see dedicated microscopy section thereafter. The dose-dependent response of the τ sensors was determined by plotting the τ change (Δτ) over the concentration of the cognate analyte using equation (**7**):

$$\Delta\tau = \tau_i - \tau_0 \quad (\mathbf{7})$$

where $\tau_i$ is the τ at a given analyte concentration i and $\tau_0$ is the τ in absence of the analyte. The maximum τ change ($^{max}\Delta\tau$) is calculated with $\tau_i$ at saturating concentration of analyte.

**Data processing and fitting**

Data were mathematically processed as explained using the Excel software (Microsoft). GraphPad Prism (version 8.1.0) was used to fit a sigmoidal dose response to the titration data (FRET, intensiometric, fluorescence lifetime and BRET-FRET) using equation (**8**):

$$Y = \text{Bottom} + \frac{x^H * (Top - Bottom)}{x^H + C50^H} \quad \textbf{(8)}$$

where Y is the response ($\Delta R/R_0$, $\Delta FI/FI_0$ or $\Delta\tau$), Bottom and Top are the lower and upper plateau of the response, respectively, x is the analyte concentration, H is the Hill coefficient (i.e. slope factor) and C50 is the analyte concentration at which the response is half-maximal.

For some data representation, the ratio R was chosen instead of $\Delta R/R_0$, the FI was chosen instead of $\Delta FI/FI_0$ and $\tau$ was chosen instead of $\Delta\tau$. Data and fit were represented using the GraphPad Prism software (version 8.1.0).

**Bioluminescence spectroscopy**

Bioluminescence measurements were performed in 100 µL Activity buffer supplemented with 0.5 mg/mL BSA in white non-binding flat bottom 96 well plates (Perkin Elmer), except for measurements of calcium sensors, which were performed in 20 µL calcium buffers (prepared as described above in section Analyte titrations of biosensors) in white low volume non-binding flat bottom 384 well plates (Corning). Proteins were diluted to 200 nM and labeled with 400 nM CA-fluorophore substrate for 1 h at RT. Labeled proteins were further diluted to 0.5 nM final concentration together with the 1x final concentration of analyte (as described above in section Analyte titrations of biosensors) and 1:1000 diluted Nano-Glo Luciferase Assay Substrate (Promega). Pipetted plates were incubated for 20 min at 37 °C before the measurement to equilibrate the temperature. Bioluminescence emission spectra were acquired with a Multimode Spark 20M microplate reader (Tecan) using the integrated luminescence module at 37 °C. Emission spectra were acquired from 398-653 nm with a step size of 15 nm and an integration time of 200 ms.

Luminescent sensors are constructed such that a BRET phenomenon occurs between NanoLuc (maximum emission wavelength observed ($^{max}_{em}\lambda$) = 460 nm) and EGFP ($^{max}_{em}\lambda$ = 518 nm), and a BRET-FRET phenomenon between EGFP and the fluorophore CPY ($^{max}_{em}\lambda$ = 638 nm). The BRET-FRET/EGFP ratios (R) were calculated by dividing the maximum luminescence intensity (LI) value of the BRET-FRET acceptor CPY by the maximum LI value of the BRET-FRET donor EGFP. The dose-dependent response of the luminescent biosensors was determined by plotting the ratio change ($\Delta R/R_0$) over the concentration of the cognate analyte following the equation (**5**). The maximum ratio change ($^{max}\Delta R/R_0$) is calculated with the ratio at saturating concentration of analyte. A sigmoidal dose response was fitted to the titration data using the equation (**8**), from which the C50 of the sensors were extrapolated.

**Cell culture**

HeLa Kyoto (RRID:CVCL_1922[19]) and U-2 OS Flp-In T-REx cells[20] were cultured in high glucose (4.5 g/L) DMEM + GlutaMAX™ medium (Gibco) supplemented with 10 % heat-inactivated FBS (Gibco). Cells were cultured at 37 °C and 5 % $CO_2$ in a humidified cell culture incubator. Cells were handled under a sterile laminar flow hood and kept in culture for a maximum of 4 weeks splitting them every 2-4 days or at confluency. Contamination with Mycoplasma was regularly checked by PCR. No contamination was detected in the course of this study.

**Generation of stable cell lines**

Stable cell lines were generated using the Flp-In T-REx system[20]. U-2 OS Flp-In T-REx cells were grown to 80% confluency in a T-25 cell culture dish and co-transfected with a pCDNA5/FRT or pCDNA5/FRT/TO plasmid encoding the gene of interest (GOI) and the plasmid pOG44 (Invitrogen) in a 1:10 ratio (total 4 µg DNA) using Lipofectamine 3000 Transfection Reagent (Invitrogen) according to the manufacturer protocol. 14-16 h post-transfection, the transfection mix was exchanged with fresh cell culture medium supplemented with 100 µg/mL Hygromycin B (ThermoScientific) to select cells that stably integrated the plasmid into the genome. After 48 h of selection, cells were recovered in fresh cell culture medium until confluency. Cells were sorted in bulk (total of 100,000 cells) for moderate expression levels of the GOI by fluorescence activated cell sorting (FACS) using a FACSMelody Cell sorter (BD Biosciences). Cells were sorted for EGFP fluorescence (Blue laser, 488 nm with 530/30 BP filter) or, in case EGFP was not encoded by the GOI, the cells were labeled with SiR-halo (= SiR-CA) and sorted for SiR fluorescence (Red laser, 640 nm with 660/10 BP filter). For cells that integrated a pCDNA5/FRT/TO plasmid, the protein expression was induced by 200 ng/mL doxycycline for 24h before sorting or labeling with SiR-halo. Stable cell lines generated in this study are listed in **Methods Table M3**.

**Transient transfection of mammalian cell lines**

For transient transfections, 30,000 or 200,000 cells were seeded into a black 96 or 24 well imaging plate with glass bottom (Cellvis), respectively, and reverse transfected with pCDNA5/FRT or pCDNA5/FRT/TO plasmid DNA using Lipofectamine 3000 Transfection Reagent (Invitrogen) according to the manufacturer's protocol. Cells were incubated with the transfection mix for 8-12 h before the medium was exchanged with fresh cell culture medium.

**Labeling of mammalian cell lines**

For transiently transfected cells, cells were incubated for 12 h in cell culture medium supplemented with 500 nM fluorophore-CA substrate 24 h post-transfection to achieve labeling of the constructs. For stable cell lines, 24 h prior to cell labeling, 10,000 or 50,000 cells were seeded into a black 96 or 24 well imaging plate with glass bottom (Cellvis), respectively. For stable cell lines that integrated a pCDNA5/FRT/TO plasmid, 200 ng/mL doxycycline was added to the medium for 24h to induce the protein expression. Cells were then labeled as explained for transiently transfected cells. Cells that express GOIs not encoding HaloTag7, were incubated in normal cell culture medium. Excess of fluorophore-CA substrate was removed after labeling by washing the cells three times for 5, 15 and then 30 min in phenol red-free cell culture medium (Gibco) prior to imaging.

Staining of the nucleus and mitochondria with Hoechst 33342 (Invitrogen) and Mitotracker Red FM (Invitrogen), respectively, was performed according to the manufacturer's protocol. For co-localization experiments, cells expressing biosensors were not labeled with fluorophore-CA substrates to avoid potential spectral crosstalk with MitoTracker RedFM. Images were acquired on a confocal microscope (as described below). Images of Hoechst 33342-stained cells were acquired at 355 nm excitation wavelength and 400-450 nm emission wavelengths. Images of Miotracker Red FM-stained cells were acquired at 600 nm excitation wavelength and 620-670 nm emission wavelengths.

**Preparation of neuron cultures**

Prior to the preparation, black 24 well imaging plates with glass bottom were coated with 100 µg/mL poly-L-ornithine (diluted in water) for 20 min at RT, then washed 2x with 1x PBS pH 7.4 and subsequently coated with 1 µg/mL laminin (dissolved in 1x HBSS) for 1 h at RT. Hippocampi were isolated from sacrificed new born rat pups (0-1 day, WISTAR rats) as described previously.[21] In brief, the brain was extracted by dissecting the skull-cap in a posterior-anterior direction. The hippocampi were removed and placed into ice-cold 1x HBSS in presence of 0.25 % trypsin (final concentration) and incubated for 20 min at 37 °C. Tryptic digestion was quenched by addition of DMEM (Gibco) containing 10 % heat-inactivated FBS. The neurons were centrifuged at 200 g for 5 min at RT, washed 3x with 1x HBSS and then resuspended in 5 mL phenol red-free Neurobasal medium (NB, Gibco). Neurons were mechanically separated using a pipette until obtaining a homogeneous solution. The solution was filtered through a cell strainer (40 µm pore diameter) and live cell numbers were determined using the Countess® II FL Automated Cell Counter (ThermoScientific). 55,000 cells were seeded per well of a pre-coated 24 well imaging plate in 1 mL NB medium. 2 h after seeding, medium was exchanged with fresh NB medium. Neurons were kept in a humidified cell culture incubator at 37 °C and 5 % $CO_2$ and handled under a sterile laminar flow hood. All reagents were sterile filtered with a 0.2 µm filter. 1x HBSS used during the preparation was supplemented with 1x Penicillin/Streptomycin (Pen/Strep, Gibco) and 200 µM kynurenic acid. NB medium used during the preparation and for culturing of the neurons was supplemented with 1x Pen/Strep, GlutaMAX™ and B27.

**Generation of recombinant AAVs**

Recombinant AAVs rAAVs were obtained using pAAV2-hSyn plasmids that were extracted from *E. coli* NEB stable bacteria using the GeneJET Endo-Free Plasmid-Maxiprep-Kit (ThermoFisher) according to the manufacturer's protocol and verifying proper open reading frame (ORF) and inverted terminal repeat (ITR) sequences by Sanger sequencing (Eurofins). rAAVs were generated as described previously[22]. In brief, plasmids pRV1 (AAV2 Rep and Cap sequences), pH21 (AAV1 Rep and Cap sequences), pFD6 (Adenovirus helper plasmid) and the AAV plasmid containing the recombinant expression cassette flanked by AAV2 packaging signals (ITRs) were transfected via Polyethylenimine 25,000 (PEI25,000, Sigma-Aldrich) into HEK293 cells (ACC305, DSMZ[23]). 5 days post transfection, the medium and cells were harvested by centrifugation at 1,000 g for 5 min at 4 °C. The cells were lysed using TNT extraction buffer (20 mM Tris pH 7.5, 150 mM NaCl, 1 % Triton-X-100, 10 mM $MgCl_2$). The cell debris were spun down at 3,000 g for 5 min at 4 °C. The supernatant was treated with 50 U/mL Benzonase (final concentration, Sigma-Aldrich) for 30-60 min at 37 °C (samples were inverted every 20 min to mix the content). The rAAVs were purified from supernatant via Äkta-Quick FPLC (Cytiva) using AVB Sepharose HiTrap columns

(Cytiva). The columns were equilibrated with PBS pH 7.4 and the virus particles were eluted with 50 mM glycine-HCl pH 2.7. The purified virus particles were concentrated and the buffer was exchanged to PBS pH 7.3 using Amicon Ultra Centrifugal Filters (Millipore) with a MWCO of 100 kDa. rAAVs were aliquoted in 10 µL, flash frozen and stored at -80 °C until further use. The precise rAAV titer for AAV2/1-hSyn1-NES-ChemoG-CaM was 1 x $10^{13}$ particles/mL as evaluated by qPCR (genome copies) as described previously[24].

**AAV-transduction and labeling of rat hippocampal neurons**

Cultured rat hippocampal neurons were transduced with rAAVs after 8 days *in vitro* (DIV). rAAVs (~ 5 x $10^9$ particles, 0.5 µL) were diluted in 50 µL of phenol red-free NB medium and added to the medium (1 mL) of the cultured neurons. After 12 DIV, the neurons were labeled with 200 nM fluorophore-CA substrate (final concentration) by adding 100 µL of phenol red-free NB medium supplemented with 2 µM fluorophore-CA substrate to the ~ 1 mL of NB medium in which the AAV-transduced neurons were cultured. After 12 h labeling, the neurons were used for live cell imaging on a widefield microscope (for experimental details see section Widefield microscopy below) without washing out the excess of fluorophore-CA substrate.

**Confocal microscopy**

Confocal microscopy experiments were performed on a laser-scanning confocal microscope (Leica SP8, Leica Microsystems) equipped with a Leica TCS SP8 X scanhead, a HC PL APO 40x/1.10 water motCORR CS2 objective, a SuperK white light laser, a 405 nm diode laser and hybrid photodetectors for single molecule detection (HyD SMD). The microscope was maintained in an environmental chamber with temperature control set to 37 °C, $CO_2$ control set to 5 % and humidity control set to 68%. The imaging plate was placed on the confocal microscope stage and temperature-equilibrated for 30 min. Confocal images were recorded in 512x512 pixel resolution (12- or 16-bit), with a scan speed of 400 Hz, a pixel dwell time of 3.16 µs, pinhole size of 1 Airy unit and laser pulse rate of 80 MHz, unless stated differently. A sequential scan mode (between frames) was used for imaging with multiple excitation wavelengths. Z-stacks were acquired where necessary with a step size of 1 µm. For further details, notably excitation and emission settings, see **Methods Table M7**.

**Widefield microscopy**

Widefield microscopy experiments were performed on a DMi8 widefield microscope (Leica, Leica Microsystems) equipped with a HC PL APO 20x/0.80 dry objective and an external filter wheel (Leica Microsystems). The microscope was maintained in an environmental chamber with a temperature of 37 °C and a $CO_2$ concentration of 5 %. The imaging plate was placed on the widefield microscope stage and temperature-equilibrated for 30 min. Widefield images of mammalian cells were recorded in 512x512 pixel resolution (16-bit), with an exposure time of 500 ms, 4x4 binning and a cycle time of 766 ms. Widefield images of rat hippocampal neurons were recorded in 256x256 pixel resolution (16-bit), with an exposure time of 50 ms, 8x8 binning and a cycle time of 81 ms. Z-stacks were acquired where necessary with a step size of 1 µm. EGFP fluorescence was acquired using a 470 nm LED together with a 474/24 nm bandpass filter for excitation and a 525/50 nm bandpass filter for emission detection. FRET fluorescence was acquired using a 470 nm LED together with a 474/24 nm bandpass filter for excitation and a 700/75 nm bandpass filter for emission detection (for further details see **Methods Table M7**).

**Fluorescence lifetime imaging microscopy**

Fluorescence lifetime imaging microscopy (FLIM) experiments were performed on a confocal microscope (Leica SP8, Leica Microsystems; as described above) containing the FALCON system (Leica). The microscope was maintained in an environmental chamber with a temperature control set to 37 °C, $CO_2$ control set to of 5 % and humidity control set to 68%.

For *in vitro* titrations of τ biosensors, proteins were diluted to 2 μM in Activity buffer supplemented with 0.5 mg/mL BSA and labeled with 500 nM fluorophore-CA substrate for 1 h at RT. Labeled sensors were mixed with different analyte concentrations (as described in section Analyte titrations of biosensors) inside a black 96 well imaging plate with glass bottom (Cellvis). The pipetted plates were placed on the confocal microscope stage and temperature-equilibrated at 37 °C for 30 min. The confocal volume was focused to the maximum fluorescence intensity of the labeled fluorophore and images were taken and processed as for FLIM cell images (explained thereafter). Fluorescence lifetime values were used to characterize the sensor behavior in terms of maximum response ($^{max}\Delta\tau$) and C50 as explained in the dedicated section above.

For live cell experiments, stable cell lines expressing the τ biosensors were seeded and labeled with fluorophore substrates in black 24 well imaging plates with glass bottom (as described in the section Labeling of mammalian cells). Imaging plates were placed on the confocal microscope stage and temperature-equilibrated for 30 min. The motorized correction ring of the objective was adjusted for each imaging plate (*in vitro* titrations and live cell experiments). Excitation and emission detection wavelengths were setup depending on the fluorophore (for details see **Methods Table M7**). FLIM images were recorded in 512x512 pixel resolution (8-bit), pinhole size of 1 Airy unit, a scan speed of 400 Hz, 12 line repetitions and 1 frame repetition, and laser pulse rate of 40 MHz. The laser power was adjusted to obtain less than 1 photon arrival per laser pulse. The acquired images were intensity-thresholded to remove unspecific background signal. Average fluorescence lifetimes were determined in the LAS X Software (Leica Microsystems) using n-exponential reconvolution and globally fitting mono- (ChemoG-NAD and HaloTag7) or triexponential (ChemoD-NAD) decay models to the decay data. For time course experiments in cells, regions of interest (ROIs) were manually defined for each cell. Cells with saturated fluorescence intensity (FI) values or $X^2$>1.2 were excluded from the analysis. The calculated average intensity-weighted fluorescence lifetimes are represented. FastFLIM images were used to make figure panels.

**Live cell imaging and drug treatment**

For time course experiments, transiently or stably expressing cells were seeded in 24 well imaging plates (Cellvis) and labeled as previously described. Media were exchanged 1 h prior to the start of the time course as following:

- HeLa Kyoto cells - ATP sensors - phenol red-free cell culture medium without glucose (Gibco),

- U-2 OS cells - NAD+ sensors - phenol red-free 1x HBSS with calcium and magnesium (Corning),
- HeLa Kyoto cells - calcium sensors - phenol red-free cell culture medium (Gibco).

Drug treatments were applied directly on the fluorescence microscope during the imaging acquisition (for details about imaging acquisition see the corresponding microscope sections above). Stock solutions of drugs were freshly prepared in the same medium/buffer in which the cells are incubated and pre-warmed inside the microscope chamber 30 min before the start of the measurement:

- 2x solution of *N*-methyl-*N*'-nitro-*N*-nitrosoguanidine (MNNG, 200 µM),
- 2x solution of 2-deoxy-D-glucose (2DG, 20 mM),
- 5x solution of glucose (100 mM) and
- 2x solution of histamine (20 µM).

The 2x solutions were added in a 1:1 ratio (1 mL:1 mL) and the 5x solution in a 1:4 ratio (0.5 mL:2 mL) to the wells during the time course using a pipette.

For endpoint measurements of NAD+ biosensors, cells were seeded in 96 well imaging plates (Cellvis) and labeled as previously described. After labeling/washing, the cells were incubated in phenol red-free cell culture medium supplemented with 0.01 % DMSO (v/v, control), 100 nM FK866 (prepared from 1 mM DMSO stock solution), and/or 1 mM nicotinamide riboside (NR, prepared from 1 M water stock solution) for for 24 h. Cells were kept in a humidified cell culture incubator at 37 °C and 5 % $CO_2$ between the preparation steps. After the treatment, the cells were directly used for imaging by fluorescence confocal microscopy.

**Electric field stimulation of neurons**

Time course experiments of rat hippocampal neurons expressing calcium sensors were performed in phenol red-free NB medium supplemented with Pen/Strep, GlutaMAX™ and B27. Rat hippocampal neurons were seeded in black 24 well imaging plates (as described above) and imaged on a DMi8 widefield microscope (as described above). 30 min prior to the experiment, 100 µL phenol-red free NB medium supplemented with a synaptic blocker cocktail (25 µM (DL)-2-amino-5-phosphonovaleric acid (APV, SantaCruz) and 10 µM 2,3-dihydroxy-6-nitro-7-sulphamoyl-benzo(F)quinoxaline (NBQX, Sigma; final concentrations) was added to the neurons. Neurons were placed on the widefield microscope stage and temperature-equilibrated for 30 min. A custom-built 24 well cap stimulator with platin electrodes inserted into the medium was mounted on top of the imaging plate linked to a stimulation control unit as previously described[25]. Trains of action potentials (APs) from 1-200 APs were evoked by field stimulation at 80 Hz, 100 mA and 1 ms pulse width in 10 s intervals.

**Bioluminescence measurements in mammalian cells**

Cells transiently or stably expressing luminescent biosensors were seeded in white cell culture-treated 96 well plates with transparent bottom (BrandTech) and labeled as descried above. For endpoint measurements of NAD+ biosensors, cells were treated as for fluorescent microscopy experiments (as described above), then washed 2x with sterile filtered 1x PBS pH 7.4 (Gibco) and then incubated in 100 µL phenol red-free cell culture medium supplemented with 1:500 diluted cell-permeable NanoBRET Nano-Glo substrate (Promega) and 1:1000 diluted cell-impermeable NanoLuc inhibitor (Promega). The cells were then incubated at 37 °C for 30 min before the measurement. Bioluminescence spectra were acquired as described for *in vitro* characterization.

For time course measurements, cells were washed 2x with sterile filtered 1x PBS pH 7.4 (Gibco) and then incubated in 100 µL of

- phenol red-free cell culture medium ($Ca^{2+}$ sensor),
- phenol red-free cell culture medium without glucose (ATP sensor) or
- sterile filtered phenol red-free 1x HBSS with calcium and magnesium (NAD+ sensor).

All media were supplemented with 1:250 diluted cell permeable NanoBRET Nano-Glo substrate (Promega) and 1:1000 diluted cell-impermeable NanoLuc inhibitor (Promega). Cells were then incubated inside a Multimode Spark 20M microplate reader (Tecan) at 37 °C for 30 min before the start of the measurement to equilibrate the temperature. Maximum emission peaks of the BRET-FRET acceptor CPY (625-650 nm) and BRET-FRET donor EGFP (505-530 nm) were recorded during the time course to trace the BRET-FRET/EGFP ratio over time using the same parameter settings as for the acquisition of emission spectra *in vitro*. Drug treatments were applied directly on the plate reader. Stock solutions were freshly prepared in the same medium/buffer in which the cells are incubated and pre-warmed at 37 °C 30 min before the start of the measurement:

- 2x solution for *N*-methyl-*N*'-nitro-*N*-nitrosoguanidine (MNNG, 200 µM),
- 2x solution for 2-deoxy-D-glucose (2DG, 20 mM),
- 5x solution for glucose (100 mM),
- 2x solution for histamine (20 µM) and
- 2x solution for ionomycin (2 µM).

For addition of the reagents, the time course was paused, the 96 well plate was ejected and solutions were added in a 1:1 ratio (100 µL:100 µL, 2x solutions) or 1:4 ratio (50 µL:200 µL, 5x solution) to the wells using a multichannel pipette. The time course was immediately resumed after addition. Bioluminescence emission spectra were recorded for each well at the end of the time course (as described for *in vitro* characterization) to verify that the sensor still provide an emission spectra in which each channel strongly provide photons at the end of the treatment.

**Image analysis**

All images were analyzed using the software Fiji[26]. Z-planes of images (12- or 16-bit) were combined into a composite image by maximum intensity Z-projection. Background signal was subtracted with a rolling ball radius of 50 pixels. Brightness and contrast were adjusted identically for each channel of a processed image unless stated differently. The processed images were further used for quantitative analysis and figure panels.

For the generation of FRET/FP ratio images, the image of the FRET acceptor channel was duplicated and thresholded (Otsu's method). A binary mask was created from the thresholded image and applied to the

FRET acceptor and FRET donor channels that were subsequently divided (FRET acceptor channel/FRET donor channel) and converted into a 32-bit format, all using the Image calculator function.

For quantitative analysis, regions of interest (ROIs) were manually defined for each cell. The FI values of ROIs were measured in the respective channel of background-subtracted images. Cells with saturated FI values were excluded from the analysis. For FRET experiments, the FRET/FP ratios were determined by dividing the measured FI of the FRET acceptor by the measured FI of the FRET donor channel of each ROI. Data were mathematically processed using the Excel software (Microsoft). Data were represented using the GraphPad Prism software (version 8.1.0).

For the representation of cells expressing τ biosensors, FastFLIM images and their corresponding intensity images acquired during FLIM experiments (as described above), were exported and further processed. The intensity image was tresholded (Otsu's method), converted into a binary mask and applied to the FastFLIM image using the Image calculator function. The processed FastFLIM was used for figure panels, representing the average photon arrival time. FastFLIM images were not used for quantitative analysis of fluorescence lifetimes. Fluorescence lifetimes used for quantitative analysis were determined as described in the section Fluorescence lifetime imaging microscopy.

**Statistics and reproducibility**

All *in vitro* measurements were performed at least in 3 technical replicates. All cell experiments were performed at least in 3 biological replicates (*i.e.* on 3 different days), unless stated differently. Statistical significance of a sample group over a reference group was determined by performing a two-tailed unpaired t-test with Welch's correction using the Software GraphPad Prism (version 8.1.0). Statistical analyses were performed on sample size > 30 cells (for fluorescence microscopy), assuming normal distribution according to the central limit theorem, or ≥ 6 wells (for bioluminescence). Sample sizes are indicated in the caption of corresponding figures. Statistical analysis: non-significant (n.s.) $p \geq 0.5$, * $p < 0.05$, ** $p < 0.01$, *** $p < 0.001$ and **** $p < 0.0001$. Microscopy images are representative snapshots of experiments.

#### Method Tables

**Table M1| Chemicals and reagents used in this study.**

| Chemical/Reagent | Manufacturer | Catalogue number |
|---|---|---|
| KOD Hot Start Master Mix | Sigma-Aldrich | 71842 |
| Q5® Site-Directed Mutagenesis Kit | NEB | E0554S |
| QIAprep Spin Miniprep Kit | Qiagen | 27106 |
| GeneJET Endo-Free Plasmid-Maxiprep-Kit | ThermoFisher | K0861 |
| Isopropyl-β-D-thiogalactopyranoside (IPTG) | Roth | CN084 |
| Phenylmethylsulfonyl fluoride (PMSF) | ThermoScientific | 36978 |
| Lysozyme | ThermoScientific | 89833 |
| HisPur™ Ni-NTA Superflow Agarose | ThermoScientific | 25217 |
| 4-20% Mini Protean TGX stain-free gel | Bio-Rad | 568094 |
| Amicon® Ultra 4 mL Centrifugal Filters | Merck | UFC803024 (30 kDa)<br>UFC805024 (50 kDa) |
| Glycerol | Merck | 356350 |
| Bovine serum albumin (BSA) | Roth | 01634 |
| 4-(2-hydroxyethyl)-1-piperazineethanesulfonic acid (HEPES) | Sigma-Aldrich | H4034 |
| Sodium chloride (NaCl) | Merck | 106404 |
| Dimethyl sulfoxide (DMSO) | Applichem | A36720100 |
| Calcium chloride | Roth | A1191 |
| ethylene glycol-bis(β-aminoethyl ether)-*N*,*N*,*N*′,*N*′-tetraacetic acid (EGTA) | Sigma-Aldrich | E4378 |
| Calcium Calibration buffer Kit #1 | Life technologies | C3008MP |
| SPG pH 4.0 - 1 M buffer | Jena Bioscience | CSS-389 |
| SPG pH 10.0 - 1 M buffer | Jena Bioscience | CSS-390 |
| Histamine | Sigma-Aldrich | H7250 |
| Ionomycin | Sigma-Aldrich | I9657 |
| 1x PBS pH 7.4 | Gibco | 10010015 |
| 1x HBSS with calcium and magnesium | Corning | 21-023-CMR |
| TrypLE™ Express | Gibco | 12604013 |
| DMEM high glucose +GlutaMAX™ | Gibco | 31966021 |
| DMEM high glucose, phenol red-free | Gibco | 31053028 |
| DMEM no glucose, phenol red-free | Gibco | A1443001 |
| Sodium pyruvate (100X) | Gibco | 11360070 |
| GlutaMAX™ Supplement (100x) | Gibco | 35050038 |
| Fetal bovine serum (FBS, heat-inactivated) | Gibco | 10500064 |
| Opti-MEM™, reduced serum | Gibco | 31985047 |
| Lipofectamine 3000 Transfection Reagent | Invitrogen | L3000001 |
| Adenosine-5′-triphosphate (ATP) magnesium salt | Sigma-Aldrich | A9187 |
| Adenosine-5′-diphosphate (ADP) disodium salt | Sigma-Aldrich | 1897 |
| Adenosine-5′-monophosphate (AMP) sodium salt | Sigma-Aldrich | A1752 |
| Guanosine-5′-triphosphate (GTP) sodium salt | Sigma-Aldrich | 10106399001 |
| 2-Deoxy-D-glucose (2DG) | TCI Chemicals | D0051 |
| D-glucose monohydrate | Roth | 6780 |
| Nicotinamide (NAM) | Sigma-Aldrich | 72340 |
| Nicotinamide riboside (NR) | Combi-Blocks | HB-5832 |
| Nicotinamide mononucleotide (NMN) | Sigma-Aldrich | N3501 |
| Nicotinamide adenine dinucleotide (NAD+) | Roche | 10127965001 |
| Nicotinamide adenine dinucleotide phosphate (NADP+) | Roth | AE13.3 |
| Nicotinamide adenine nucleotide, reduced (NADH) | Roth | AE12.2 |
| Nicotinic acid adenine dinucleotide (NAAD+) | Sigma-Aldrich | N4256 |

| | | |
|---|---|---|
| FK866 | Selleckchem | S2799 |
| *N*-methyl-*N*'-nitro-*N*-nitrosoguanidine (MNNG) | Biozol | N529925 |
| MitoTracker™ Red FM | Invitrogen | M22425 |
| Hoechst 33342 | Invitrogen | H3570 |
| Penicillin-Streptomycin (Pen/Strep) | Gibco | 15140122 |
| NanoBRET™ Nano-Glo Substrate | Promega | N157C |
| Extracellular NanoLuc® Inhibitor | Promega | N235A |
| Nano-Glo™ Substrate | Promega | N113B |
| DL-2-Amino-5-phosphonovaleric acid (APV) | SantaCruz | sc-201503 |
| NBQX disodium salt | Sigma-Aldrich | N183 |
| Black non-binding flat bottom 96 well plates | Perkin Elmer | 6005720 |
| Black low volume flat bottom 384 well plates | Corning | 3820 |
| White non-binding flat bottom 96 well plates | Perkin Elmer | 6005290 |
| White 96 well plate, cell culture treated | BrandTech | 782090 |
| White low volume flat bottom 384 well plates | Corning | 3824 |
| Black 96 well glass bottom imaging plate | IBL, Cellvis | P96-1.5H-N |
| Black 24 well glass bottom imaging plate | IBL, Cellvis | P24-1.5H-N |

**Table M2| Fluorophores used in this study.**

| Number | Structure | Name | $Ex_{max}/Em_{max}$ [nm] | Source, reference |
|---|---|---|---|---|
| 1 | 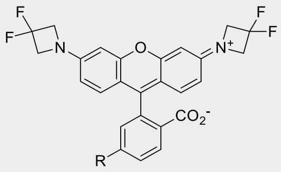 | $JF_{525}$-CA | 525/549 | Gift from Dr. Luke Lavis, HHMI, Ashburn, VA, USA[27] |
| 2 | 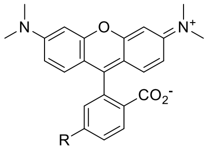 | TMR-CA | 548/572 | Purchased from Promega, Madison, WI, USA[28] |
| 3 | 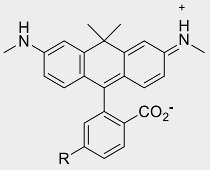 | 580CP-CA | 582/607 | Gift from Dr. Alexey N. Butkevich, MPI-MF, Heidelberg, Germany[29] |
| 4 | 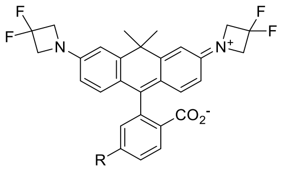 | $JF_{585}$-CA | 585/609 | Gift from Dr. Luke Lavis, HHMI, Ashburn, VA, USA[27] |
| 5 | 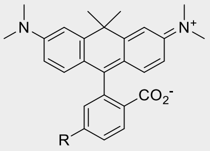 | CPY-CA | 606/626 | Butkevich *et al*.[30] |
| 6 | 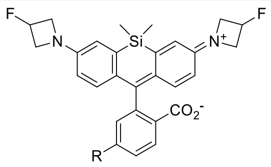 | $JF_{635}$-CA | 635/652 | Gift from Dr. Luke Lavis, HHMI, Ashburn, VA, USA[27] |
| 7 | 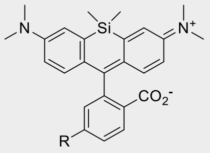 | SiR-halo (=SiR-CA) | 643/662 | Lukinavicius *et al*.[31] |
| 8 | 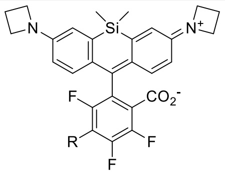 | $JF_{669}$-CA | 669/682 | Gift from Dr. Luke Lavis, HHMI, Ashburn, VA, USA[27] |
| 9 | 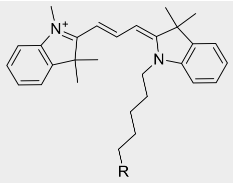 | Cy3-CA | 554/568 | Wilhelm and Kuehn *et al*.[9] |

| | | | | |
|---|---|---|---|---|
| 10 | 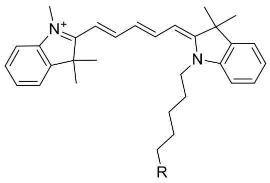 | Cy5-CA | 649/666 | Wilhelm and Kuehn *et al*.[9] |

R = 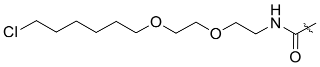 (Chloroalkane (CA))

JF = Janelia Fluor

**Table M3| Plasmids and stable cell lines used in this study.**

| Construct | Plasmid | Gene | Entry plasmids (Addgene#) | Addgene# | Stable cell line |
|---|---|---|---|---|---|
| pET-51b(+) HaloTag7-EGFP | pET-51b(+) | HaloTag7-EGFP | 167266[9], 130706[32] | n.a. | n.a. |
| pET-51b(+) ChemoG1 | pET-51b(+) | ChemoG1 | 167266[9], 130706[32] | 193799 | n.a. |
| pET-51b(+) ChemoG1$^{Y39A}$ | pET-51b(+) | ChemoG1$^{Y39A}$ | ChemoG1 | n.a. | n.a. |
| pET-51b(+) ChemoG1$^{K41A}$ | pET-51b(+) | ChemoG1$^{K41A}$ | ChemoG1 | n.a. | n.a. |
| pET-51b(+) ChemoG1$^{F223R}$ | pET-51b(+) | ChemoG1$^{F223R}$ | ChemoG1 | n.a. | n.a. |
| pET-51b(+) ChemoG2 | pET-51b(+) | ChemoG2 | ChemoG1 | 193800 | n.a. |
| pET-51b(+) ChemoG3 | pET-51b(+) | ChemoG3 | ChemoG2 | 193801 | n.a. |
| pET-51b(+) ChemoG3.1 | pET-51b(+) | ChemoG3 | ChemoG2 | 193802 | n.a. |
| pET-51b(+) ChemoG3.2 | pET-51b(+) | ChemoG3 | ChemoG2 | 193803 | n.a. |
| pET-51b(+) ChemoG4 | pET-51b(+) | ChemoG4 | ChemoG3 | 193804 | n.a. |
| pET-51b(+) ChemoG5 | pET-51b(+) | ChemoG5 | ChemoG4 | 193805 | n.a. |
| pET-51b(+) ChemoB | pET-51b(+) | ChemoB | 54572[33] | n.a. | n.a. |
| pET-51b(+) ChemoC | pET-51b(+) | ChemoC | 48203[34] | n.a. | n.a. |
| pET-51b(+) ChemoY | pET-51b(+) | ChemoY | 39813[35] | n.a. | n.a. |
| pET-51b(+) ChemoR | pET-51b(+) | ChemoR | 85042[36] | n.a. | n.a. |
| pCDNA5/FRT-ChemoG1 | pCDNA5/FRT | ChemoG1 | 167266[9], 130706[32] | 193806 | n.a. |
| pCDNA5/FRT-ChemoG2 | pCDNA5/FRT | ChemoG2 | ChemoG1 | n.a. | n.a. |
| pCDNA5/FRT-ChemoG3 | pCDNA5/FRT | ChemoG3 | ChemoG2 | n.a. | n.a. |
| pCDNA5/FRT-ChemoG4 | pCDNA5/FRT | ChemoG4 | ChemoG3 | n.a. | n.a. |
| pCDNA5/FRT-ChemoG5 | pCDNA5/FRT | ChemoG5 | ChemoG4 | 193807 | n.a. |
| pCDNA5/FRT-ChemoB | pCDNA5/FRT | ChemoB | 54572[33] | 193808 | n.a. |
| pCDNA5/FRT-ChemoC | pCDNA5/FRT | ChemoC | 48203[34] | 193809 | n.a. |
| pCDNA5/FRT-ChemoY | pCDNA5/FRT | ChemoY | 39813[35] | 193810 | n.a. |
| pCDNA5/FRT-ChemoR | pCDNA5/FRT | ChemoR | 85042[36] | 193811 | n.a. |
| pCDNA5/FRT-NES-ChemoG5 | pCDNA5/FRT | NES-ChemoG5 | ChemoG5 | n.a. | n.a. |
| pCDNA5/FRT-ChemoG5-PDGFR$_{tm}$ | pCDNA5/FRT | ChemoG5-PDGFR$_{tm}$ | ChemoG5, in-house plasmid[37] | n.a. | n.a. |
| pCDNA5/FRT-ChemoG5-NLS3x | pCDNA5/FRT | ChemoG5-NLS3x | ChemoG5 | n.a. | n.a. |
| pCDNA5/FRT-[2xCox8]-ChemoG5 | pCDNA5/FRT | [2xCox8]-ChemoG5 | ChemoG5, 113916[4] | n.a. | n.a. |
| pCDNA5/FRT-ChemoG5-LaminB1 | pCDNA5/FRT | ChemoG5-LaminB1 | 55069 | n.a. | n.a. |
| pET-51b(+) EGFP-CaM-P30-M13-HaloTag7 | pET-51b(+) | EGFP-CaM-P30-M13-HaloTag7 | 40755[38] | n.a. | n.a. |
| pET-51b(+) EGFP$^{A206K}$-CaM-P30-M13-HaloTag7 | pET-51b(+) | EGFP$^{A206K}$-CaM-P30-M13-HaloTag7 | 40755[38] | n.a. | n.a. |
| pET-51b(+) ChemoG-CaM | pET-51b(+) | ChemoG-CaM | 40755[38] | n.a. | n.a. |
| pET-51b(+) EGFP$^{A206K}$-CaM-P30-M13-HaloTag7$^{E143R-E147R-L271E}$ | pET-51b(+) | EGFP$^{A206K}$-CaM-P30-M13-HaloTag7$^{E143R-E147R-L271E}$ | 40755[38] | n.a. | n.a. |
| pET-51b(+) EGFP$^{A206K-T225R}$-CaM-P30-M13-HaloTag7$^{E143R-E147R-L271E}$ | pET-51b(+) | EGFP$^{A206K-T225R}$-CaM-P30-M13-HaloTag7$^{E143R-E147R-L271E}$ | 40755[38] | n.a. | n.a. |
| pET-51b(+) ChemoB-CaM | pET-51b(+) | ChemoB-CaM | 40755[38] | 193812 | n.a. |
| pET-51b(+) ChemoC-CaM | pET-51b(+) | ChemoC-CaM | 40755[38] | 193813 | n.a. |
| pET-51b(+) ChemoY-CaM | pET-51b(+) | ChemoY-CaM | 40755[38] | 193814 | n.a. |

| | | | | | |
|---|---|---|---|---|---|
| pET-51b(+) ChemoR-CaM | pET-51b(+) | ChemoR-CaM | 40755[38] | 193815 | n.a. |
| pET-51b(+) YC 3.6 | pET-51b(+) | YC 3.6 | 51966[8] | n.a. | n.a. |
| pCDNA5/FRT-ChemoG-CaM | pCDNA5/FRT | ChemoG-CaM | 40755[38] | 193816 | n.a. |
| pGP-AAV2-hSyn1-NES-ChemoG-CaM | pGP-AAV2 | NES-ChemoG-CaM | 101061[2] | 193817 | n.a. |
| pET-51b(+) EGFP$^{A206K}$-$F_OF_1$-HT7 | pET-51b(+) | EGFP$^{A206K}$-$F_OF_1$-HT7 | 51958[39] | n.a. | n.a. |
| pET-51b(+) ChemoG-ATP | pET-51b(+) | ChemoG-ATP | 51958[39] | n.a. | U-2 OS Flp-In T-REx |
| pET-51b(+) EGFP$^{A206K-T225R}$-$F_OF_1$-HT7 | pET-51b(+) | EGFP$^{A206K-T225R}$-$F_OF_1$-HT7 | 51958[39] | n.a. | n.a. |
| pET-51b(+) EGFP$^{A206K-T225R}$-$F_OF_1$-HT7$^{L271E}$ | pET-51b(+) | EGFP$^{A206K-T225R}$-$F_OF_1$-HT7$^{L271E}$ | 51958[39] | n.a. | n.a. |
| pET-51b(+) ChemoB-ATP | pET-51b(+) | ChemoB-ATP | 51958[39] | n.a. | n.a. |
| pET-51b(+) ChemoR-ATP | pET-51b(+) | ChemoR-ATP | 51958[39] | n.a. | n.a. |
| pET-51b(+) ATeam 1.03 | pET-51b(+) | ATeam 1.03 | 51958[39] | n.a. | n.a. |
| pCDNA5/FRT-ChemoG-ATP | pCDNA5/FRT | ChemoG-ATP | 51958[39] | 193818 | n.a. |
| pCDNA5/FRT-ChemoB-ATP | pCDNA5/FRT | ChemoB-ATP | 51958[39] | 193819 | n.a. |
| pCDNA5/FRT-ChemoR-ATP | pCDNA5/FRT | ChemoR-ATP | 51958[39] | 193820 | n.a. |
| pCDNA5/FRT/TO-ATeam 1.03 | pCDNA5/FRT | ATeam 1.03 | 51958[39] | n.a. | U-2 OS Flp-In T-REx |
| pET-51b(+) EGFP$^{A206K}$-ttLigA$^{K118L-D289N}$-HT7 | pET-51b(+) | EGFP$^{A206K}$-ttLigA$^{K118L-D289N}$-HT7 | - | n.a. | n.a. |
| pET-51b(+) EGFP$^{A206K}$-ttLigA$^{K118L-D289N-V292A}$-HT7 | pET-51b(+) | EGFP$^{A206K}$-ttLigA$^{K118L-D289N-V292A}$-HT7 | - | n.a. | n.a. |
| pET-51b(+) EGFP$^{A206K}$-ttLigA$^{K118L-Y226W-D289N-V292A}$-HT7 | pET-51b(+) | EGFP$^{A206K}$-ttLigA$^{K118L-Y226W-D289N-V292A}$-HT7 | - | n.a. | n.a. |
| pET-51b(+) EGFP$^{A206K-T225R}$-ttLigA$^{K118L-Y226W-D289N-V292A}$-HT7 | pET-51b(+) | EGFP$^{A206K-T225R}$-ttLigA$^{K118L-Y226W-D289N-V292A}$-HT7 | - | n.a. | n.a. |
| pET-51b(+) ChemoG-NAD | pET-51b(+) | ChemoG-NAD | - | n.a. | n.a. |
| pET-51b(+) EGFP$^{A206K-T225R}$-ttLigA$^{K118L-Y226W-D289N-V292A}$-HT7$^{E143R-E147R-L271E}$ | pET-51b(+) | EGFP$^{A206K-T225R}$-ttLigA$^{K118L-Y226W-D289N-V292A}$-HT7$^{E143R-E147R-L271E}$ | - | n.a. | n.a. |
| pET-51b(+) ChemoB-NAD | pET-51b(+) | ChemoB-NAD | - | n.a. | n.a. |
| pET-51b(+) ChemoR-NAD | pET-51b(+) | ChemoR-NAD | - | n.a. | n.a. |
| pCDNA5/FRT-ChemoG-NAD | pCDNA5/FRT | ChemoG-NAD | - | 193821 | U-2 OS Flp-In T-REx |
| pCDNA5/FRT-ChemoG-NAD-NLS3x | pCDNA5/FRT | ChemoG-NAD-NLS3x | - | 193822 | U-2 OS Flp-In T-REx |
| pCDNA5/FRT/TO-[4xCox8]-ChemoG-NAD | pCDNA5/FRT/TO | [4xCox8]-ChemoG-NAD | - | 193823 | U-2 OS Flp-In T-REx |
| pCDNA5/FRT-ChemoB-NAD | pCDNA5/FRT | ChemoB-NAD | - | 193824 | U-2 OS Flp-In T-REx |
| pCDNA5/FRT-ChemoB-NAD-NLS3x | pCDNA5/FRT | ChemoB-NAD-NLS3x | - | n.a. | n.a. |
| pCDNA5/FRT-ChemoR-NAD | pCDNA5/FRT | ChemoR-NAD | - | 193825 | U-2 OS Flp-In T-REx |
| pCDNA5/FRT/TO-ChemoB-NAD[hsOpti]-NLS3x-T2A-[4xCox8]-ChemoG-NAD[zfOpti] | pCDNA5/FRT/TO | ChemoB-NAD[hsOpti]-NLS3x-T2A-4xCox8a-ChemoG-NAD[zfOpti] | - | n.a. | n.a. |
| pET-51b(+) ChemoD-NAD | pET-51b(+) | ChemoD-NAD | 104620[40] | n.a. | n.a. |
| pCDNA5/FRT-ChemoD-NAD | pCDNA5/FRT | ChemoD-NAD | 104620[40] | 193826 | U-2 OS Flp-In T-REx |
| pET-51b(+) ChemoL-NAD | pET-51b(+) | ChemoL-NAD | 117909[41] | n.a. | U-2 OS Flp-In T-REx |
| pET-51b(+) ChemoL-CaM | pET-51b(+) | ChemoL-CaM | 117909[41] | n.a. | n.a. |
| pET-51b(+) ChemoL-ATP | pET-51b(+) | ChemoL-ATP | 117909[41] | n.a. | n.a. |
| pCDNA5/FRT-ChemoL-NAD | pCDNA5/FRT | ChemoL-NAD | 117909[41] | 193827 | n.a. |
| pCDNA5/FRT-ChemoL-NAD-NLS3x | pCDNA5/FRT | ChemoL-NAD-NLS3x | 117909[41] | 193828 | n.a. |
| pCDNA5/FRT-[4xCox8]-ChemoL-NAD | pCDNA5/FRT | [4xCox8]-ChemoL-NAD | 117909[41] | 193829 | n.a. |
| pCDNA5/FRT-ChemoL-CaM | pCDNA5/FRT | ChemoL-CaM | 117909[41] | 193830 | n.a. |
| pCDNA5/FRT-ChemoL-ATP | pCDNA5/FRT | ChemoL-ATP | 117909[41] | 193831 | n.a. |
| pET-51b(+) His-TEV-ChemoG1 | pET-51b(+) | ChemoG1 | 167266[9] | n.a. | n.a. |
| pET-51b(+) His-TEV-ChemoG5 | pET-51b(+) | ChemoG5 | 167266[9] | n.a. | n.a. |
| pET-51b(+) His-TEV-HaloTag7 | pET-51b(+) | HT7 | | 167266[9] | n.a. |

n.a. = not available.

**Table M4| Data collection and refinement statistics.**

| | HaloTag7-Cy3<br>8B6R | ChemoG1-TMR<br>8B6S | ChemoG5-TMR<br>8B6T |
|---|---|---|---|
| **Data collection** | | | |
| Space group | $P4_22_12$ | $P1$ | $P12_11$ |
| Unit-cell parameters | | | |
| *a, b, c* (Å) | 112.56, 112.56, 44.33 | 46.19, 63.71, 89.42 | 46.60, 64.04, 172.95 |
| $\alpha, \beta, \gamma$ (°) | 90.00, 90.00, 90.00 | 93.56, 91.02, 90.85 | 90.00, 97.67, 90.00 |
| Radiation source | PXII-X10SA, SLS | PXII-X10SA, SLS | PXII-X10SA, SLS |
| Wavelength (Å) | 0.99988 | 0.99996 | 0.99992 |
| Temperature (K) | 100 | 100 | 100 |
| Resolution range (Å) | 50-1.50 (1.60-1.50) | 50-1.80 (1.90-1.80) | 50-2.00 (2.10-2.00) |
| No. of observed reflections | 341056 (60343) | 182229 (26711) | 216345 (30251) |
| No. of unique reflections | 46121 (7965) | 89852 (13310) | 66470 (9089) |
| Multiplicity | 7.4 (7.6) | 2.0 (2.0) | 3.3 (3.3) |
| Completeness (%) | 99.9 (99.9) | 95.3 (94.3) | 97.0 (97.8) |
| $R_{merge}$ (%) | 6.8 (65.7) | 4.1 (40.0) | 8.6 (41.0) |
| $<I/\sigma(I)>$ | 18.2 (3.4) | 12.0 (2.1) | 8.5 (3.4) |
| $CC_{1/2}$ (%)[#] | 99.9 (90.2) | 99.8 (75.2) | 99.5 (87.4) |
| **Refinement** | | | |
| Molecules per a.u. | 1 | 2 | 2 |
| No. of reflections | 46120 | 89842 | 66470 |
| No. of reflections in test set | 2306 | 4492 | 3399 |
| Resolution range (Å) | 41.25-1.50 | 46.18-1.80 | 46.18-2.00 |
| No. of non-hydrogen atoms | | | |
| Protein | 2365 | 8273 | 8276 |
| Ligand/ion | 72 | 146 | 134 |
| Water | 297 | 460 | 308 |
| Total | 2734 | 8879 | 8718 |
| *R* (%) | 16.20 | 17.28 | 22.06 |
| $R_{free}$ (%) | 19.19 | 20.08 | 24.51 |
| RMS deviations from ideal | | | |
| bonds (Å) | 0.013 | 0.007 | 0.002 |
| angles (°) | 1.229 | 1.094 | 0.779 |
| *B*-factors ($Å^2$) | | | |
| Protein | 14.80 | 26.03 | 20.62 |
| Ligand/ion | 23.87 | 22.14 | 17.73 |
| Water | 24.40 | 29.14 | 19.67 |
| Average | 16.08 | 26.13 | 20.54 |
| Wilson B ( $Å^2$) | 14.42 | 24.95 | 22.88 |
| Ramachandran statistics (%) | | | |
| favored regions | 95.9 | 97.5 | 96.8 |
| allowed regions | 4.1 | 2.5 | 3.2 |
| disallowed regions | 0 | 0 | 0 |
| Clashscore | 1.04 | 1.33 | 3.03 |

[#]as implemented in XDS[42]. Values in parentheses are for the highest resolution shell.

**Table M5| Spectral settings for fluorescence spectroscopy measurements.**

| Chromophore/fluorophore | Max. emission wavelength | Excitation wavelength used | Emission wavelength range measured |
|---|---|---|---|
| EBFP2 | 446 nm | 360 nm | 400-800 nm |
| mCerulean3 | 474 nm | 400 nm | 440-800 nm |
| EGFP | 510 nm | 440 nm | 480-800 nm |
| Venus | 528 nm | 460 nm | 494-800 nm |
| mKO2 | 566 nm | 510 nm | 550-800 nm |
| TagRFP | 584 nm | 510 nm | 550-800 nm |
| mRuby2 | 594 nm | 510 nm | 550-800 nm |
| mRuby3 | 594 nm | 510 nm | 550-800 nm |
| mScarlet | 594 nm | 520 nm | 560-800 nm |
| mCherry | 610 nm | 530 nm | 570-800 nm |
| $JF_{525}$ | 554 nm | - | - |
| TMR | 576 nm | - | - |
| Cy3 | 576 nm | - | - |
| 580CP | 606 nm | - | - |
| $JF_{585}$ | 612 nm | - | - |
| CPY | 628 nm | 580 nm | 610-750 nm |
| $JF_{635}$ | 662 nm | 610 nm | 640-750 nm |
| Cy5 | 664 nm | - | - |
| SiR | 666 nm | 610 nm | 640-750 nm |
| $JF_{669}$ | 688 nm | - | - |
| YC 3.6 | 474/528 nm | 400 nm | 440-650 nm |
| ATeam 1.03 | 474/528 nm | 400 nm | 440-650 nm |

**Table M6| Analyte concentration ranges used for sensor titrations.**

| Experiment | Analyte | 10x concentration (range) | Final 1x concentration (range) |
|---|---|---|---|
| ChemoX-CaM titration with free $Ca^{2+}$ (Fig. 2c, d, S7d, g, j) | Free $Ca^{2+}$ | - | 10 nM – 39 µM* |
| ChemoX-CaM response to free $Ca^{2+}$ at different pH (Fig. 2c, d, S7d, g, j) | $CaCl_2$ | 20 mM | 2 mM |
| | EGTA | 20 mM | 2 mM |
| RFP-based calcium sensor responses to free $Ca^{2+}$ (Fig. S7) | $CaCl_2$ | 20 mM | 2 mM |
| | EGTA | 20 mM | 2 mM |
| ChemoX-ATP titration with ATP or structurally similar analytes (Fig. 3c, S9c) | ATP | 0.1 - 100 mM | 0.01 - 10 mM |
| | ADP | 0.1 - 100 mM | 0.01 - 10 mM |
| | AMP | 0.1 - 100 mM | 0.01 - 10 mM |
| | GTP | 0.1 - 100 mM | 0.01 - 10 mM |
| ChemoX-NAD titration with $NAD^+$ or structurally similar analytes (Fig. 4c, d, S10e) | $NAD^+$ | 10 nM – 100 mM | 1 nM – 10 mM |
| | NAM | 10 nM – 100 mM | 1 nM – 10 mM |
| | NR | 10 nM – 100 mM | 1 nM – 10 mM |
| | NMN | 10 nM – 100 mM | 1 nM – 10 mM |
| | NADH | 10 nM – 100 mM | 1 nM – 10 mM |
| | $NADP^+$ | 10 nM – 100 mM | 1 nM – 10 mM |
| | $NAAD^+$ | 10 nM – 100 mM | 1 nM – 10 mM |
| | ATP | 10 nM – 100 mM | 1 nM – 10 mM |
| | ADP | 10 nM – 100 mM | 1 nM – 10 mM |
| ChemoX-NAD titration with $NAD^+$ in presence of structurally similar analytes (Fig. S10f, g) | $NAD^+$ (titration) | 10 nM – 100 mM | 1 nM – 10 mM |
| | NAM (constant) | 10 mM | 1 mM |
| | NR (constant) | 1 mM | 0.1 mM |
| | NMN (constant) | 1 mM | 0.1 mM |
| | NADH (constant) | 1 mM | 0.1 mM |
| | $NAAD^+$ (constant) | 1 mM | 0.1 mM |
| | $NADP^+$ (constant) | 1 mM | 0.1 mM |
| | ATP (constant) | 10 mM | 1 mM |
| | ADP (constant) | 10 mM | 1 mM |
| | AMP (constant) | 10 mM | 1 mM |
| ChemoD-NAD titration with $NAD^+$ (intensiometric, Fig. 5c, S15b, d, e) | $NAD^+$ | 100 nM – 100 mM | 10 nM – 10 mM |
| ChemoD-NAD titration with $NAD^+$ (fluorescence lifetime, Fig. 5f, S15f-i) | $NAD^+$ | 1 µM – 100 mM | 100 nM – 10 mM |
| ChemoL-NAD titration with $NAD^+$ (Fig. 6c) | $NAD^+$ | 100 nM – 100 mM | 10 nM – 10 mM |
| ChemoL-CaM titration with free $Ca^{2+}$ (Fig. S16d) | Free $Ca^{2+}$ | - | 50 nM – 39 µM* |
| ChemoL-ATP titration with ATP (Fig. S16f) | $NAD^+$ | 100 nM – 100 mM | 10 nM – 10 mM |

*For titrations of calcium sensors, special calcium buffers with defined concentrations of free $Ca^{2+}$ were prepared (see Analyte titrations of biosensors below for details).

**Table M7| Settings for confocal and widefield fluorescence microscopy.**

| Figure | Construct | Label | Microscope | Objective | Excitation [nm] | Emission [nm] | Pixel dwell time [µs] | Size [pixels] | Z size [µm] |
|---|---|---|---|---|---|---|---|---|---|
| 1f | ChemoG5-NLS | - | Confocal | 40x/1.10 water | 480 | 490-540 | 3.16 | 512x512 | 5 |
| | | TMR | | | 480 | 490-540/550-600 | 3.16 | 512x512 | 5 |
| | | CPY | | | 480 | 490-540/620-670 | 3.16 | 512x512 | 5 |
| | | SiR | | | 480 | 490-540/650-700 | 3.16 | 512x512 | 5 |
| 1h | ChemoB | SiR | Confocal | 40x/1.10 water | 405 | 420-470/650-700 | 3.16 | 512x512 | 5 |
| | ChemoC | SiR | | | 405 | 460-500/650-700 | 3.16 | 512x512 | 5 |
| | ChemoG5 | SiR | | | 480 | 490-540/650-700 | 3.16 | 512x512 | 5 |
| | ChemoY | SiR | | | 505 | 515-565/650-700 | 3.16 | 512x512 | 5 |
| | ChemoR | SiR | | | 550 | 570-620/650-700 | 3.16 | 512x512 | 5 |
| 2e | ChemoG-CaM | SiR | Widefield | 20x/0.80 dry | 470* | 525/50, 700/75** | n.d. | 512x512 | 2 |
| 2g | ChemoG-CaM | SiR | Widefield | 20x/0.80 dry | 470* | 525/50, 700/75** | n.d. | 256x256 | 0 |
| 3d | ChemoG-ATP | SiR | Confocal | 40x/1.10 water | 480 | 490-540/650-700 | 3.16 | 512x512 | 5 |
| 3f | ChemoB-ATP | SiR | Confocal | 40x/1.10 water | 405 | 420-470/650-700 | 3.16 | 512x512 | 5 |
| | ChemoG-ATP | SiR | | | 480 | 490-540/650-700 | 3.16 | 512x512 | 5 |
| | ChemoR-ATP | SiR | | | 550 | 570-620/650-700 | 3.16 | 512x512 | 5 |
| | ATeam 1.03 | - | | | 405 | 460-500/520-560 | 3.16 | 512x512 | 5 |
| 4e | ChemoG-NAD | SiR | Confocal | 40x/1.10 water | 480 | 490-540/650-700 | 3.16 | 512x512 | 0 |
| 4g | ChemoB-NAD-cyto | SiR | Confocal | 40x/1.10 water | 405 | 420-460/650-700 | 3.84 | 2048x2048 | 0 |
| | ChemoG-NAD-mito | SiR | | | 480 | 490-540/650-700 | 3.84 | 2048x2048 | 0 |
| 4h | ChemoB-NAD-cyto | SiR | Confocal | 40x/1.10 water | 405 | 420-460/650-700 | 3.16 | 512x512 | 4 |
| | ChemoG-NAD-mito | SiR | | | 480 | 490-540/650-700 | 3.16 | 512x512 | 4 |
| 5f | ChemoD-NAD | CPY | Confocal | 40x/1.10 water | 610 | 620-670 | 1.75 | 512x512 | 0 |
| | | $JF_{635}$ | | | 640 | 650-700 | 1.75 | 512x512 | 0 |
| | | SiR | | | 625 | 635-685 | 1.75 | 512x512 | 0 |
| 5g/h | ChemoD-NAD | CPY | Confocal | 40x/1.10 water | 610 | 620-670 | 7.69 | 512x512 | 0 |
| S4a | HT-EGFP | SiR | Confocal | 40x/1.10 water | 480 | 490-540/650-700 | 3.16 | 512.x512 | 5 |
| | ChemoG1-5 | SiR | | | 480 | 490-540/650-700 | 3.16 | 512.x512 | 5 |
| | ChemoG5-NES | SiR | | | 480 | 490-540/650-700 | 3.16 | 512.x512 | 5 |
| | $ChemoG5\text{-}PDGFR_{tm}$ | SiR | | | 480 | 490-540/650-700 | 3.16 | 512.x512 | 0 |
| | ChemoG5-NLS | SiR | | | 480 | 490-540/650-700 | 3.16 | 512.x512 | 5 |
| | ChemoG5-Cox8 | SiR | | | 480 | 490-540/650-700 | 3.16 | 512.x512 | 0 |
| | ChemoG5-LaminB1 | SiR | | | 480 | 490-540/650-700 | 3.16 | 512.x512 | 0 |
| S12a | ChemoG-NAD-NLS | SiR | Confocal | 40x/1.10 water | 480 | 490/540/650-700 | 3.16 | 512x512 | 0 |
| S12d | ChemoG-NAD-mito | SiR | Confocal | 40x/1.10 water | 480 | 490/540/650-700 | 3.16 | 512x512 | 0 |
| S13a | ChemoB-NAD | SiR | Confocal | 40x/1.10 water | 405 | 420-470/650/700 | 3.16 | 512x512 | 0 |

| | | | | | | | | | |
|---|---|---|---|---|---|---|---|---|---|
| S13c | ChemoR-NAD | SiR | Confocal | 40x/1.10 water | 550 | 570-620/650-700 | 3.16 | 512x512 | 0 |
| S14a | ChemoB-NAD-cyto | SiR | Confocal | 40x/1.10 water | 405 | 420-460/650-700 | 3.16 | 512x512 | 4 |
| | ChemoG-NAD-mito | SiR | | | 480 | 490-540/650-700 | 3.16 | 512x512 | 4 |
| S14b | ChemoB-NAD-NLS | SiR | Confocal | 40x/1.10 water | 405 | 420-460/650-700 | 3.16 | 512x512 | 0 |
| | ChemoG-NAD-mito | SiR | | | 480 | 490-540/650-700 | 3.16 | 512x512 | 0 |
| S14c | ChemoB-NAD-NLS | SiR | Confocal | 40x/1.10 water | 405 | 420-460/650-700 | 3.16 | 512x512 | 4 |
| | ChemoG-NAD-mito | SiR | | | 480 | 490-540/650-700 | 3.16 | 512x512 | 4 |
| S14d | ChemoB-NAD-NLS | SiR | Confocal | 40x/1.10 water | 405 | 420-460/650-700 | 3.16 | 512x512 | 4 |
| | ChemoG-NAD-mito | SiR | | | 480 | 490-540/650-700 | 3.16 | 512x512 | 4 |
| S15f/g | ChemoG-NAD | SiR | Confocal | 40x/1.10 water | 640 | 650-700 | 1.75 | 512x512 | 0 |
| | ChemoD-NAD | SiR | | | 640 | 650-700 | 1.75 | 512x512 | 0 |
| S15h | ChemoD-NAD | CPY | Confocal | 40x/1.10 water | 610 | 620-670 | 1.75 | 512x512 | 0 |
| S15i | ChemoD-NAD | $JF_{635}$ | Confocal | 40x/1.10 water | 620 | 635-685 | 1.75 | 512x512 | 0 |
| S15j/k | ChemoD-NAD | SiR | Confocal | 40x/1.10 water | 640 | 650-700 | 7.69 | 512x512 | 0 |
| S15l/m | ChemoD-NAD | $JF_{635}$ | Confocal | 40x/1.10 water | 620 | 635-685 | 7.69 | 512x512 | 0 |

*470 nm LED was used together with a 474/24 nm bandpass filter. **525/50 nm and 700/75 nm bandpass filters were used for acquisition of EGFP and FRET(SiR) fluorescence, respectively

#### Supplementary Methods

##### Protein sequences

###### *Static FRET constructs*

>ChemoG5
**MVSKGEELFTGVVPILVELDGDVNGHKFSVSGEGEGDATYGKLTLKFICTTGKLPVPWPTLVTTLTYGVQCFSRYPD**
**HMKQHDFFKSAMPEGYVQERTIFFKDDGNYKTRAEVKFEGDTLVNRIELKGIDFKEDGNILGHKLEYNYNSHNVYIM**
**ADKQKNGIKVNFKIRHNIEDGSVQLADHYQQNTPIGDGPVLLPDNHYLSTQSKLSKDPNEKRDHMVLLEFVRAAGIT**
**LGMDELYK**IGTGFPFDPHYVEVLGERMHYVDVGPRDGTPVLFLHGNPTSSYVWRNIIPHVAPTHRCIAPDLIGMGKS
DKPDLGYFFDDHVRFMDAFIEALGLEEVVLVIHDWGSALGFHWAKRNPERVKGIAFMEFIRPIPTWDEWPRFARRTF
QAFRTTDVGRKLIIDQNVFIEGTLPMGVVRPLTEVEMDHYREPFLNPVDREPLWRFPNELPIAGEPANIVALVEEYM
DWLHQSPVPKLLFWGTPGVLIPPAEAARLAKSLPNCKAVDIGPGENLLQEDNPDLIGSEIARWLSTLEISG

>ChemoB
**MVSKGEELFTGVVPILVELDGDVNGHKFSVRGEGEGDATYGKLTLKFICTTGKLPVPWPTLVTTLSHGVQCFARYPD**
**HMKQHDFFKSAMPEGYVQERTIFFKDDGTYKTRAEVKFEGDTLVNRIELKGVDFKEDGNILGHKLEYNFNSHNIYIM**
**AVKQKNGIKVNFKIRHNVEDGSVQLADHYQQNTPIGDGPVLLPDSHYLSTQSKLSKDPNEKRDHMVLLEFRRAAGIT**
**LGMDELYK**IGTGFPFDPHYVEVLGERMHYVDVGPRDGTPVLFLHGNPTSSYVWRNIIPHVAPTHRCIAPDLIGMGKS
DKPDLGYFFDDHVRFMDAFIEALGLEEVVLVIHDWGSALGFHWAKRNPERVKGIAFMEFIRPIPTWDEWPRFARRTF
QAFRTTDVGRKLIIDQNVFIEGTLPMGVVRPLTEVEMDHYREPFLNPVDREPLWRFPNELPIAGEPANIVALVEEYM
DWLHQSPVPKLLFWGTPGVLIPPAEAARLAKSLPNCKAVDIGPGENLLQEDNPDLIGSEIARWLSTLEISG

>ChemoC
**MVSKGEELFTGVVPILVELDGDVNGHKFSVSGEGEGDATYGKLTLKFICTTGKLPVPWPTLVTTLSWGVQCFARYPD**
**HMKQHDFFKSAMPEGYVQERTIFFKDDGNYKTRAEVKFEGDTLVNRIELKGIDFKEDGNILGHKLEYNAIHGNVYIT**
**ADKQKNGIKANFGLNCNIEDGSVQLADHYQQNTPIGDGPVLLPDNHYLSTQSKLSKDPNEKRDHMVLLEFVRAAGIT**
**LGMDELYK**IGTGFPFDPHYVEVLGERMHYVDVGPRDGTPVLFLHGNPTSSYVWRNIIPHVAPTHRCIAPDLIGMGKS
DKPDLGYFFDDHVRFMDAFIEALGLEEVVLVIHDWGSALGFHWAKRNPERVKGIAFMEFIRPIPTWDEWPRFARRTF
QAFRTTDVGRKLIIDQNVFIEGTLPMGVVRPLTEVEMDHYREPFLNPVDREPLWRFPNELPIAGEPANIVALVEEYM
DWLHQSPVPKLLFWGTPGVLIPPAEAARLAKSLPNCKAVDIGPGENLLQEDNPDLIGSEIARWLSTLEISG

>ChemoY
**MVSKGEELFTGVVPILVELDGDVNGHKFSVSGEGEGDATYGKLTLKLICTTGKLPVPWPTLVTTLGYGLQCFARYPD**
**HMKQHDFFKSAMPEGYVQERTIFFKDDGNYKTRAEVKFEGDTLVNRIELKGIDFKEDGNILGHKLEYNYNSHNVYIT**
**ADKQKNGIKANFKIRHNIEDGGVQLADHYQQNTPIGDGPVLLPDNHYLSYQSKLSKDPNEKRDHMVLLEFVRAAGIT**
**LGMDELYK**IGTGFPFDPHYVEVLGERMHYVDVGPRDGTPVLFLHGNPTSSYVWRNIIPHVAPTHRCIAPDLIGMGKS
DKPDLGYFFDDHVRFMDAFIEALGLEEVVLVIHDWGSALGFHWAKRNPERVKGIAFMEFIRPIPTWDEWPRFARRTF
QAFRTTDVGRKLIIDQNVFIEGTLPMGVVRPLTEVEMDHYREPFLNPVDREPLWRFPNELPIAGEPANIVALVEEYM
DWLHQSPVPKLLFWGTPGVLIPPAEAARLAKSLPNCKAVDIGPGENLLQEDNPDLIGSEIARWLSTLEISG

>ChemoR
**MVSKGEAVIKEFMRFKVHMEGSMNGHEFEIEGEGEGRPYEGTQTAKLKVTKGGPLPFSWDILSPQFMYGSRAFTKHP**
**ADIPDYYKQSFPEGFKWERVMNFEDGGAVTVTQDTSLEDGTLIYKVKLRGTNFPPDGPVMQKKTMGWEASTERLYPE**
**DGVLKGDIKMALRLKDGGRYLADFKTTYKAKKPVQMPGAYNVDRKLKITSHNEDYTVVEQYERSEGRHSTGGMDELY**
**K**IGTGFPFDPHYVEVLGERMHYVDVGPRDGTPVLFLHGNPTSSYVWRNIIPHVAPTHRCIAPDLIGMGKSDKPDLGY
FFDDHVRFMDAFIEALGLEEVVLVIHDWGSALGFHWAKRNPERVKGIAFMEFIRPIPTWDEWPEFARETFQAFRTTD
VGRKLIIDQNVFIEGTLPMGVVRPLTEVEMDHYREPFLNPVDREPLWRFPNELPIAGEPANIVALVEEYMDWLHQSP
VPKLLFWGTPGVLIPPAEAARLAKSLPNCKAVDIGPGLNLLQEDNPDLIGSEIARWLSTLEISG

EGFP, EBFP2, mCerulean3, Venus, mScarlet
HaloTag7
Interface mutations (XFP$^{A206K}$, XFP$^{T225R}$, HT7$^{E143R}$, HT7$^{E147R}$, HT7$^{L271E}$, mScarlet$^{D201K}$, EBFP2$^{N39Y}$)

**Calcium sensors**

>ChemoG-CaM
MVSKGEELFTGVVPILVELDGDVNGHKFSVSGEGEGDATYGKLTLKFICTTGKLPVPWPTLVTTLTYGVQCFSRYPD
HMKQHDFFKSAMPEGYVQERTIFFKDDGNYKTRAEVKFEGDTLVNRIELKGIDFKEDGNILGHKLEYNYNSHNVYIM
ADKQKNGIKVNFKIRHNIEDGSVQLADHYQQNTPIGDGPVLLPDNHYLSTQSKLSKDPNEKRDHMVLLEFVTAAGIT
GGTLPDQLTEEQIAEFKEAFSLFDKDGDGTITTKELGTVMRSLGQNPTEAELQDMINEVDADGDGTIDFPEFLTMMA
RKMKDTDSEEEIREAFRVFDKDGNGYISAAELRHVMTNLGEKLTDEEVDEMIREADIDGDGQVNYEEFVVMMTAKEF
PPPPPPPPPPPPPPPPPPPPPPPPPPPPPPGGSMVDSSRRKFNKTGKALRAIGRLSSLESGGIGTGFPFDPHYVEVL
GERMHYVDVGPRDGTPVLFLHGNPTSSYVWRNIIPHVAPTHRCIAPDLIGMGKSDKPDLGYFFDDHVRFMDAFIEAL
GLEEVVLVIHDWGSALGFHWAKRNPERVKGIAFMEFIRPIPTWDEWPEFARETFQAFRTTDVGRKLIIDQNVFIEGT
LPMGVVRPLTEVEMDHYREPFLNPVDREPLWRFPNELPIAGEPANIVALVEEYMDWLHQSPVPKLLFWGTPGVLIPP
AEAARLAKSLPNCKAVDIGPGENLLQEDNPDLIGSEIARWLSTLEISG

>ChemoB-CaM
MVSKGEELFTGVVPILVELDGDVNGHKFSVRGEGEGDATYGKLTLKFICTTGKLPVPWPTLVTTLSHGVQCFARYPD
HMKQHDFFKSAMPEGYVQERTIFFKDDGTYKTRAEVKFEGDTLVNRIELKGVDFKEDGNILGHKLEYNFNSHNIYIM
AVKQKNGIKVNFKIRHNVEDGSVQLADHYQQNTPIGDGPVLLPDSHYLSTQSKLSKDPNEKRDHMVLLEFRTAAGIT
GGTLPDQLTEEQIAEFKEAFSLFDKDGDGTITTKELGTVMRSLGQNPTEAELQDMINEVDADGDGTIDFPEFLTMMA
RKMKDTDSEEEIREAFRVFDKDGNGYISAAELRHVMTNLGEKLTDEEVDEMIREADIDGDGQVNYEEFVVMMTAKEF
PPPPPPPPPPPPPPPPPPPPPPPPPPPPPPGGSMVDSSRRKFNKTGKALRAIGRLSSLESGGIGTGFPFDPHYVEVL
GERMHYVDVGPRDGTPVLFLHGNPTSSYVWRNIIPHVAPTHRCIAPDLIGMGKSDKPDLGYFFDDHVRFMDAFIEAL
GLEEVVLVIHDWGSALGFHWAKRNPERVKGIAFMEFIRPIPTWDEWPEFARETFQAFRTTDVGRKLIIDQNVFIEGT
LPMGVVRPLTEVEMDHYREPFLNPVDREPLWRFPNELPIAGEPANIVALVEEYMDWLHQSPVPKLLFWGTPGVLIPP
AEAARLAKSLPNCKAVDIGPGENLLQEDNPDLIGSEIARWLSTLEISG

>ChemoC-CaM
MVSKGEELFTGVVPILVELDGDVNGHKFSVSGEGEGDATYGKLTLKFICTTGKLPVPWPTLVTTLSWGVQCFARYPD
HMKQHDFFKSAMPEGYVQERTIFFKDDGNYKTRAEVKFEGDTLVNRIELKGIDFKEDGNILGHKLEYNAIHGNVYIT
ADKQKNGIKANFGLNCNIEDGSVQLADHYQQNTPIGDGPVLLPDNHYLSTQSKLSKDPNEKRDHMVLLEFVTAAGIT
GGTLPDQLTEEQIAEFKEAFSLFDKDGDGTITTKELGTVMRSLGQNPTEAELQDMINEVDADGDGTIDFPEFLTMMA
RKMKDTDSEEEIREAFRVFDKDGNGYISAAELRHVMTNLGEKLTDEEVDEMIREADIDGDGQVNYEEFVVMMTAKEF
PPPPPPPPPPPPPPPPPPPPPPPPPPPPPPGGSMVDSSRRKFNKTGKALRAIGRLSSLESGGIGTGFPFDPHYVEVL
GERMHYVDVGPRDGTPVLFLHGNPTSSYVWRNIIPHVAPTHRCIAPDLIGMGKSDKPDLGYFFDDHVRFMDAFIEAL
GLEEVVLVIHDWGSALGFHWAKRNPERVKGIAFMEFIRPIPTWDEWPEFARETFQAFRTTDVGRKLIIDQNVFIEGT
LPMGVVRPLTEVEMDHYREPFLNPVDREPLWRFPNELPIAGEPANIVALVEEYMDWLHQSPVPKLLFWGTPGVLIPP
AEAARLAKSLPNCKAVDIGPGENLLQEDNPDLIGSEIARWLSTLEISG

>ChemoY-CaM
MVSKGEELFTGVVPILVELDGDVNGHKFSVSGEGEGDATYGKLTLKLICTTGKLPVPWPTLVTTLGYGLQCFARYPD
HMKQHDFFKSAMPEGYVQERTIFFKDDGNYKTRAEVKFEGDTLVNRIELKGIDFKEDGNILGHKLEYNYNSHNVYIT
ADKQKNGIKANFKIRHNIEDGGVQLADHYQQNTPIGDGPVLLPDNHYLSYQSKLSKDPNEKRDHMVLLEFVTAAGIT
GGTLPDQLTEEQIAEFKEAFSLFDKDGDGTITTKELGTVMRSLGQNPTEAELQDMINEVDADGDGTIDFPEFLTMMA
RKMKDTDSEEEIREAFRVFDKDGNGYISAAELRHVMTNLGEKLTDEEVDEMIREADIDGDGQVNYEEFVVMMTAKEF
PPPPPPPPPPPPPPPPPPPPPPPPPPPPPPGGSMVDSSRRKFNKTGKALRAIGRLSSLESGGIGTGFPFDPHYVEVL
GERMHYVDVGPRDGTPVLFLHGNPTSSYVWRNIIPHVAPTHRCIAPDLIGMGKSDKPDLGYFFDDHVRFMDAFIEAL
GLEEVVLVIHDWGSALGFHWAKRNPERVKGIAFMEFIRPIPTWDEWPEFARETFQAFRTTDVGRKLIIDQNVFIEGT
LPMGVVRPLTEVEMDHYREPFLNPVDREPLWRFPNELPIAGEPANIVALVEEYMDWLHQSPVPKLLFWGTPGVLIPP
AEAARLAKSLPNCKAVDIGPGENLLQEDNPDLIGSEIARWLSTLEISG

>ChemoR-CaM
MVSKGEELIKENMRMKVVMEGSVNGHQFKCTGEGEGNPYMGTQTMRIKVIEGGPLPFAFDILATSFMYGSRTFIKYP
KGIPDFFKQSFPEGFTWERVTRYEDGGVVTVMQDTSLEDGCLVYHVQVRGVNFPSNGPVMQKKTKGWEPNTEMMYPA
DGGLRGYTHMALKVDGGGHLSCSFVTTYRSKKTVGNIKMPGIHAVDHRLERLEESDNEMFVVQREHAVAKFAGLGGG

MDELYKGGTLPDQLTEEQIAEFKEAFSLFDKDGDGTITTKELGTVMRSLGQNPTEAELQDMINEVDADGDGTIDFPE
FLTMMARKMKDTDSEEEIREAFRVFDKDGNGYISAAELRHVMTNLGEKLTDEEVDEMIREADIDGDGQVNYEEFVVM
MTAKEFPPPPPPPPPPPPPPPPPPPPPPPPPPPPPPGGSMVDSSRRKFNKTGKALRAIGRLSSLESGGIGTGFPFDP
HYVEVLGERMHYVDVGPRDGTPVLFLHGNPTSSYVWRNIIPHVAPTHRCIAPDLIGMGKSDKPDLGYFFDDHVRFMD
AFIEALGLEEVVLVIHDWGSALGFHWAKRNPERVKGIAFMEFIRPIPTWDEWPEFARETFQAFRTTDVGRKLIIDQN
VFIEGTLPMGVVRPLTEVEMDHYREPFLNPVDREPLWRFPNELPIAGEPANIVALVEEYMDWLHQSPVPKLLFWGTP
GVLIPPAEAARLAKSLPNCKAVDIGPGENLLQEDNPDLIGSEIARWLSTLEISG

>ChemoL-CaM
MGLSGDQMGQIEKIFKVVYPVDDHHFKVILHYGTLVIDGVTPNMIDYFGRPYEGIAVFDGKKITVTGTLWNGNKIID
ERLINPDGSLLFRVTINGVTGWRLCERILAGGTGGSGGTGGSMVFTLEDFVGDWRQTAGYNLDQVLEQGGVSSLFQN
LGVSVTPIQRIVLSGENGLKIDIHVIIPYEVSKGEELFTGVVPILVELDGDVNGHKFSVSGEGEGDATYGKLTLKFI
CTTGKLPVPWPTLVTTLTYGVQCFSRYPDHMKQHDFFKSAMPEGYVQERTIFFKDDGNYKTRAEVKFEGDTLVNRIE
LKGIDFKEDGNILGHKLEYNYNSHNVYIMADKQKNGIKVNFKIRHNIEDGSVQLADHYQQNTPIGDGPVLLPDNHYL
STQSKLSKDPNEKRDHMVLLEFVTAAGITGGTLPDQLTEEQIAEFKEAFSLFDKDGDGTITTKELGTVMRSLGQNPT
EAELQDMINEVDADGDGTIDFPEFLTMMARKMKDTDSEEEIREAFRVFDKDGNGYISAAELRHVMTNLGEKLTDEEV
DEMIREADIDGDGQVNYEEFVVMMTAKEFPPPPPPPPPPPPPPPPPPPPPPPPPPPPPPGGSMVDSSRRKFNKTGKA
LRAIGRLSSLESGGIGTGFPFDPHYVEVLGERMHYVDVGPRDGTPVLFLHGNPTSSYVWRNIIPHVAPTHRCIAPDL
IGMGKSDKPDLGYFFDDHVRFMDAFIEALGLEEVVLVIHDWGSALGFHWAKRNPERVKGIAFMEFIRPIPTWDEWPE
FARETFQAFRTTDVGRKLIIDQNVFIEGTLPMGVVRPLTEVEMDHYREPFLNPVDREPLWRFPNELPIAGEPANIVA
LVEEYMDWLHQSPVPKLLFWGTPGVLIPPAEAARLAKSLPNCKAVDIGPGENLLQEDNPDLIGSEIARWLSTLEISG

EGFP, EBFP2, mCerulean3, Venus, mRuby2, cpNanoLuc
HaloTag7
Calmodulin
M13 peptide
Linker
Interface mutations (XFP$^{A206K}$, HT7$^{L271E}$, EBFP2$^{N39Y}$)

**ATP sensors**

>ChemoG-ATP
MVSKGEELFTGVVPILVELDGDVNGHKFSVSGEGEGDATYGKLTLKFICTTGKLPVPWPTLVTTLTYGVQCFSRYPD
HMKQHDFFKSAMPEGYVQERTIFFKDDGNYKTRAEVKFEGDTLVNRIELKGIDFKEDGNILGHKLEYNYNSHNVYIM
ADKQKNGIKVNFKIRHNIEDGSVQLADHYQQNTPIGDGPVLLPDNHYLSTQSKLSKDPNEKRDHMVLLEFVTAAGIT
GGGMKTVKVNITTPDGPVYDADIEMVSVRAESGDLGILPGHIPTKAPLKIGAVRLKKDGQTEMVAVSGGTVEVRPDH
VTINAQAAETAEGIDKERAEAARQRAQERLNSQSDDTDIRRAELALQRALNRLDVAGKANEFGGGIGTGFPFDPHYV
EVLGERMHYVDVGPRDGTPVLFLHGNPTSSYVWRNIIPHVAPTHRCIAPDLIGMGKSDKPDLGYFFDDHVRFMDAFI
EALGLEEVVLVIHDWGSALGFHWAKRNPERVKGIAFMEFIRPIPTWDEWPEFARETFQAFRTTDVGRKLIIDQNVFI
EGTLPMGVVRPLTEVEMDHYREPFLNPVDREPLWRFPNELPIAGEPANIVALVEEYMDWLHQSPVPKLLFWGTPGVL
IPPAEAARLAKSLPNCKAVDIGPGENLLQEDNPDLIGSEIARWLSTLEISG

>ChemoB-ATP
MVSKGEELFTGVVPILVELDGDVNGHKFSVRGEGEGDATYGKLTLKFICTTGKLPVPWPTLVTTLSHGVQCFARYPD
HMKQHDFFKSAMPEGYVQERTIFFKDDGTYKTRAEVKFEGDTLVNRIELKGVDFKEDGNILGHKLEYNFNSHNIYIM
AVKQKNGIKVNFKIRHNVEDGSVQLADHYQQNTPIGDGPVLLPDSHYLSTQSKLSKDPNEKRDHMVLLEFRTAAGIT
GGGMKTVKVNITTPDGPVYDADIEMVSVRAESGDLGILPGHIPTKAPLKIGAVRLKKDGQTEMVAVSGGTVEVRPDH
VTINAQAAETAEGIDKERAEAARQRAQERLNSQSDDTDIRRAELALQRALNRLDVAGKANEFGGGIGTGFPFDPHYV
EVLGERMHYVDVGPRDGTPVLFLHGNPTSSYVWRNIIPHVAPTHRCIAPDLIGMGKSDKPDLGYFFDDHVRFMDAFI
EALGLEEVVLVIHDWGSALGFHWAKRNPERVKGIAFMEFIRPIPTWDEWPEFARETFQAFRTTDVGRKLIIDQNVFI
EGTLPMGVVRPLTEVEMDHYREPFLNPVDREPLWRFPNELPIAGEPANIVALVEEYMDWLHQSPVPKLLFWGTPGVL
IPPAEAARLAKSLPNCKAVDIGPGENLLQEDNPDLIGSEIARWLSTLEISG

>ChemoR-ATP
MVSKGEELIKENMRMKVVMEGSVNGHQFKCTGEGEGNPYMGTQTMRIKVIEGGPLPFAFDILATSFMYGSRTFIKYP
KGIPDFFKQSFPEGFTWERVTRYEDGGVVTVMQDTSLEDGCLVYHVQVRGVNFPSNGPVMQKKTKGWEPNTEMMYPA
DGGLRGYTHMALKVDGGGHLSCSFVTTYRSKKTVGNIKMPGIHAVDHRLERLEESDNEMFVVQREHAVAKFAGLGGG
MDELYKGGGMKTVKVNITTPDGPVYDADIEMVSVRAESGDLGILPGHIPTKAPLKIGAVRLKKDGQTEMVAVSGGTV
EVRPDHVTINAQAAETAEGIDKERAEAARQRAQERLNSQSDDTDIRRAELALQRALNRLDVAGKANEFGGGIGTGFP
FDPHYVEVLGERMHYVDVGPRDGTPVLFLHGNPTSSYVWRNIIPHVAPTHRCIAPDLIGMGKSDKPDLGYFFDDHVR
FMDAFIEALGLEEVVLVIHDWGSALGFHWAKRNPERVKGIAFMEFIRPIPTWDEWPEFARETFQAFRTTDVGRKLII
DQNVFIEGTLPMGVVRPLTEVEMDHYREPFLNPVDREPLWRFPNELPIAGEPANIVALVEEYMDWLHQSPVPKLLFW
GTPGVLIPPAEAARLAKSLPNCKAVDIGPGENLLQEDNPDLIGSEIARWLSTLEISG

>ChemoL-ATP
MGLSGDQMGQIEKIFKVVYPVDDHHFKVILHYGTLVIDGVTPNMIDYFGRPYEGIAVFDGKKITVTGTLWNGNKIID
ERLINPDGSLLFRVTINGVTGWRLCERILAGGTGGSGGTGGSMVFTLEDFVGDWRQTAGYNLDQVLEQGGVSSLFQN
LGVSVTPIQRIVLSGENGLKIDIHVIIPYEVSKGEELFTGVVPILVELDGDVNGHKFSVSGEGEGDATYGKLTLKFI
CTTGKLPVPWPTLVTTLTYGVQCFSRYPDHMKQHDFFKSAMPEGYVQERTIFFKDDGNYKTRAEVKFEGDTLVNRIE
LKGIDFKEDGNILGHKLEYNYNSHNVYIMADKQKNGIKVNFKIRHNIEDGSVQLADHYQQNTPIGDGPVLLPDNHYL
STQSKLSKDPNEKRDHMVLLEFVTAAGITGGGMKTVKVNITTPDGPVYDADIEMVSVRAESGDLGILPGHIPTKAPL
KIGAVRLKKDGQTEMVAVSGGTVEVRPDHVTINAQAAETAEGIDKERAEAARQRAQERLNSQSDDTDIRRAELALQR
ALNRLDVAGKANEFGGGIGTGFPFDPHYVEVLGERMHYVDVGPRDGTPVLFLHGNPTSSYVWRNIIPHVAPTHRCIA
PDLIGMGKSDKPDLGYFFDDHVRFMDAFIEALGLEEVVLVIHDWGSALGFHWAKRNPERVKGIAFMEFIRPIPTWDE
WPEFARETFQAFRTTDVGRKLIIDQNVFIEGTLPMGVVRPLTEVEMDHYREPFLNPVDREPLWRFPNELPIAGEPAN
IVALVEEYMDWLHQSPVPKLLFWGTPGVLIPPAEAARLAKSLPNCKAVDIGPGENLLQEDNPDLIGSEIARWLSTLE
ISG

EGFP, EBFP2, mRuby2, cpNanoLuc
HaloTag7
$F_O$-$F_1$ ε subunit
Linker
Interface mutation (EGFP$^{A206K}$, HT7$^{L271E}$, EBFP2$^{N39Y}$)

**NAD$^{+}$ sensors**

>ChemoG-NAD
MVSKGEELFTGVVPILVELDGDVNGHKFSVSGEGEGDATYGKLTLKFICTTGKLPVPWPTLVTTLTYGVQCFSRYPD
HMKQHDFFKSAMPEGYVQERTIFFKDDGNYKTRAEVKFEGDTLVNRIELKGIDFKEDGNILGHKLEYNYNSHNVYIM
ADKQKNGIKVNFKIRHNIEDGSVQLADHYQQNTPIGDGPVLLPDNHYLSTQSKLSKDPNEKRDHMVLLEFVRAAGIT
GGTMTLEEARKRVNELRDLIRYHNYRYYVLADPEISDAEYDRLLRELKELEERFPELKSPDSPTLQVGARPLEATFR
PVRHPTRMYSLDNAFNLDELKAFEERIERALGRKGPFAYTVEHLVDGLSVNLYYEEGVLVYGATRGDGEVGEEVTQN
LLTIPTIPRRLKGVPERLEVRGEVYMPIEAFLRLNEELEERGERIFKNPRNAAAGSLRQKDPRITAKRGLRATFWAL
GLGLEEVEREGVATQFALLHWLKEKGFPVEHGYARAVGAEGVEAVYQDWLKKRRALPFEANGVAVKLDELALWRELG
YTARAPRFAIAYKFPSGGIGTGFPFDPHYVEVLGERMHYVDVGPRDGTPVLFLHGNPTSSYVWRNIIPHVAPTHRCI
APDLIGMGKSDKPDLGYFFDDHVRFMDAFIEALGLEEVVLVIHDWGSALGFHWAKRNPERVKGIAFMEFIRPIPTWD
EWPEFARETFQAFRTTDVGRKLIIDQNVFIEGTLPMGVVRPLTEVEMDHYREPFLNPVDREPLWRFPNELPIAGEPA
NIVALVEEYMDWLHQSPVPKLLFWGTPGVLIPPAEAARLAKSLPNCKAVDIGPGENLLQEDNPDLIGSEIARWLSTL
EISG

>ChemoB-NAD
MVSKGEELFTGVVPILVELDGDVNGHKFSVRGEGEGDATYGKLTLKFICTTGKLPVPWPTLVTTLSHGVQCFARYPD
HMKQHDFFKSAMPEGYVQERTIFFKDDGTYKTRAEVKFEGDTLVNRIELKGVDFKEDGNILGHKLEYNFNSHNIYIM
AVKQKNGIKVNFKIRHNVEDGSVQLADHYQQNTPIGDGPVLLPDSHYLSTQSKLSKDPNEKRDHMVLLEFRRAAGIT
GGTMTLEEARKRVNELRDLIRYHNYRYYVLADPEISDAEYDRLLRELKELEERFPELKSPDSPTLQVGARPLEATFR
PVRHPTRMYSLDNAFNLDELKAFEERIERALGRKGPFAYTVEHLVDGLSVNLYYEEGVLVYGATRGDGEVGEEVTQN
LLTIPTIPRRLKGVPERLEVRGEVYMPIEAFLRLNEELEERGERIFKNPRNAAAGSLRQKDPRITAKRGLRATFWAL
GLGLEEVEREGVATQFALLHWLKEKGFPVEHGYARAVGAEGVEAVYQDWLKKRRALPFEANGVAVKLDELALWRELG
YTARAPRFAIAYKFPSGGIGTGFPFDPHYVEVLGERMHYVDVGPRDGTPVLFLHGNPTSSYVWRNIIPHVAPTHRCI
APDLIGMGKSDKPDLGYFFDDHVRFMDAFIEALGLEEVVLVIHDWGSALGFHWAKRNPERVKGIAFMEFIRPIPTWD
EWPEFARETFQAFRTTDVGRKLIIDQNVFIEGTLPMGVVRPLTEVEMDHYREPFLNPVDREPLWRFPNELPIAGEPA
NIVALVEEYMDWLHQSPVPKLLFWGTPGVLIPPAEAARLAKSLPNCKAVDIGPGENLLQEDNPDLIGSEIARWLSTL
EISG

>ChemoR-NAD
MVSKGEELIKENMRMKVVMEGSVNGHQFKCTGEGEGNPYMGTQTMRIKVIEGGPLPFAFDILATSFMYGSRTFIKYP
KGIPDFFKQSFPEGFTWERVTRYEDGGVVTVMQDTSLEDGCLVYHVQVRGVNFPSNGPVMQKKTKGWEPNTEMMYPA
DGGLRGYTHMALKVDGGGHLSCSFVTTYRSKKTVGNIKMPGIHAVDHRLERLEESDNEMFVVQREHAVAKFAGLGGG
MDELYKGGTMTLEEARKRVNELRDLIRYHNYRYYVLADPEISDAEYDRLLRELKELEERFPELKSPDSPTLQVGARP
LEATFRPVRHPTRMYSLDNAFNLDELKAFEERIERALGRKGPFAYTVEHLVDGLSVNLYYEEGVLVYGATRGDGEVG
EEVTQNLLTIPTIPRRLKGVPERLEVRGEVYMPIEAFLRLNEELEERGERIFKNPRNAAAGSLRQKDPRITAKRGLR
ATFWALGLGLEEVEREGVATQFALLHWLKEKGFPVEHGYARAVGAEGVEAVYQDWLKKRRALPFEANGVAVKLDELA
LWRELGYTARAPRFAIAYKFPSGGIGTGFPFDPHYVEVLGERMHYVDVGPRDGTPVLFLHGNPTSSYVWRNIIPHVA
PTHRCIAPDLIGMGKSDKPDLGYFFDDHVRFMDAFIEALGLEEVVLVIHDWGSALGFHWAKRNPERVKGIAFMEFIR
PIPTWDEWPEFARETFQAFRTTDVGRKLIIDQNVFIEGTLPMGVVRPLTEVEMDHYREPFLNPVDREPLWRFPNELP
IAGEPANIVALVEEYMDWLHQSPVPKLLFWGTPGVLIPPAEAARLAKSLPNCKAVDIGPGENLLQEDNPDLIGSEIA
RWLSTLEISG

>ChemoD-NAD
MVSKGEELFTGVVPILVELDGDVNGHKFSVSGEGEGDATYGKLTLKLICTTGKLPVPWPTLVTTFGYGLMCFARYPD
HMKQHDFFKSAMPEGYVQERTIFFKDDGNYKTRAEVKFEGDTLVNRIELKGIDFKEDGNILGHKLEYNWNSHNVYIM
ADKQKNGIKVNFKIRHNIEDGSVQLADHYQQNTPIGDGPVLLPDNHYLSTQSKLSKDPNEKRDHMVLLEFVRAAGIT
GGTMTLEEARKRVNELRDLIRYHNYRYYVLADPEISDAEYDRLLRELKELEERFPELKSPDSPTLQVGARPLEATFR
PVRHPTRMYSLDNAFNLDELKAFEERIERALGRKGPFAYTVEHLVDGLSVNLYYEEGVLVYGATRGDGEVGEEVTQN
LLTIPTIPRRLKGVPERLEVRGEVYMPIEAFLRLNEELEERGERIFKNPRNAAAGSLRQKDPRITAKRGLRATFWAL
GLGLEEVEREGVATQFALLHWLKEKGFPVEHGYARAVGAEGVEAVYQDWLKKRRALPFEANGVAVKLDELALWRELG
YTARAPRFAIAYKFPSGGIGTGFPFDPHYVEVLGERMHYVDVGPRDGTPVLFLHGNPTSSYVWRNIIPHVAPTHRCI
APDLIGMGKSDKPDLGYFFDDHVRFMDAFIEALGLEEVVLVIHDWGSALGFHWAKRNPERVKGIAFMEFIRPIPTWD

EWPEFARETFQAFRTTDVGRKLIIDQNVFIEGTLWMGVVRPLTEVEMDHYREPFLNPVDREPLWRFPNELPIAGEPA
NIVALVEEYMDWLHQSPVPKLLFWGTPGVLIPPAEAARLAKSLPNCKAVDIGPGENLLQEDNPDLIGSEIARWLSTL
EISG

>ChemoL-NAD
MGLSGDQMGQIEKIFKVVYPVDDHHFKVILHYGTLVIDGVTPNMIDYFGRPYEGIAVFDGKKITVTGTLWNGNKIID
ERLINPDGSLLFRVTINGVTGWRLCERILAGGTGGSGGTGGSMVFTLEDFVGDWRQTAGYNLDQVLEQGGVSSLFQN
LGVSVTPIQRIVLSGENGLKIDIHVIIPYEVSKGEELFTGVVPILVELDGDVNGHKFSVSGEGEGDATYGKLTLKFI
CTTGKLPVPWPTLVTTLTYGVQCFSRYPDHMKQHDFFKSAMPEGYVQERTIFFKDDGNYKTRAEVKFEGDTLVNRIE
LKGIDFKEDGNILGHKLEYNYNSHNVYIMADKQKNGIKVNFKIRHNIEDGSVQLADHYQQNTPIGDGPVLLPDNHYL
STQSKLSKDPNEKRDHMVLLEFVRAAGITGGTMTLEEARKRVNELRDLIRYHNYRYYVLADPEISDAEYDRLLRELK
ELEERFPELKSPDSPTLQVGARPLEATFRPVRHPTRMYSLDNAFNLDELKAFEERIERALGRKGPFAYTVEHLVDGL
SVNLYYEEGVLVYGATRGDGEVGEEVTQNLLTIPTIPRRLKGVPERLEVRGEVYMPIEAFLRLNEELEERGERIFKN
PRNAAAGSLRQKDPRITAKRGLRATFWALGLGLEEVEREGVATQFALLHWLKEKGFPVEHGYARAVGAEGVEAVYQD
WLKKRRALPFEANGVAVKLDELALWRELGYTARAPRFAIAYKFPSGGIGTGFPFDPHYVEVLGERMHYVDVGPRDGT
PVLFLHGNPTSSYVWRNIIPHVAPTHRCIAPDLIGMGKSDKPDLGYFFDDHVRFMDAFIEALGLEEVVLVIHDWGSA
LGFHWAKRNPERVKGIAFMEFIRPIPTWDEWPEFARETFQAFRTTDVGRKLIIDQNVFIEGTLPMGVVRPLTEVEMD
HYREPFLNPVDREPLWRFPNELPIAGEPANIVALVEEYMDWLHQSPVPKLLFWGTPGVLIPPAEAARLAKSLPNCKA
VDIGPGENLLQEDNPDLIGSEIARWLSTLEISG

EGFP, EBFP2, mRuby2, ShadowG, cpNanoLuc
*tt*LigA
HaloTag7
Linker
Interface mutations (EGFP$^{A206K}$, EGFP$^{T225R}$, HT7$^{L271E}$)
Catalytic mutations *tt*LigA (K117L, D289N)
Affinity mutations *tt*LigA (Y226W, V292A)

HT7$^{P174W}$

**Purification sequences**

>Strep-tag®II + <u>enterokinase cleavage </u>sequence (N-terminal)

WSHPQFEKGA<u>DDDDK</u>VPH[…]      (pET-51b(+) plasmids)

>Poly-histidine tag sequence (C-terminal)

[…]APGFSSISAHHHHHHHHHH

>Poly-histidine tag + <u>TEV cleavage</u> sequence (N-terminal)

HHHHHHHHHH<u>ENLYFQG</u>GG[…]  (pET-51b(+) plasmids for crystallography)

**Localization sequences**

>Nuclear exit signal (NES) (N-terminal or C-terminal)

[…]LPPLERLTL                    (pCDNA5 plasmids)
LQNELALKLAGLDINKTGGS[…]         (pAAV plasmids)

>Nuclear localization sequence (NLS) (C-terminal, 3 copies)

[…]KSGLRSRADPKKKRKVDPKKKRKVDPKKKRKVGSTGSR

>Exterior plasma membrane localization sequence (IgKchL[…]PDGFR$_{tm}$) (N-terminal and C-terminal)

METDTLLLWVLLLWVPGSTGDYPYDVPDYA[…]EQKLISEEDLNAVGQDTQEVIVVPHSLPFKVVVISAILALVVLT
IISLIIILIMLWQKKPR

>Nuclear envelope (LaminB1) localization sequence (C-terminal)

[…]MATATPVPPRMGSRAGGPTTPLSPTRLSRLQEKEELRELNDRLAVYIDKVRSLETENSALQLQVTEREEVRGRE
LTGLKALYETELADARRALDDTARERAKLQIELGKCKAEHDQLLLNYAKKESDLNGAQIKLREYEAALNSKDAALAT
ALGDKKSLEGDLEDLKDQIAQLEASLAAAKKQLADETLLKVDLENRCQSLTEDLEFRKSMYEEEINETRRKHETRLV
EVDSGRQIEYEYKLAQALHEMREQHDAQVRLYKEELEQTYHAKLENARLSSEMNTSTVNSAREELMESRMRIESLSS
QLSNLQKESRACLERIQELEDLLAKEKDNSRRMLTDKEREMAEIRDQMQQQLNDYEQLLDVKLALDMEISAYRKLLE
GEEERLKLSPSPSSRVTVSRASSSRSVRTTRGKRKRVDVEESEASSSVSISHSASATGNVCIEEIDVDGKFIRLKNT
SEQDQPMGGWEMIRKIGDTSVSYKYTSRYVLKAGQTVTIWAANAGVTASPPTDLIWKNQNSWGTGEDVKVILKNSQG
EEVAQRSTVFKTTIPEEEEEEEEEAAGVVVEEELFHQQGTPRASNRSCAIM

>Mitochondrial localization sequence (Cox8) (N-terminal, 4 copies)

4x[MSVLTPLLLRGLTGSARRLPVPRAKIHSLSVLTPLLLRGLTGSARRLPVPRAKIHSL][…]

#### Supplementary Figures

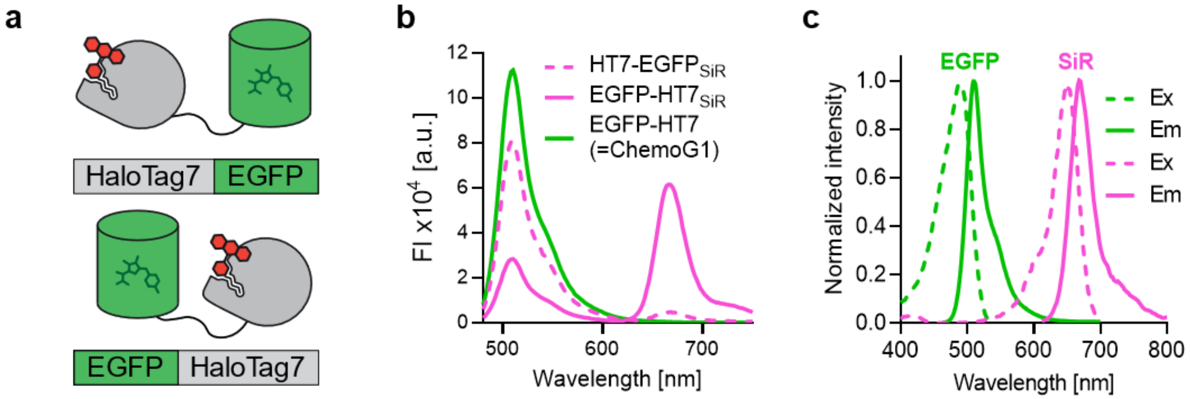

**Figure S 1| Initial design of the chemogenetic FRET pair.**

**a**. Schematic representation of the chemogenetic FRET pair based on EGFP and HaloTag7 (HT7) labeled with a synthetic rhodamine fluorophore. Shown are cartoons of the fusion of HT7 to the N- (HT7-EGFP) or C-terminus of EGFP (EGFP-HT7) **b**. Fluorescence intensity (FI) emission spectra of HT7-EGFP and EGFP-HT7 (= ChemoG1) labeled with SiR or not labeled. Represented are the means of 3 technical replicates. **c**. Normalized excitation (Ex) and emission (Em) spectra of EGFP and SiR. Represented are the means of 3 technical replicates.

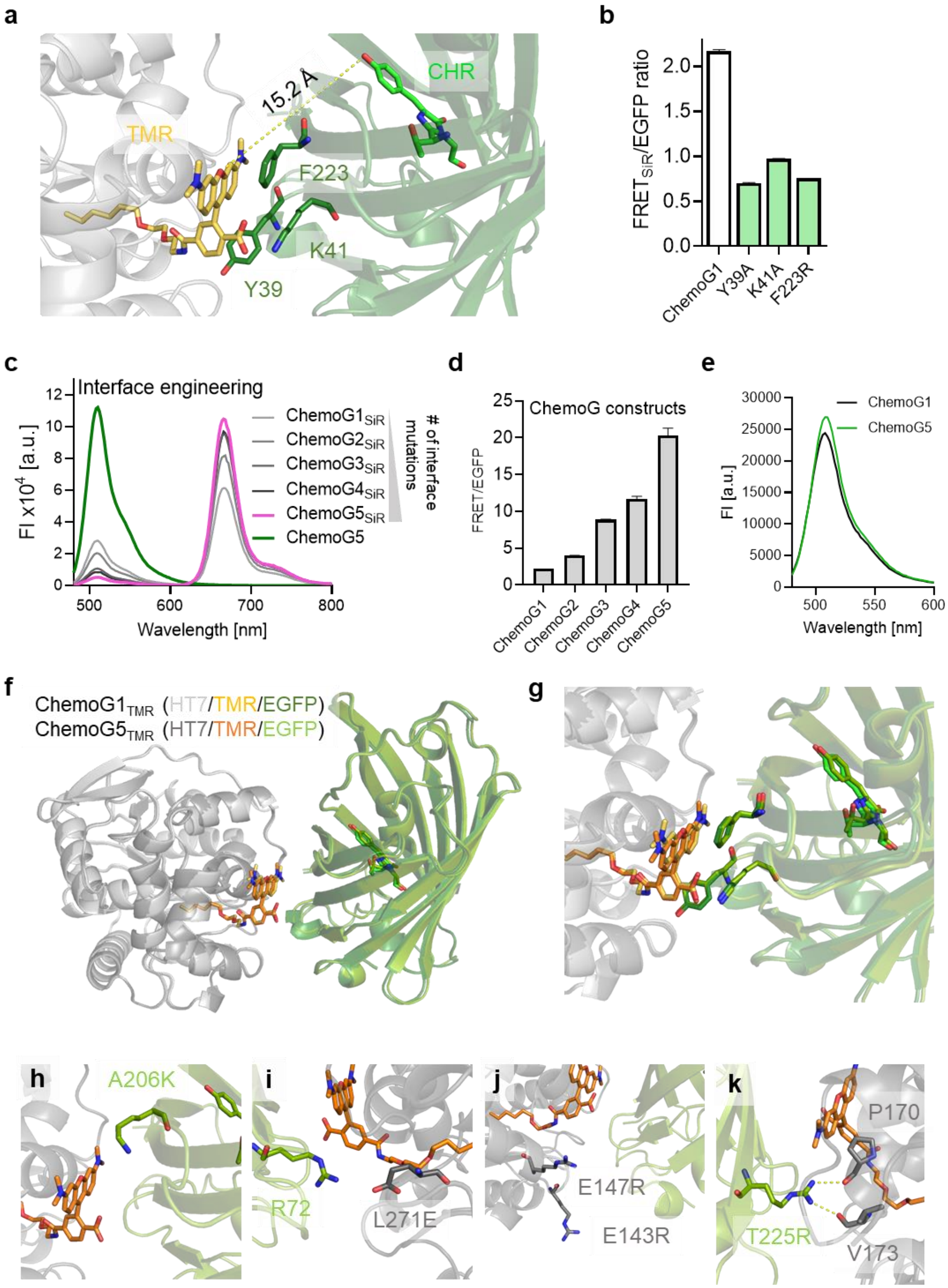

**Figure S 2| ChemoG-FRET surface engineering.**

**a**. X-ray structure of ChemoG1 labeled with TMR (PDB ID: 8B6S). Shown is the interface between EGFP and TMR labeled to HT. The EGFP chromophore and TMR are shown as sticks and their distance is marked with a dotted line. Residues Y39, K41 and F223 of EGFP involved in the direct interaction with TMR are annotated and shown as sticks.

**b**. FRET ratios of ChemoG1 and variants carrying mutations of residues involved in the EGFP/HT7$_{TMR}$ interface (n = 3 technical replicates, shown is the mean ±s.d.). **c**. Fluorescence intensity (FI) emission spectra of SiR-labeled ChemoG1-ChemoG5 and unlabeled ChemoG5. Represented are the means of 3 technical replicates. **d**. FRET ratios of SiR-labeled ChemoG1-ChemoG5 (n = 3 technical replicates, shown is the mean ±s.d.). **e**. Fluorescence intensity (FI) emission spectra of unlabeled ChemoG1 and ChemoG5. Represented are the means of 3 technical replicates. **f**, **g**. Structural comparison between ChemoG1$_{TMR}$ (PDB ID: 8B6S) and ChemoG5$_{TMR}$ (PDB ID: 8B6T).Structures represented as in **a**. Overview (**f**) and zoom-in (**g**) of the interface between EGFP and TMR are shown. **h**-**k**. Zoom-ins of the ChemoG5$_{TMR}$ X-ray structure showing the interface between EGFP and HT7 for each interface mutation (EGFP$^{A206K}$, HT7$^{L271E}$, HT7$^{E143R\text{-}E147R}$ and EGFP$^{T225R}$).

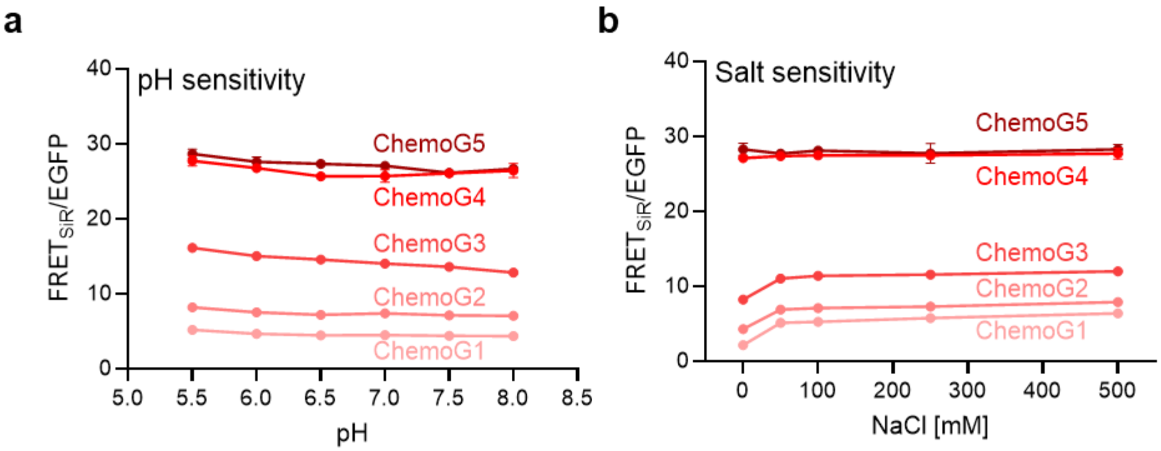

**Figure S 3| Sensitivity of ChemoG$_{SiR}$ to environmental changes.**
**a, b**. pH (**a**) and salt (**b**) sensitivity of the FRET/EGFP ratio of purified ChemoG constructs labeled with SiR. Shown are the means ±s.d. of 3 technical replicates.

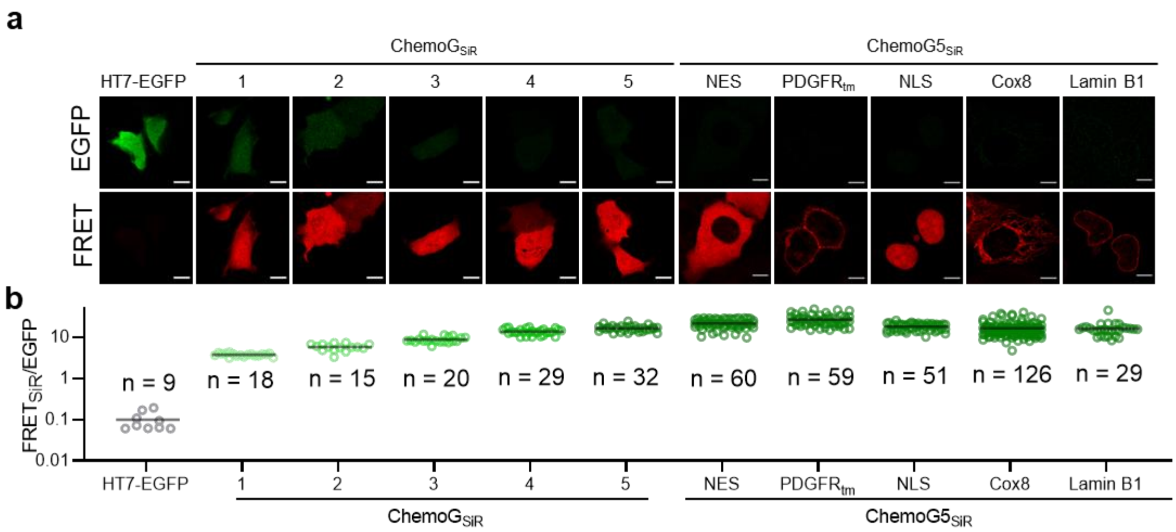

**Figure S 4| ChemoG performance in fluorescence microscopy.**
**a**. Confocal images of U-2 OS cells expressing untargeted HT7-EGFP, untargeted ChemoG1-5 or ChemoG5 targeted to different subcellular localizations. Cells were labeled with SiR. Shown are the EGFP and FRET channels. Scale bars = 10 µm. **b**. FRET/EGFP ratios of U-2 OS cells expressing different ChemoG constructs labeled with SiR as explained in **a**. Plotted for each construct are the FRET/EGFP ratios of individual cells (circles) and the mean (black line). The number of cells acquired for each construct are indicated and are derived from 2 independent experiments.

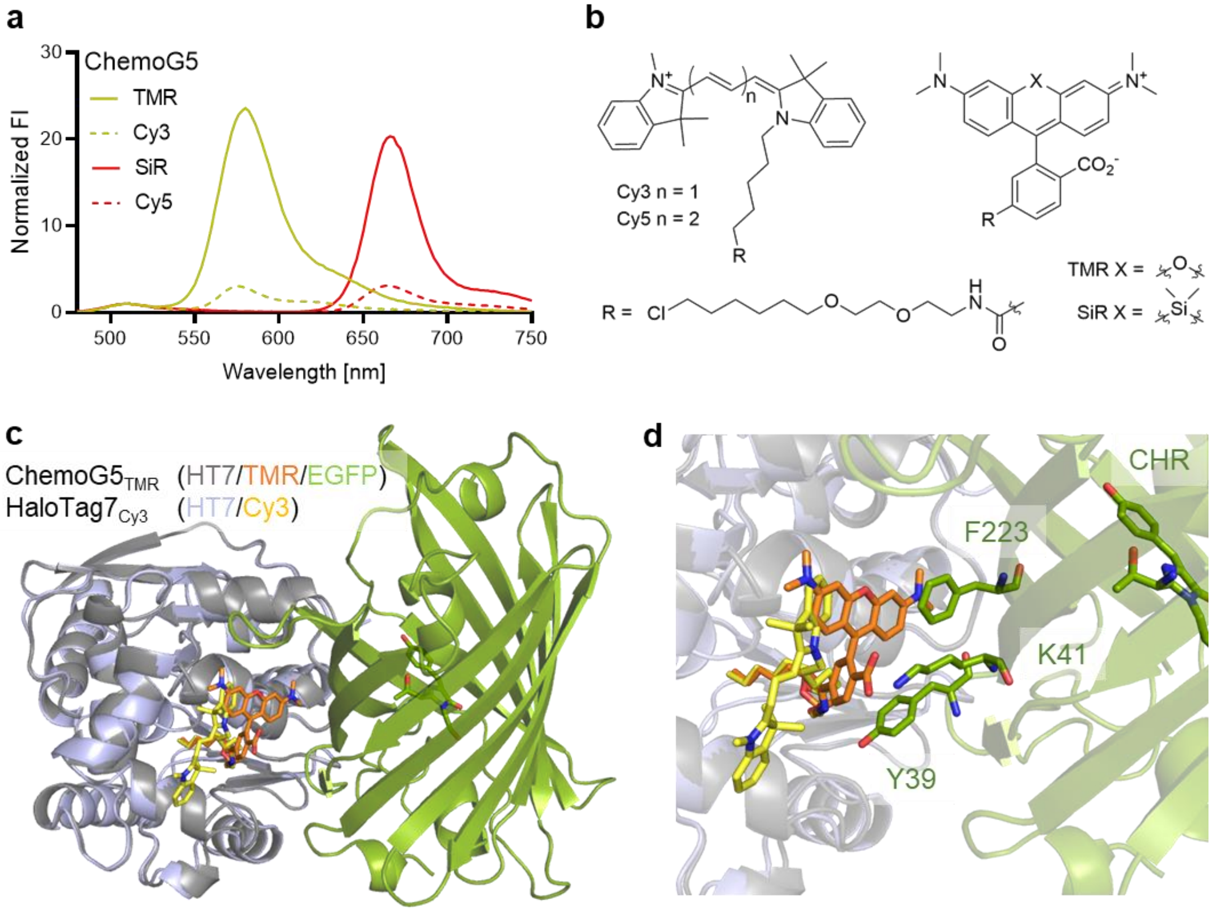

**Figure S 5| Impact of the fluorophore structure on the FRET efficiency of ChemoG5.**

**a**. Normalized fluorescence intensity (FI) emission spectra of ChemoG5 labeled with spectrally similar but structurally different fluorophores. Spectra were normalized to the maximum FI of EGFP. Shown are the means of 3 technical replicates. **b**. Chemical structures of cyanines (Cy3, Cy5) and rhodamines (TMR, SiR) coupled to the chloroalkane substrate (R) for HaloTag7. **c**. Structural comparison of the HaloTag7$_{Cy3}$ (PDB ID: 8B6R) and ChemoG5$_{TMR}$ (PDB ID: 8B6T) X-ray structures. HaloTag7$_{Cy3}$ was structurally aligned with the HaloTag7$_{TMR}$ component of ChemoG5$_{TMR}$. HaloTag7 (grey or light blue) and EGFP (green) are shown as cartoon. The EGFP chromophore (green), TMR (orange) and Cy3 (yellow) are shown as sticks. **d**. Zoom-on the interface HaloTag7/EGFP with representations as described in **c**. Residues Y39, K41 and F223 of EGFP involved in the direct interaction with TMR are annotated and shown as sticks.

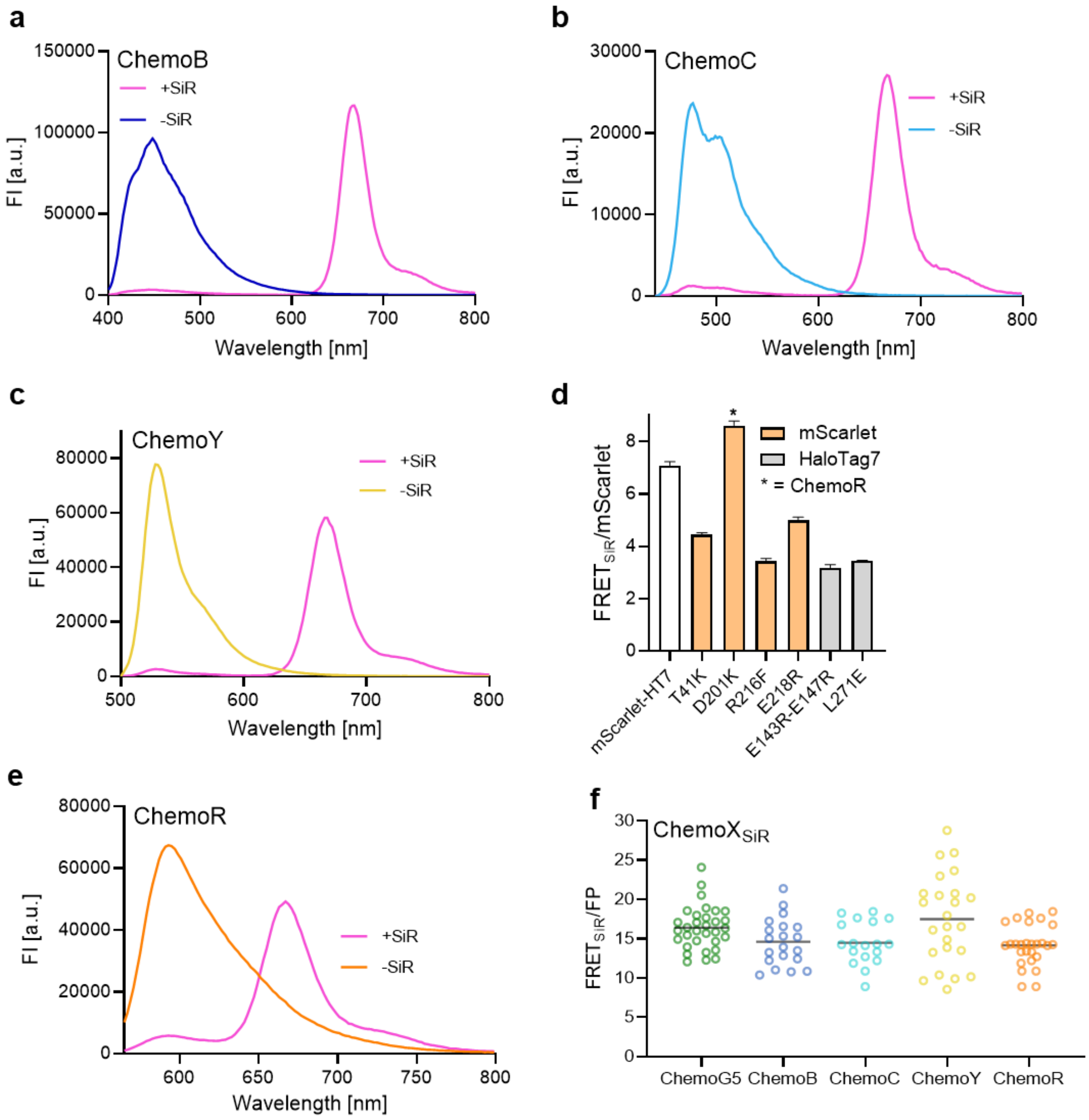

**Figure S 6| Expansion of ChemoG to other donor FPs.**
**a, c, e**. Fluorescence intensity (FI) emission spectra of optimized ChemoB (**a**), ChemoC (**b**), ChemoY (**c**) or ChemoR (**e**) labeled with SiR (+SiR) or unlabeled (-SiR). Interface mutations, FRET ratios and FRET efficiencies are listed in **Table S4**. Shown are the means of 3 technical replicates. **d**. FRET/mScarlet ratios of SiR-labeled mScarlet-HT7 and mScarlet-HT7 variants with different mutations on mScarlet (orange) or HT7 (grey). The mutation D201K used in the optimized ChemoR construct is marked with an asterisk. Shown are the means ±s.d. (n = 3 technical replicates) **f**. FRET ratios of ChemoX constructs expressed in U-2 OS cells and labeled with SiR (n > 17 cells). Plotted for each construct are the FRET/FP ratios of individual cells (circles) and the mean (black line). The values are derived from 2 independent experiments.

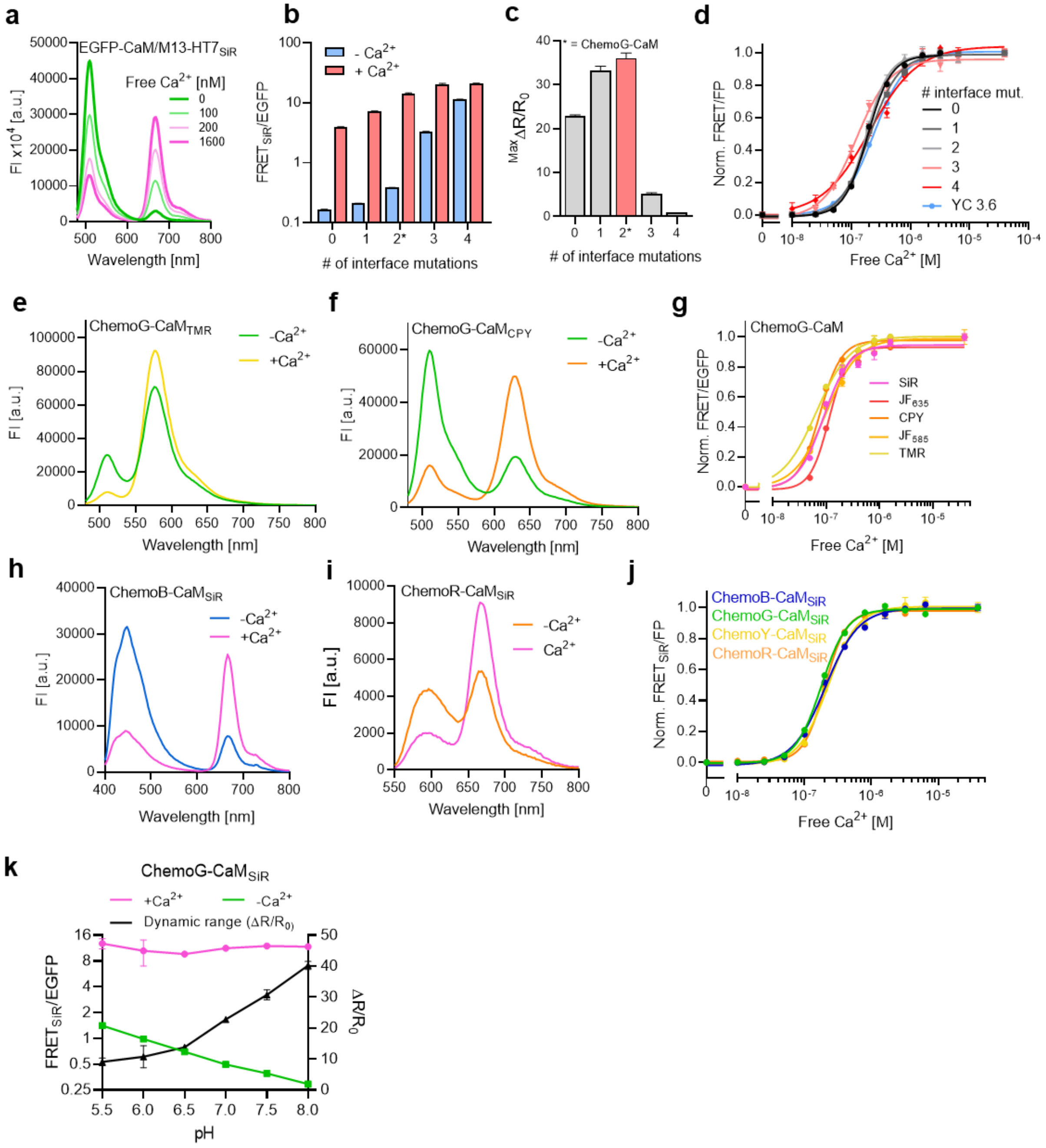

**Figure S 7| Engineering and characterization of ChemoX-CaM calcium sensors.**

**a**. Fluorescence intensity (FI) emission spectra of EGFP-CaM/M13-HT7 labeled with SiR in presence of different concentrations of free $Ca^{2+}$. Shown are the means of 3 technical replicates. **b**. FRET/EGFP ratios of calcium sensors differing in the number of interface mutations (**Table S5**) in presence (39 µM) or absence (0 µM) of free $Ca^{2+}$. Sensors were labeled with SiR. The variant corresponding to the final calcium sensor ChemoG-CaM is marked with an asterisk. Shown are the means ±s.d. n = 3 technical replicates. **c**. Maximal FRET/EGFP ratio changes ($^{Max}\Delta R/R_0$) of calcium sensors differing in the number of interface mutations (**Table S5**). Sensors were labeled with SiR. The variant corresponding to the final calcium sensor ChemoG-CaM is marked in red and with an asterisk. Shown are the means ±s.d. n = 3 technical replicates. **d**, **g**, **j**. $Ca^{2+}$ titrations of SiR-labeled calcium sensors differing in the number of interface mutations (**d**, **Table S5**), ChemoG-CaM labeled with different fluorophores (**g, Table S6**) or ChemoX-CaM calcium sensors labeled with SiR (**j, Table S5**). Shown are the means ±s.d. n = 3 technical replicates. **e, f**. Fluorescence intensity (FI) emission spectra of ChemoG-CaM labeled with TMR (**e**) or CPY (**f**) in presence (39 µM) or absence (0 µM) of free $Ca^{2+}$. Shown are the means of 3 technical replicates. **h**, **i**. Fluorescence intensity (FI) emission spectra of ChemoB-CaM (**h**) or ChemoR-CaM (**i**) labeled with SiR in presence (39 µM) or absence (0 µM) of free $Ca^{2+}$. Shown are the means of 3 technical replicates. **k**. pH sensitivity of the FRET/EGFP ratio of ChemoG-CaM$_{SiR}$ in presence (2 mM $CaCl_2$, +Ca2$^{+}$) or absence (2 mM EGTA, -Ca2$^{+}$) of free $Ca^{2+}$. Shown are the means ±s.d of three technical replicates.

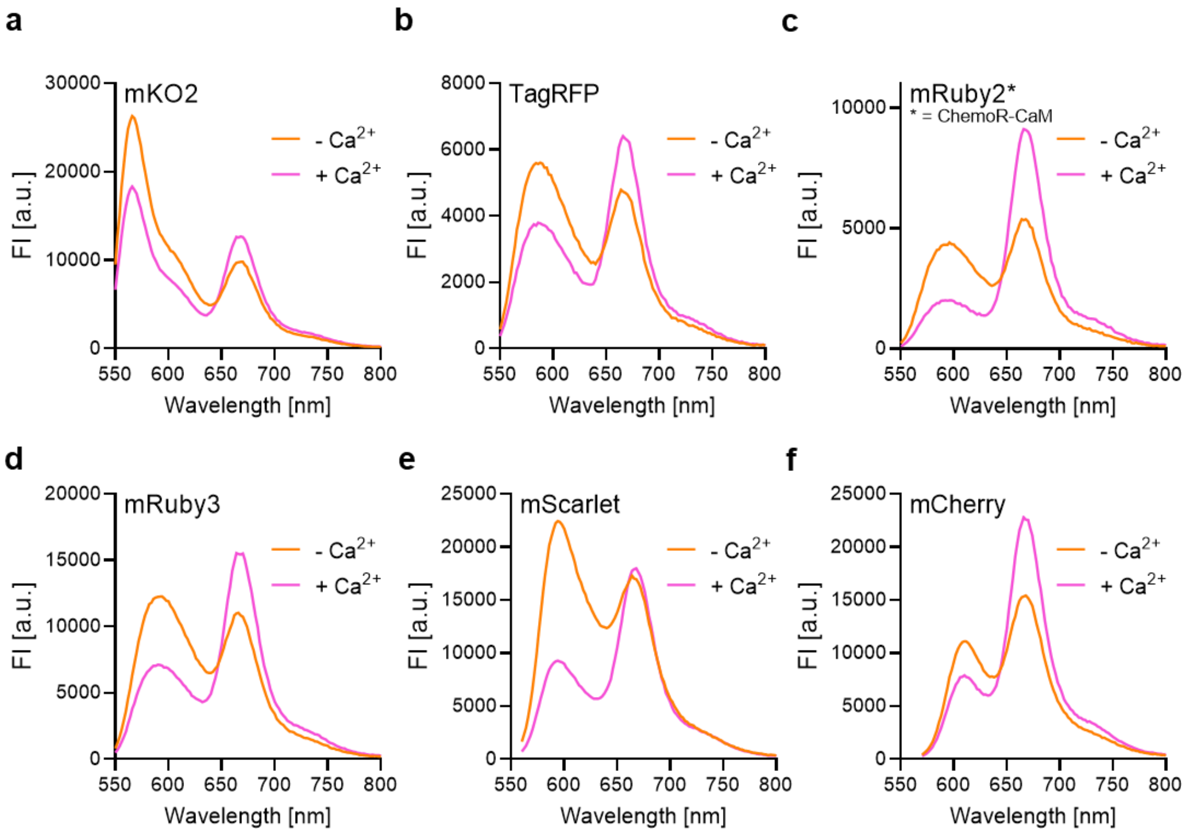

**Figure S 8| Implementation of different RFPs into the calcium sensor design.**
**a-f**. Fluorescence intensity (FI) emission spectra of RFP-CaM/M13-HT7 sensors labeled with SiR in abscence (2 mM EGTA, -$Ca^{2+}$) or presence (2 mM $CaCl_2$, +$Ca^{2+}$) of free $Ca^{2+}$. Different RFPs were used as FRET donor, indicated in the graph. Shown are the means of 3 technical replicates. *mRuby2 was chosen as the final ChemoR-CaM calcium sensor.

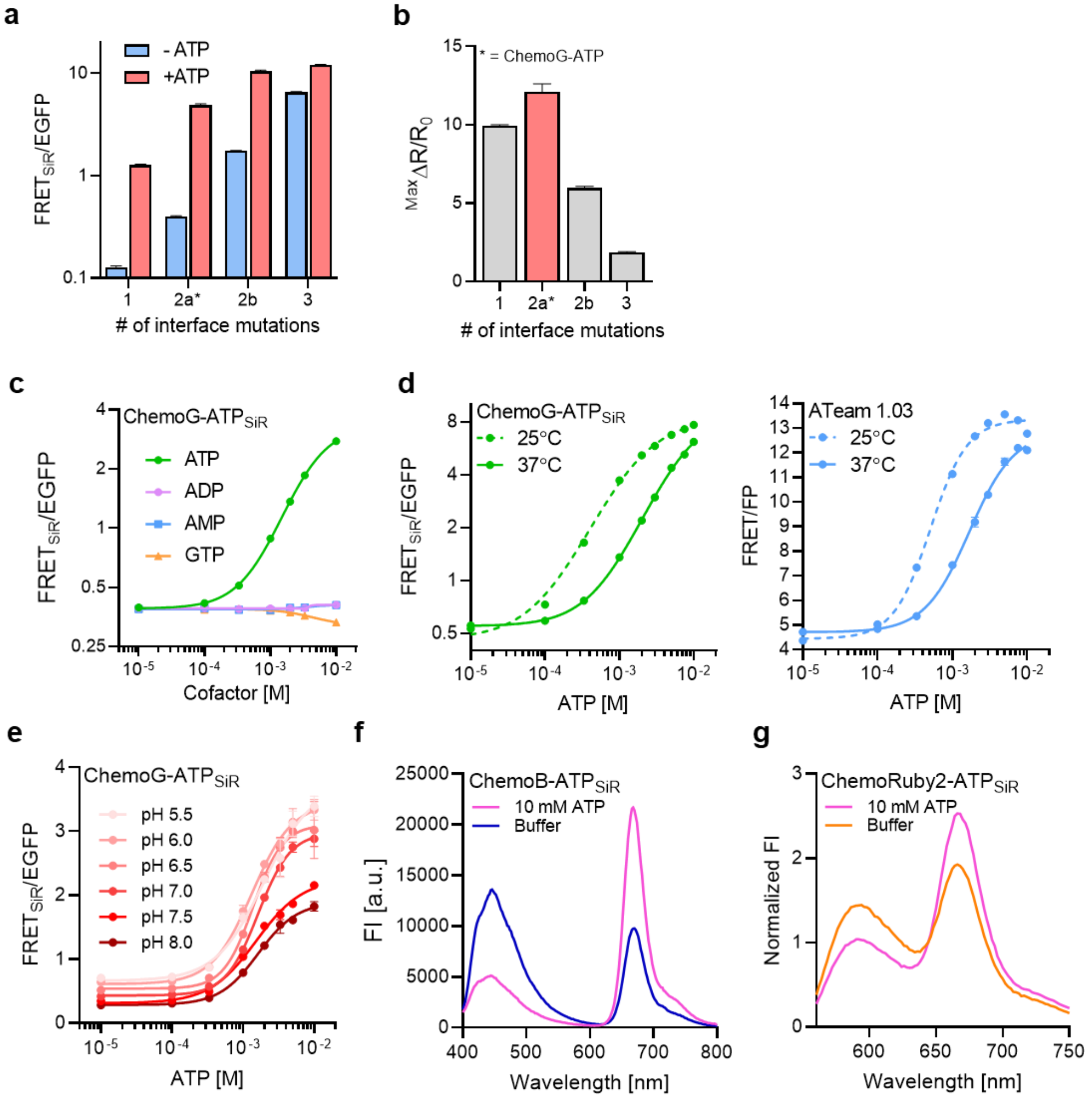

**Figure S 9| Engineering and characterization of ChemoX-ATP sensors.**

**a**. FRET/EGFP ratios of ATP sensors differing in the number of interface mutations (**Table S7**) in presence (+ATP) or absence (+ATP) of 10 mM ATP. Asterisk indicates construct corresponding to the final sensor ChemoG-ATP. Shown are the means ±s.d. n = 3 technical replicates. **b**. Maximal FRET/EGFP ratio changes ($^{Max}\Delta R/R_0$) of ATP sensors differing in the number of interface mutations (**Table S7**). Asterisk indicates construct corresponding to the final sensor ChemoG-ATP. Shown are the means ±s.d. n = 3 technical replicates. **c**. Titrations of ChemoG-ATP$_{SiR}$ with ATP and structurally related molecules. **d**. Titrations of ChemoG-ATP$_{SiR}$ and ATeam 1.03 with ATP at different temperatures. **e**. Titrations of ChemoG-ATP$_{SiR}$ and ATeam 1.03 with ATP at different pH. Shown are the means ±s.d. of 3 technical replicates. **f, g**. Fluorescence intensity (FI) emission spectra of ChemoB-ATP (**f**) and ChemoR-ATP (**g**) labeled with SiR (**Table S7**) in presence or absence of 10 mM ATP. Shown are the means of 3 technical replicates.

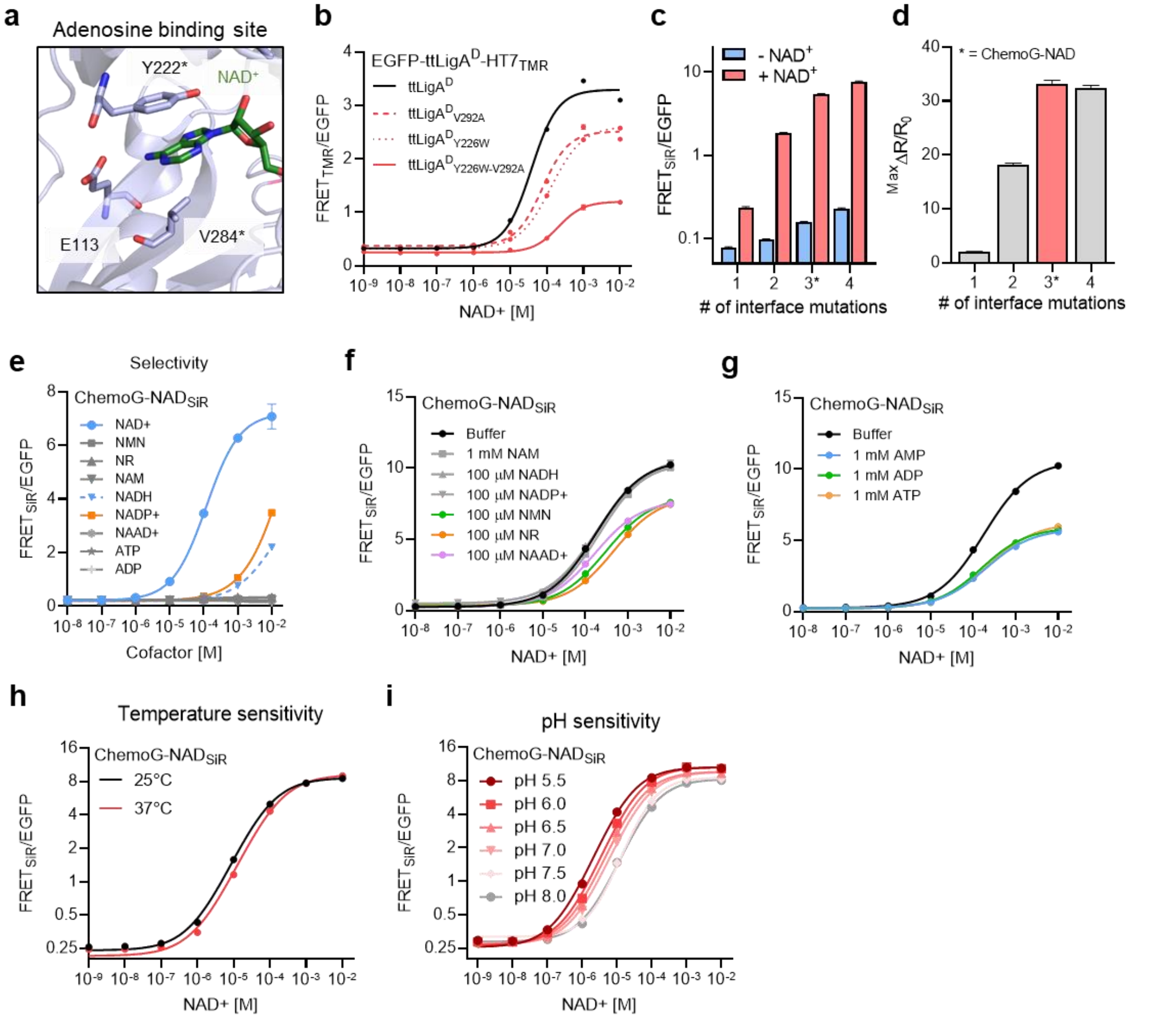

**Figure S 10| Engineering and characterization of ChemoX-NAD sensors.**

**a**. Zoom-in on the NAD$^+$ binding site of the X-ray structure of LigA from *Enterococcus faecalis (ef*LigA*)* bound to NAD$^+$ (PDB ID: 1TAE). The structure is represented as cartoon (light blue) and NAD$^+$ (green) and residues involved in the binding of NAD$^+$ (Y222 and V284, light blue) are represented as sticks. *Y222 and V284 of *ef*LigA correspond to Y226 and V292 of LigA from *Thermus thermophilus* (*tt*LigA). **b**. NAD$^+$ titrations of sensor variants labeled with TMR. *tt*LigA$^D$ carries the extra mutations K117L and D289N rendering it catalytically inactive. Mutations Y226W and Y292A shift the sensor response towards the range of free intracellular NAD$^+$. **c**. FRET ratios of NAD$^+$ sensors differing in the number of interface mutations (**Table S5**) in presence (+NAD$^+$) or absence (-NAD$^+$) of 1 mM NAD$^+$. **d**. Maximal FRET/EGFP ratio change ($^{Max}\Delta R/R_0$) of NAD+ sensors differing in the number of interface mutations. Asterisk indicates construct corresponding to the final sensor ChemoG-NAD. **e**. Titration of ChemoG-NAD with NAD$^+$ or structurally related molecules. **f**. Titration of ChemoG-NAD$_{SiR}$ with NAD+ in presence of different structurally related molecules. **g**. Titration of ChemoG-NAD$_{SiR}$ with NAD$^+$ in presence of 1 mM AMP, ADP or ATP. **h**, **i**. NAD$^+$ titrations of ChemoG-NAD$_{SiR}$ at different temperatures (**h**) or at different pH (**i**). For all graphs, the means of 3 technical replicates ±s.d. are shown.

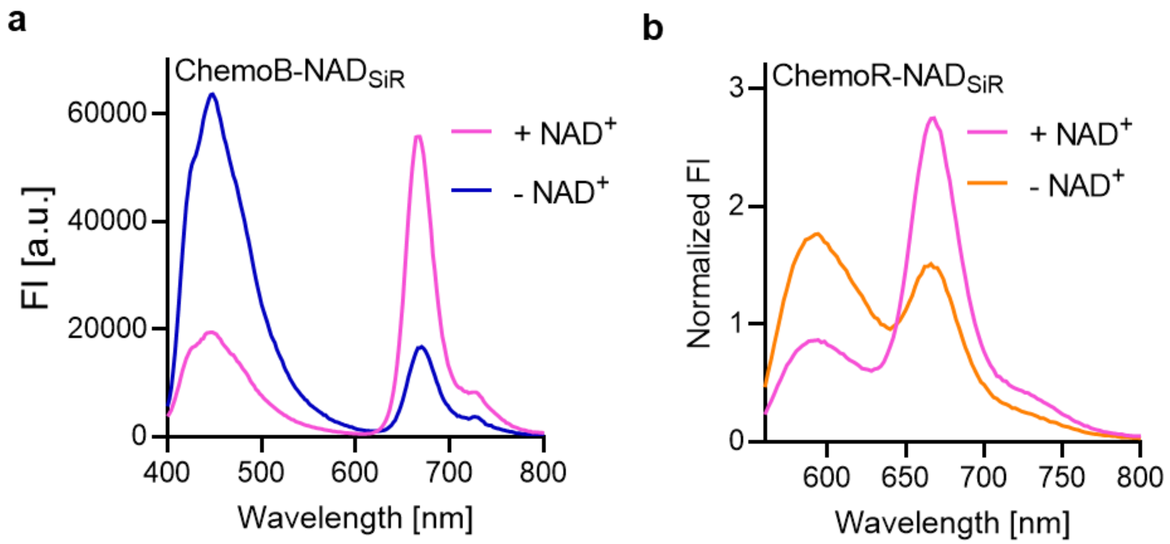

**Figure S 11| Emission spectra of ChemoB-NAD and ChemoR-NAD.**
**a**, **b**. Fluorescence intensity (FI) emission spectra of SiR-labeled ChemoB-NAD (**a**) and ChemoR-NAD (**b**) in presence or absence of 1 mM $NAD^+$. Shown are the means of 3 technical replicates.

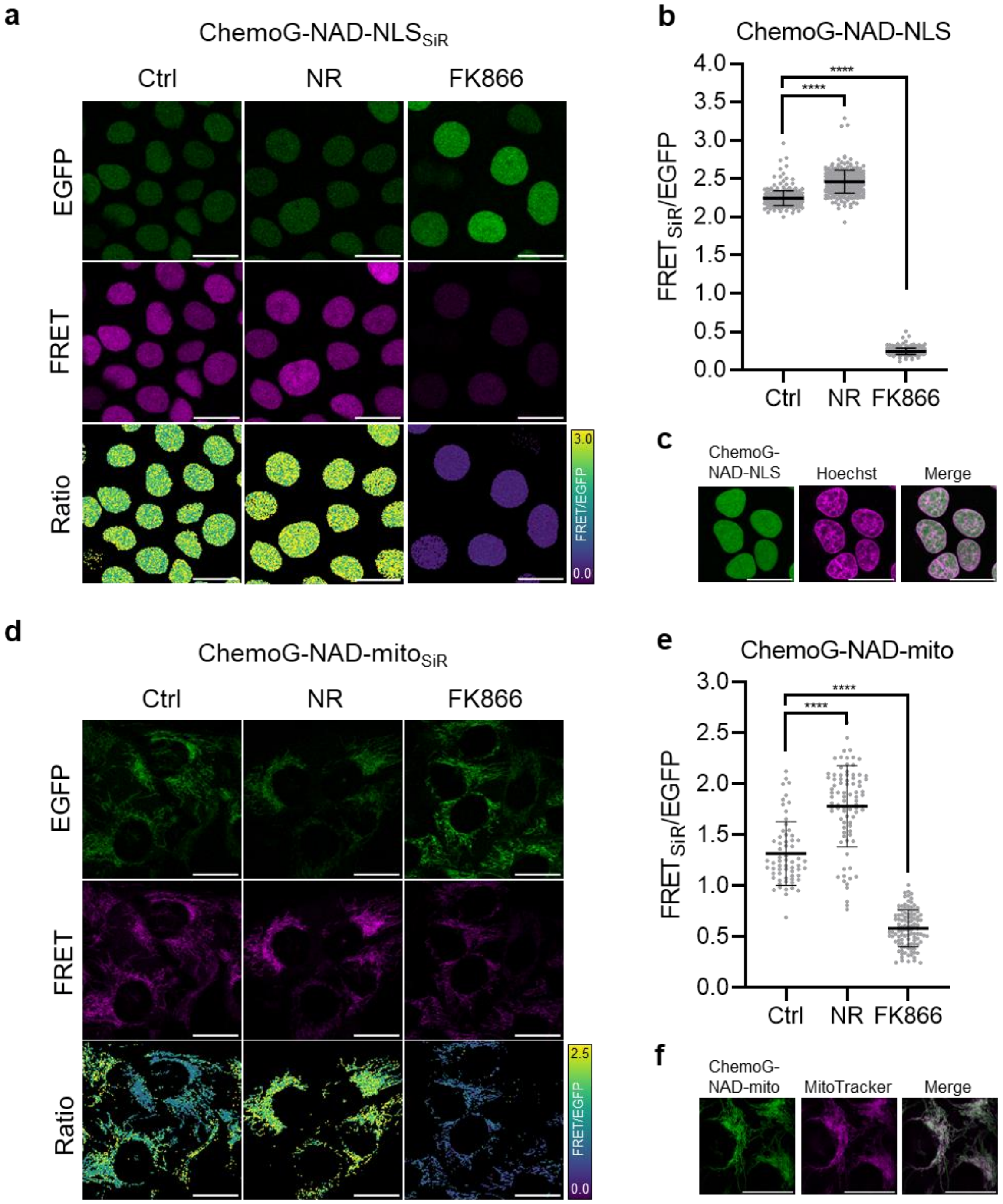

**Figure S 12| Performance of ChemoG-NAD in different subcellular compartments.**
**a**, **d**. Confocal images of U-2 OS cells expressing ChemoG-NAD in the nucleus (ChemoG-NAD-NLS, **a**) or mitochondria (ChemoG-NAD-mito, **d**) labeled with SiR. Shown are the EGFP channel, FRET channel and ratio image (FRET/EGFP) in pseudocolor (LUT = mpl-viridis). Cells were treated for 24 h either with DMSO (Ctrl), 100 nM FK866 or 1 mM NR. All scale bars = 25 µm. **b**, **e**. FRET/EGFP ratios of U-2 OS cells corresponding to panels **a** and **d**, respectively. Shown are the FRET/EGFP ratios of single cells (circles) and the mean ±s.d. (black line) (nucleus n ≥ 300 cells, mitochondria n ≥ 63 cells, from 3 independent experiments). p-values are given based on unpaired t-test with Welch's correction (**** p<0.0001). **c**. Confocal images of U-2 OS cells expressing ChemoG-NAD in the nucleus (ChemoG-NAD-NLS) and stained with Hoechst. **f**. Confocal images of U-2 OS cells expressing ChemoG-NAD in the mitochondria (ChemoG-NAD-mito) and stained with MitoTracker RedFM. All scale bars = 25 µm.

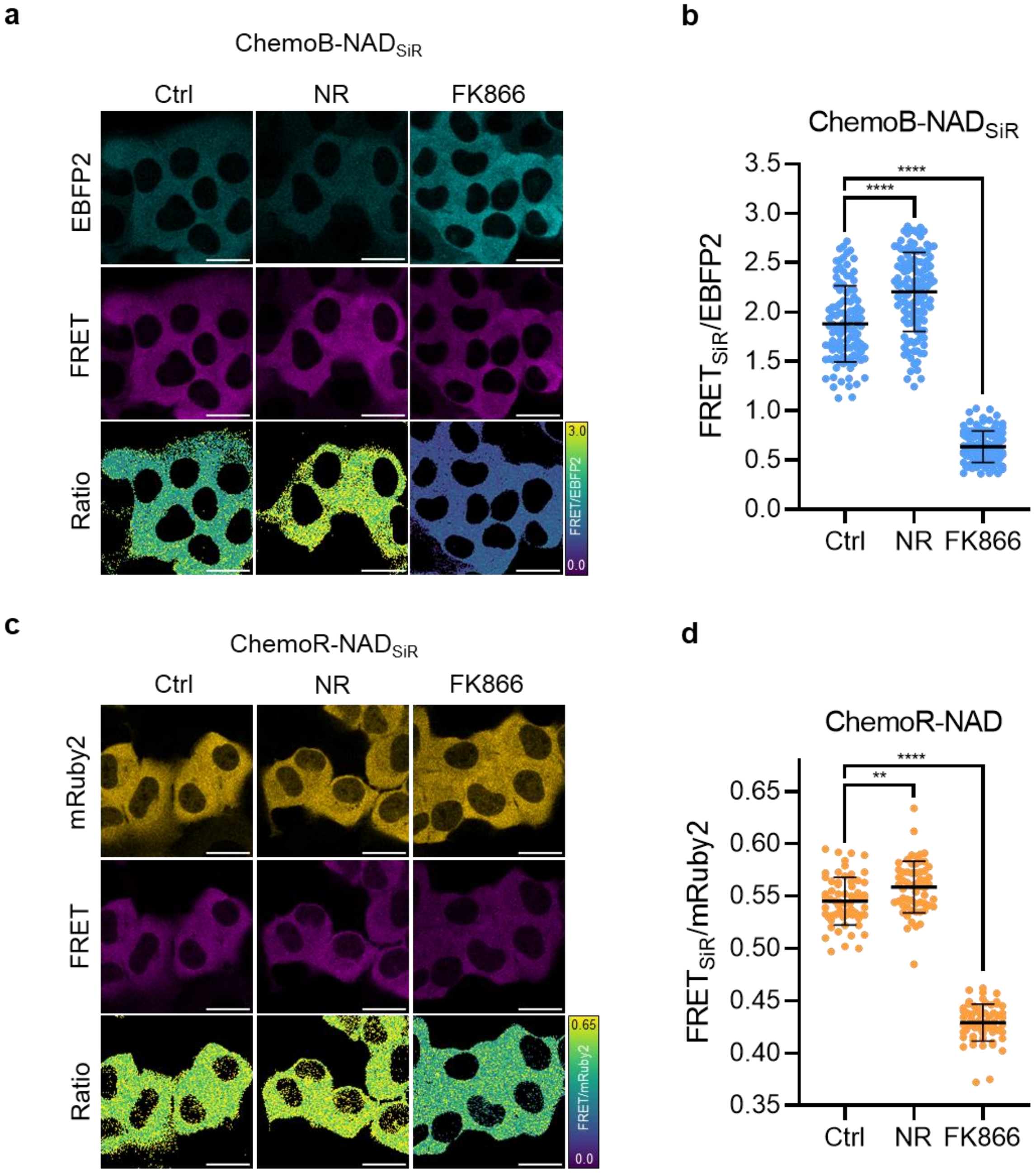

**Figure S 13| Performance of ChemoB-NAD and ChemoR-NAD in U-2 OS cells.**
**a**, **c**. Confocal images of U-2 OS cells expressing ChemoB-NAD (**a**) or ChemoR-NAD (**c**) in the cytosol labeled with SiR. Shown are the respective FP channel, FRET channel and ratio image (FRET/FP) in pseudocolor (LUT = mpl-viridis). Cells were treated for 24 h either with DMSO (Ctrl), 100 nM FK866 or 1 mM NR. All scale bars = 25 µm. **b**, **d**. FRET/FP ratios of U-2 OS cells corresponding to panels **a** and **c**, respectively. Shown are the FRET/FP values of single cells (circles) and the mean ±s.d. (black line) (n ≥ 109 cells for ChemoB-NAD, n ≥ 61 cells for ChemoR-NAD from 3 independent experiments). p-values are given based on unpaired t-test with Welch's correction (**** $p<0.0001$, ** $p<0.01$).

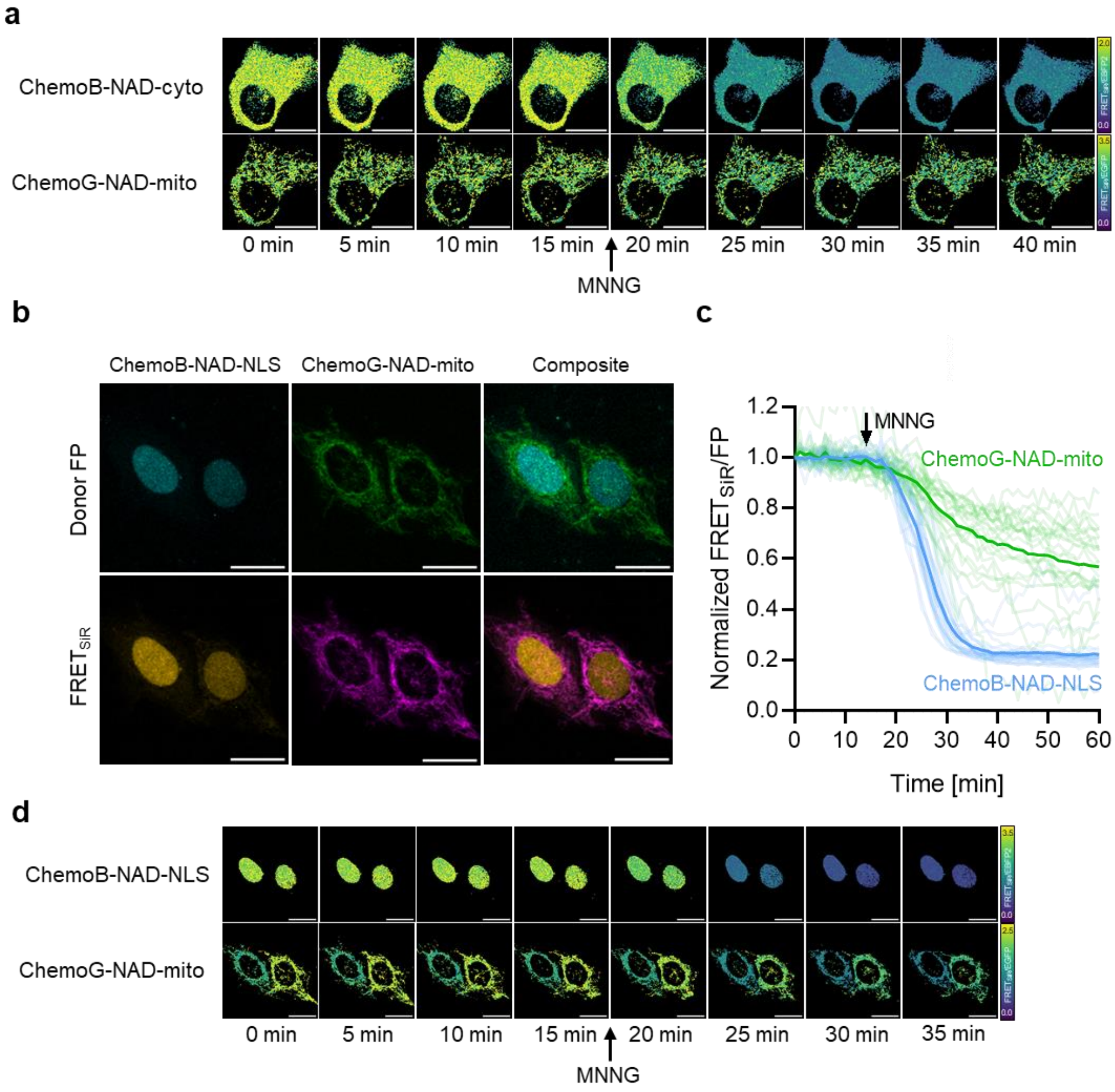

**Figure S 14| Multiplexed imaging of ChemoX-NAD sensors in U-2 OS cells.**
**a**. Confocal images of U-2 OS cells co-expressing ChemoB-NAD and ChemoG-NAD in the cytosol and mitochondria, respectively, labeled with SiR. Shown are the FRET/FP ratio images for each sensor in pseudocolor (LUT = mpl-viridis). Addition of 100 µM MNNG to the cells is indicated with an arrow (after t = 15 min). **b**. Confocal images of U-2 OS cells co-expressing ChemoB-NAD and ChemoG-NAD in the nucleus (NLS) and mitochondria, respectively. Shown are the FRET donor FP and FRET channels as well as the composite of FP or FRET channels of both sensors. **c**. Time course measurement of U-2 OS cells co-expressing ChemoB-NAD and ChemoG-NAD in the nucleus (NLS) and mitochondria, respectively, upon treatment with 100 µM MNNG. Shown are the means (line) and single cell traces (transparent lines) of the FRET/FP ratios of ChemoB-NAD and ChemoG-NAD normalized to 1 at t = 0 min (n = 25 cells from 2 biological replicates). **d**. Confocal images of U-2 OS cells co-expressing ChemoB-NAD and ChemoG-NAD in the nucleus (NLS) and mitochondria, respectively. Shown are the FRET/FP ratio images for each sensor in pseudocolor (LUT = mpl-viridis). Addition of 100 µM MNNG to the cells is indicated with an arrow (after t = 15 min).

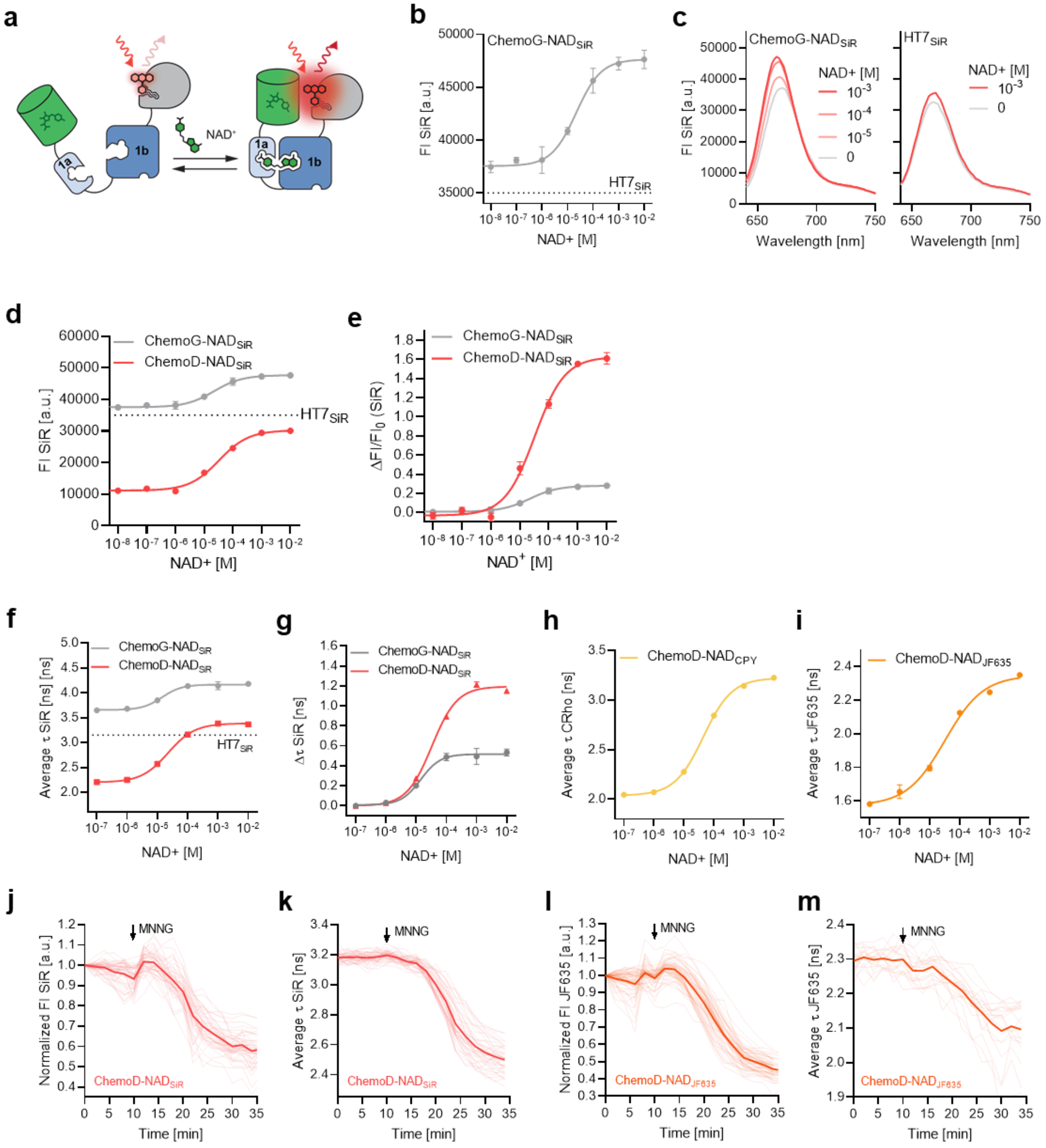

**Figure S 15| Engineering and characterization of intensiometric and fluorescence lifetime-based sensors.**
**a**. Schematic representation of intensiometric readout of ChemoG-NAD based on the labeled fluorophore. Closing of the sensor changes the environment of the rhodamine fluorophore and might thereby affect its photophysical properties. **b**. $NAD^+$ titration of intensiometric ChemoG-$NAD_{SiR}$ biosensor. Plotted are the fluorescence intensities (FI) of directly excited SiR labeled to ChemoG-NAD. The fluorescence intensity of HT7 labeled with SiR in presence of 1 mM $NAD^+$ is indicated with a dotted line. **c**. Fluorescence intensity (FI) emission spectra of SiR-labeled ChemoG-NAD (*left*) or HT7 (*right*) at different NAD+ concentrations. Shown are the means of 3 technical replicates. **d, e**. NAD+ titrations of intensiometric $NAD^+$ biosensors labeled with SiR. Plotted are the fluorescence intensities (FI) of directly excited SiR (**d**) or the change in fluorescence intensity of SiR ($\Delta FI/FI_0$, **e**). The fluorescence intensity of HT7 labeled with SiR in presence of 1 mM $NAD^+$ is indicated with a dotted line. **f**-**i**. $NAD^+$ titrations of fluorescence lifetime-based $NAD^+$ biosensors labeled with SiR (**f**, **g**), CPY (**h**) or $JF_{635}$ (**i**). Plotted are the intensity-weighted average fluorescence lifetimes ($\tau$, **f**, **h**, **i**) or the change in $\tau$ ($\Delta\tau$, **g**) of the directly excited fluorophores. The fluorescence lifetime of HT7 labeled with SiR in presence of 1 mM $NAD^+$ is indicated with a dotted line in **f**. **j-m**. Time course measurements of ChemoD-NAD

fluorescence intensity (FI) (**j, l**) or intensity-weighted average fluorescence lifetime (**k, m**) in U-2 OS cells labeled either with SiR (**j, k**) or $JF_{635}$ (**l, m**). FI was normalized to 1 at t = 0 min. Represented are the means (solid line) plus traces of single cells (transparent lines). Addition of 100 μM MNNG is indicated with an arrow. N = 44 cells from two independent experiments (**j**), n = 37 cells from two independent experiments (**k**), n = 53 cells from 2 independent experiments (**l**), n = 19 cells from one experiment (**m**). For all titrations, the mean ±s.d. from 3 technical replicates are represented.

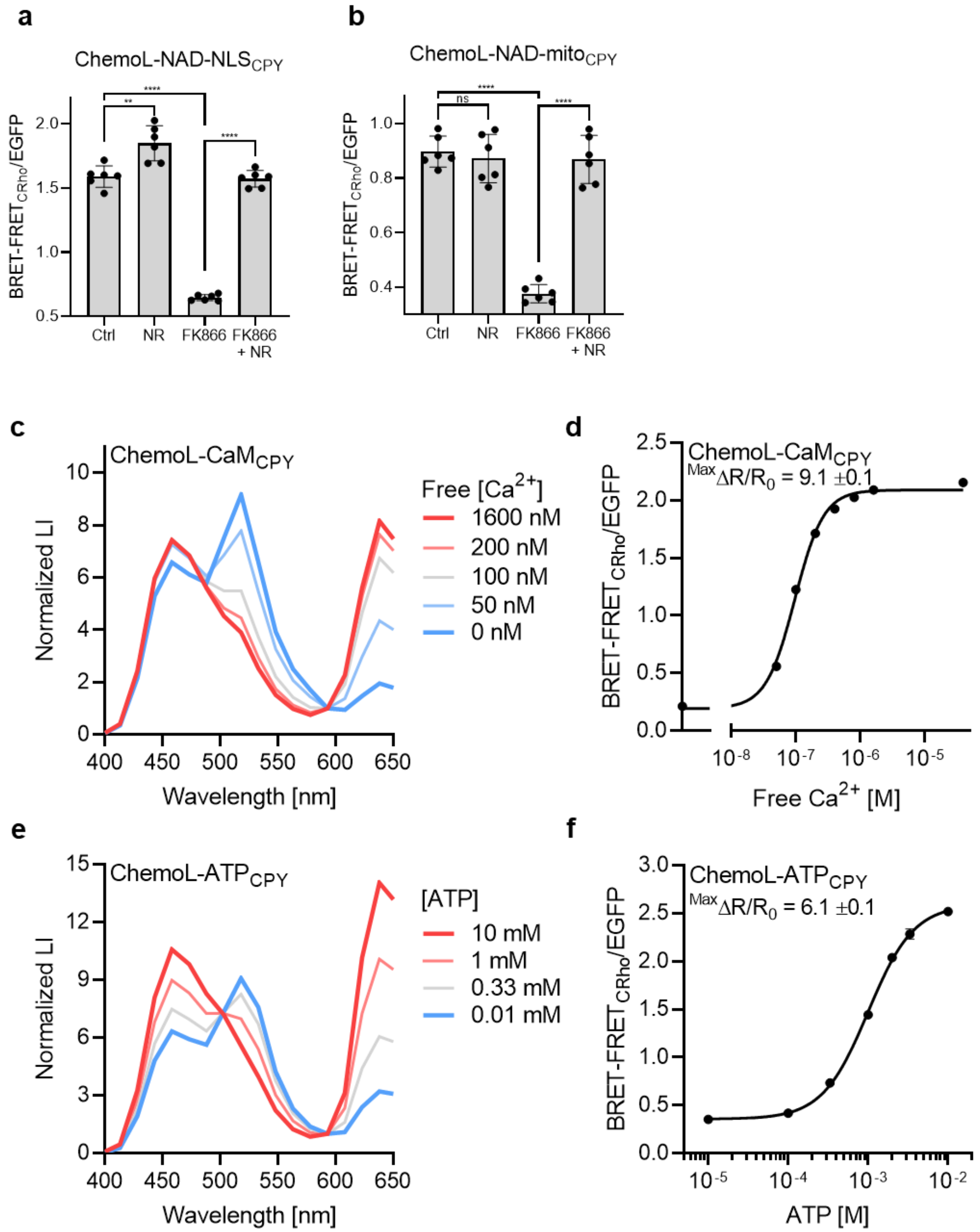

**Figure S 16| Engineering and characterization of ChemoL sensors.**
**a, b**. BRET-FRET/EGFP ratios of ChemoL-NAD$_{CPY}$ expressed in the nucleus (**a**) or the mitochondria (**b**) of U-2 OS cells upon treatment for 24 h with DMSO (Ctrl), 1 mM NR, 100 nM FK866 or 100 nM FK866 plus 1 mM NR. Represented are the BRET-FRET/EGFP ratios of single wells (circle) and means ±s.d. (black line) (n = 6 wells from a single experiment). p-values are given based on unpaired t-test with Welch's correction (**** p<0.0001, ** p<0.01, ns p>0.05). **c**, **e**. Luminescent intensity (LI) spectra of ChemoL-CaM (**c**) or ChemoL-ATP (**e**) at different concentrations of calcium and ATP, respectively. The spectra were normalized to the isosbestic point at 593 nm. Constructs were labeled with CPY. Shown are the means of 3 technical replicates. **d**, **f**. Analyte titrations of ChemoL-CaM (**d**) and ChemoL-ATP (**f**) labelled with CPY. Shown are the mean ±s.d. of BRET-FRET/EGFP ratios (n = 3 technical replicates). Indicated are also the maximum BRET-FRET/EGFP ratio changes ($^{Max}\Delta R/R_0$).

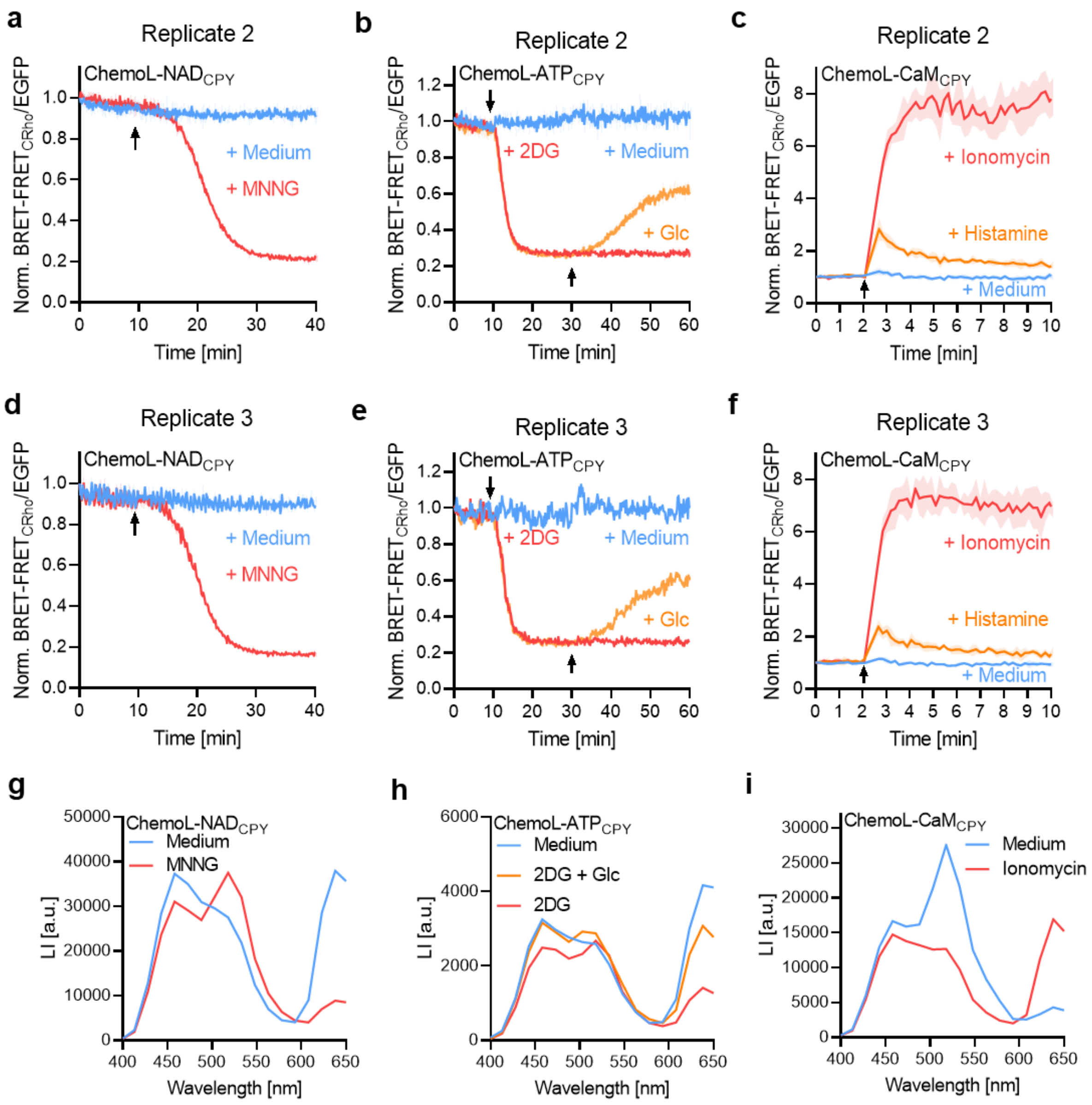

**Figure S 17| ChemoL sensor performances in U-2 OS cells.**

**a-f.** Time course measurements of ChemoL sensors expressed in U-2 OS cells (ChemoL-NAD, **a, d**) or HeLa Kyoto cells (ChemoL-ATP (**b, e**) and ChemoL-CaM (**c, f**)) upon drug treatments. Sensors were labeled with CPY. Represented are the BRET-FRET/EGFP ratios normalized to 1 at t = 0 min. Cells were untreated (+ medium) or treated with different reagents indicated with an arrow (n = 3 wells for each condition of each experiment). Represented are the mean (solid line) and the standard deviation (shade areas). The treatments are identical to time courses in **Fig. 6e-g**. **g-i**. Luminescent intensity (LI) spectra of ChemoL-NAD (**g**), ChemoL-ATP (**h**) or ChemoL-CaM (**i**) expressed in U-2 OS cells (ChemoL-NAD) or HeLa Kyoto cells (ChemoL-ATP and ChemoL-CaM). Sensors were labeled with CPY. The treatments are identical to time courses in **Fig. 6e-g**. Spectra were acquired immediately after the duration of the time courses (ChemoL-NAD = 40 min, ChemoL-ATP = 60 min, ChemoL-CaM = 10 min).

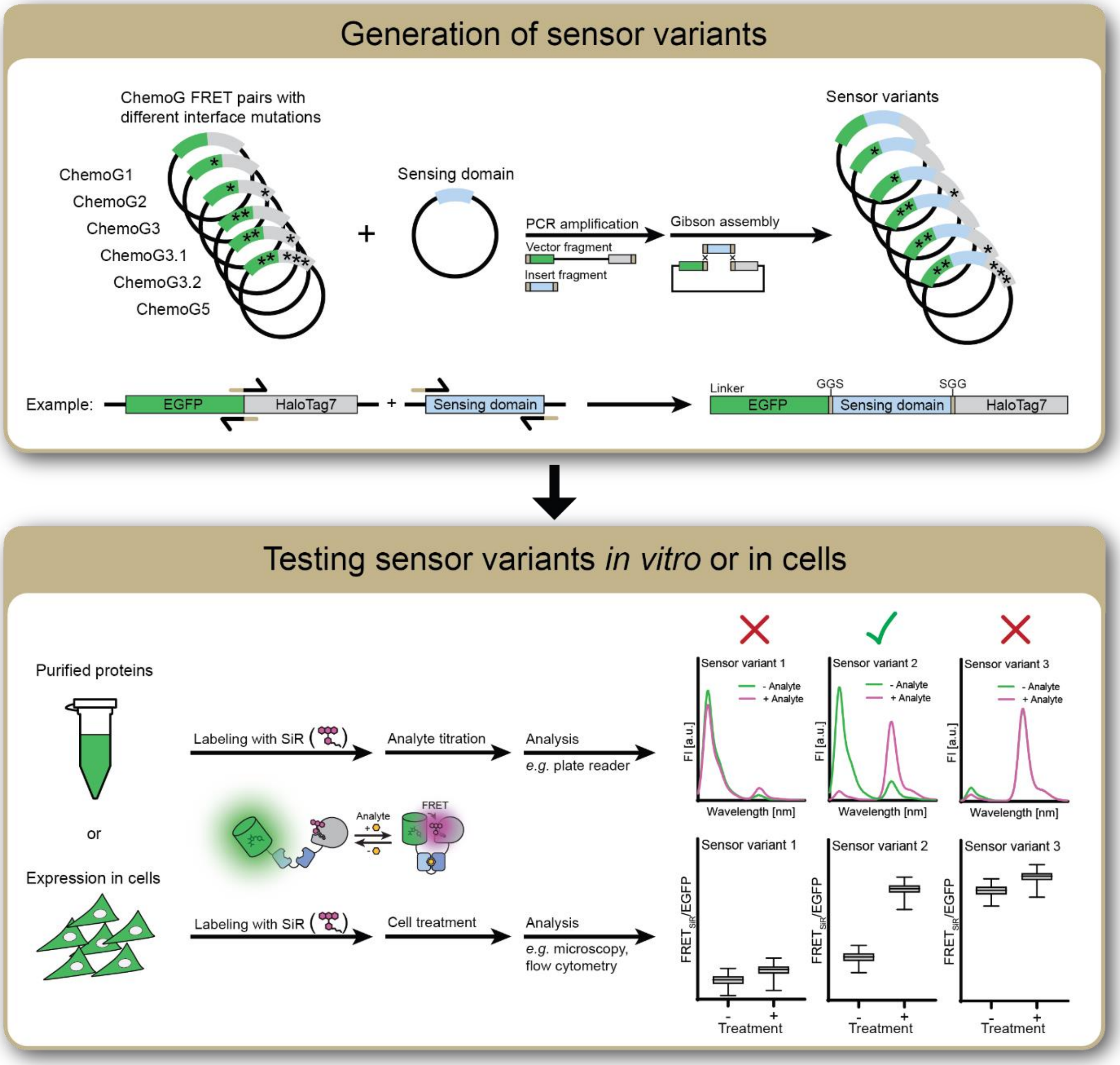

**Figure S 18| Development of ChemoG biosensors.**
See **Extended Note 1** for explanations.

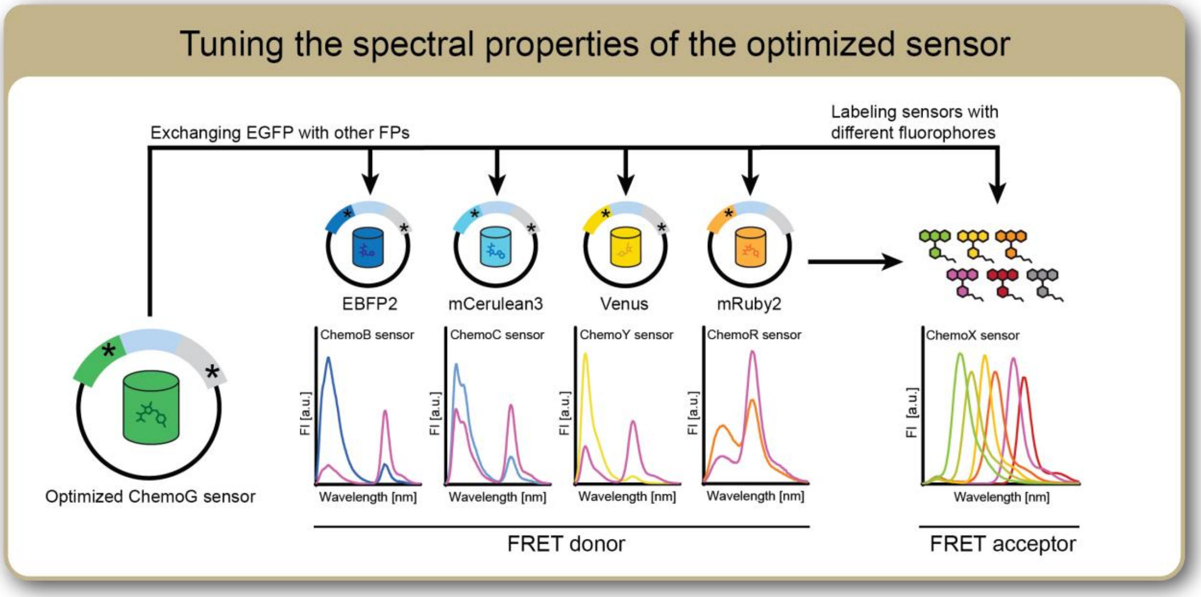

**Figure S 19| Tuning the spectral properties of the optimized ChemoG sensor.**
See **Extended Note 2** for explanations.

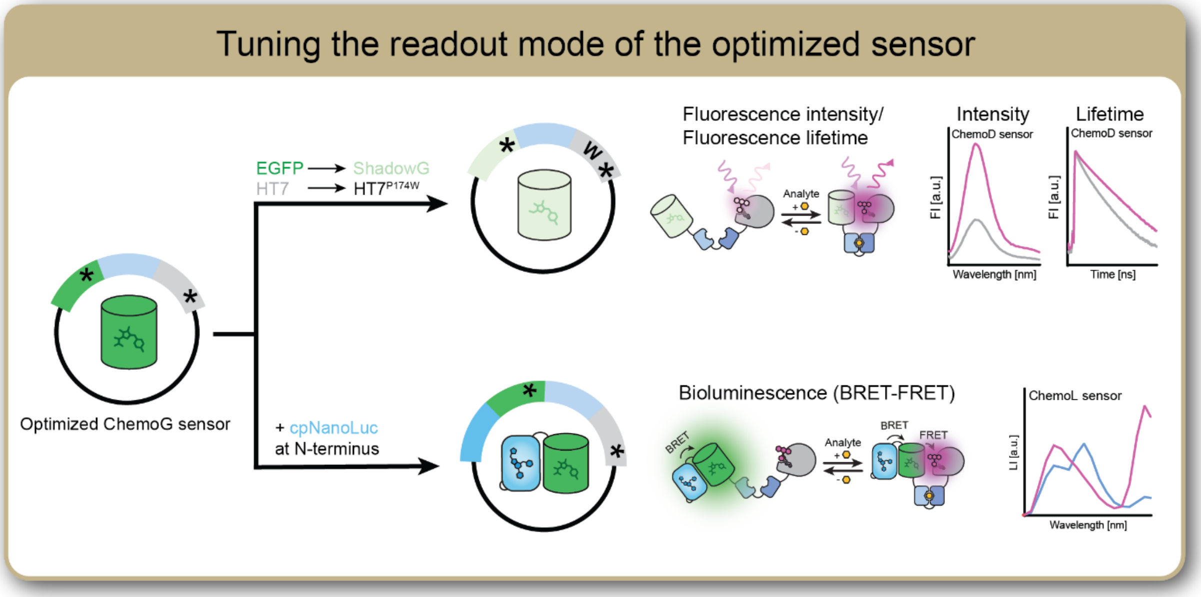

**Figure S 20| Tuning the readout mode of the optimized ChemoG sensor.**
See **Extended Note 3** for explanations.

#### Supplementary Tables

**Table S 1| FRET efficiencies of the ChemoG interface variants.**

| Construct | Interface mutations | | FRET ratio | FRET efficiency [%] |
|---|---|---|---|---|
| | EGFP | HaloTag7 | | |
| ChemoG1 | - | - | 2.2 ±0.1 | 74.8 ±0.4 |
| ChemoG2 | A206K | - | 4.0 ±0.1 | 84.1 ±0.6 |
| ChemoG3 | A206K | L271E | 8.9 ±0.1 | 90.9 ±0.1 |
| ChemoG4 | A206K | L271E-E143R-E147R | 11.6 ±0.3 | 93.2 ±0.1 |
| ChemoG5 | A206K-T225R | L271E-E143R-E147R | 20.3 ±0.8 | 95.8 ±0.1 |

FRET ratios (FRET/EGFP) and FRET efficiencies were determined for purified constructs labeled with SiR. Shown are the means ±s.d. (n = 3 technical replicates).

**Table S 2| FRET ratios of ChemoX constructs expressed in U-2 OS cells.**

| Construct | Subcellular localization | Localization tag | FRET ratio | Number of cells |
|---|---|---|---|---|
| HT7-EGFP | - | - | 0.1 ±0.05 | 9 |
| ChemoG1 | - | - | 3.7 ±0.3 | 18 |
| ChemoG2 | - | - | 5.7 ±1.1 | 15 |
| ChemoG3 | - | - | 8.7 ±1.4 | 20 |
| ChemoG4 | - | - | 13.6 ±2.2 | 29 |
| ChemoG5 | - | - | 16.4 ±2.7 | 32 |
| ChemoG5 | Cytosol | NES | 21.5 ±5.3 | 60 |
| ChemoG5 | Outer plasma membrane | $PDGFR_{tm}$ | 26.5 ±8.7 | 59 |
| ChemoG5 | Nucleus | NLS | 17.8 ±3.4 | 51 |
| ChemoG5 | Mitochondria | Cox8 | 16.3 ±7.0 | 126 |
| ChemoG5 | Nuclear envelope | Lamin B1 | 15.9 ±6.5 | 29 |
| ChemoB | - | - | 14.6 ±3.0 | 20 |
| ChemoC | - | - | 14.5 ±2.6 | 18 |
| ChemoY | - | - | 17.5 ±5.6 | 24 |
| ChemoR | - | - | 14.2 ±2.5 | 27 |

FRET ratios (FRET/FP) were determined for each construct expressed in U-2 OS cells labeled with SiR. Shown are the means ±s.d.

**Table S 3| FRET efficiencies of ChemoG5 labeled with different rhodamine fluorophores.**

| Construct | Fluorophore | Max emission [nm] | FRET ratio | FRET efficiency [%] |
|---|---|---|---|---|
| ChemoG5 | $JF_{525}$ | 556 nm | 18.0 ±1.4 | 94.9 ±0.3 |
| ChemoG5 | TMR | 580 nm | 23.6 ±2.7 | 96.6 ±0.3 |
| ChemoG5 | 580CP | 606 nm | 23.8 ±2.6 | 96.1 ±0.5 |
| ChemoG5 | CPY | 628 nm | 15.8 ±1.3 | 94.9 ±0.4 |
| ChemoG5 | SiR | 668 nm | 20.2 ±0.8 | 95.6 ±0.1 |
| ChemoG5 | $JF_{669}$ | 686 nm | 14.2 ±0.1 | 94.7 ±0.4 |

FRET/EGFP ratios and FRET efficiencies were determined for purified ChemoG5 labeled with different rhodamine fluorophores. Shown are the means ±s.d. (n = 3 technical replicates).

**Table S 4| FRET efficiencies of ChemoX FRET pairs.**

| Construct | FP | Interface mutations | | FRET ratio | FRET efficiency [%] |
|---|---|---|---|---|---|
| | | XFP | HaloTag7 | | |
| ChemoB | EBFP2 | N39Y-V206K-T225R | L271E-E143R-E147R | 36.2 ±0.3 | 96.6 ±0.1 |
| ChemoC* | mCerulean3 | T225R | L271E-E143R-E147R | 22.3 ±0.7 | 94.6 ±0.3 |
| ChemoG5 | EGFP | A206K-T225R | L271E-E143R-E147R | 20.3 ±0.8 | 95.8 ±0.1 |
| ChemoY | Venus | A206K-T225R | L271E-E143R-E147R | 22.4 ±1.9 | 96.6 ±0.1 |
| ChemoR | mScarlet | D201K | - | 8.4 ±0.2 | 91.3 ±0.3 |

FRET/FP ratios of purified ChemoX constructs were determined upon labeling with SiR. Shown are the means ±s.d. (n = 3 technical replicates). *mCerulean3 contains already K206, thus additional mutation at this position was not needed.

**Table S 5| Summarizing characteristics of the calcium sensors.**

| Construct | FP | Interface mutations | | | C50 | Max $\Delta R/R_0$ | Hill slope |
|---|---|---|---|---|---|---|---|
| | | # of mut. | XFP | HaloTag7 | | | |
| 1 | EGFP | 0 | - | - | 189 nM | 22.8 ±0.3 | 2.2 |
| 2 | EGFP | 1 | A206K | - | 203 nM | 33.3 ±0.8 | 1.8 |
| 3 (ChemoG-CaM) | EGFP | 2 | A206K | L271E | 179 nM | 36.1 ±1.0 | 2.2 |
| 4 | EGFP | 3 | A206K | L271E-E143R-E147R | 121 nM | 5.2 ±0.2 | 1.5 |
| 5 | EGFP | 4 | A206K-T225R | L271E-E143R-E147R | 207 nM | 0.8 ±0.1 | 1.1 |
| ChemoB-CaM | EBFP2 | 2 | N39Y-V206K | - | 206 nM | 12.7 ±0.2 | 1.8 |
| ChemoC-CaM | mCerulean3 | 1 | A206K | - | 158 nM | 2.3 ±0.1 | 3.2 |
| ChemoY-CaM | Venus | 1 | A206K | - | 226 nM | 21.7 ±0.6 | 2.0 |
| ChemoR-CaM0.1 | mScarlet | 1 | - | - | n.d. | 2.6 ±0.1 | n.d |
| ChemoR-CaM | mRuby2 | 0 | - | - | 202 nM | 3.4 ±0.1 | 2.7 |
| ChemoR-CaM0.2 | mRuby3 | 0 | - | - | n.d. | 2.5 ±0.1 | n.d. |
| ChemoR-CaM0.3 | mCherry | 0 | - | - | n.d. | 2.1 ±0.1 | n.d. |
| ChemoR-CaM0.4 | mKO2 | 0 | - | - | n.d. | 1.9 ±0.1 | n.d. |
| ChemoR-CaM0.4 | TagRFP | 0 | - | - | n.d. | 2.0 ±0.1 | n.d. |
| YC 3.6 | ECFP/Venus | - | - | - | 243 nM | 5.7 ±0.1 | 1.6 |

Maximum FRET/FP ratio changes ($^{Max}\Delta R/R_0$), C50 and Hill slope were determined for purified constructs. ChemoX-based calcium sensors were labeled with SiR. Values are based on titrations performed at 37 °C. Shown are the means and for $\Delta R/R_0$ also the standard deviations (n = 3-4 technical replicates).

**Table S 6| Summarizing characteristics of ChemoG-CaM labeled with different FRET acceptors.**

| Construct | Fluorophore | Max emission [nm] | C50 | Max $\Delta R/R_0$ | Hill slope |
|---|---|---|---|---|---|
| ChemoG-CaM | TMR | 580 nm | 66 nM | 3.9 ±0.1 | 1.4 |
| ChemoG-CaM | $JF_{585}$ | 610 nm | 100 nM | 10.5 ±0.4 | 1.5 |
| ChemoG-CaM | CPY | 628 nm | 76 nM | 8.6.0 ±0.1 | 2.2 |
| ChemoG-CaM | $JF_{635}$ | 656 nm | 114 nM | 24.4 ±0.3 | 2.5 |
| ChemoG-CaM | SiR | 668 nm | 179 nM | 36.8 ±0.2 | 2.2 |

Maximum FRET/EGFP ratio changes ($^{Max}\Delta R/R_0$), C50 and Hill slope were determined for purified ChemoG-CaM labeled with different fluorophores. Values are based on titrations performed at 37 °C. Shown are the mean and for $\Delta R/R_0$ also the standard deviations (n = 3 technical replicates).

**Table S 7| Summarizing characteristics of ATP sensors.**

| Construct | FP | Interface mutations | | C50 | Max ΔR/$R_0$ | Hill slope |
|---|---|---|---|---|---|---|
| | | XFP | HaloTag7 | | | |
| 1 | EGFP | A206K | - | N.D | 9.9 ±0.1 | N.D |
| 2a (ChemoG-ATP) | EGFP | A206K | L271E | 2.3 mM | 12.1 ±0.4 | 1.4 |
| 2b | EGFP | A206K-T225R | - | N.D. | 6.0 ±0.1 | N.D |
| 3 | EGFP | A206K-T225R | L271E | N.D. | 1.9 ±0.0 | N.D |
| ChemoB-ATP | EBFP2 | N39Y-V206K | L271E | 2.8 mM | 5.0 ±0.1 | 1.6 |
| ChemoR-ATP | mRuby2 | - | - | 3.2 mM | 0.8 ±0.1 | 2.0 |
| ATeam 1.03 | mseCFP/cpVenus | - | - | 1.8 mM | 1.4 ±0.1 | 1.8 |

Maximum FRET/FP ratio changes ($^{Max}\Delta R/R_0$), C50 and Hill slope were determined for purified constructs. ChemoX-based ATP sensors were labeled with SiR. Values are based on titrations performed at 37 °C. Shown are the mean and for $\Delta R/R_0$ also the standard deviations (n = 3 technical replicates).

**Table S 8| Summarizing characteristics of NAD$^+$ sensors.**

| Construct | FP | Affinity mutation *tt*LigA | Interface mutations | | Fluo | C50 | Max ΔR/$R_0$ | Hill slope |
|---|---|---|---|---|---|---|---|---|
| | | | XFP | HaloTag7 | | | | |
| 1 | EGFP | - | A206K | - | TMR | 38 µM | 10.1 ±0.1 | 1.6 |
| 2 | EGFP | V292A | A206K | - | TMR | 75 µM | 6.2 ±0.1 | 1.2 |
| 3 | EGFP | Y226W | A206K | - | TMR | 129 µM | 6.3 ±0.1 | 1.2 |
| 4 | EGFP | Y226W-V292A | A206K | - | SiR | 205 µM | 2.0 ±0.1 | 0.9 |
| 5 | EGFP | Y226W-V292A | A206K-T225R | - | SiR | 167 µM | 18.1 ±0.3 | 1.0 |
| 6 (ChemoG-NAD) | EGFP | Y226W-V292A | A206K-T225R | L271E | SiR | 200 µM | 34.7 ±0.4 | 0.8 |
| ChemoG-NAD | EGFP | Y226W-V292A | A206K-T225R | L271E | TMR | 136 µM | 7.5 ±0.1 | 1.0 |
| ChemoG-NAD | EGFP | Y226W-V292A | A206K-T225R | L271E | $JF_{585}$ | 36 µM | 18.5 ±0.1 | 0.9 |
| ChemoG-NAD | EGFP | Y226W-V292A | A206K-T225R | L271E | CPY | 117 µM | 20.4 ±0.1 | 0.8 |
| ChemoG-NAD | EGFP | Y226W-V292A | A206K-T225R | L271E | $JF_{635}$ | 52 µM | 22.5 ±0.1 | 0.9 |
| 7 | EGFP | Y226W-V292A | A206K-T225R | L271E- E143R-E147R | SiR | 25 µM | 32.5 ±0.3 | 0.8 |
| ChemoB-NAD | EBFP2 | Y226W-V292A | N39Y-A206K-T225R | L271E | SiR | 103 µM | 11.2 ±0.1 | 0.9 |
| ChemoR-NAD | mRuby2 | Y226W* | - | - | SiR | 78 µM | 3.0 ±0.1 | 1.0 |

Maximum FRET/FP ratio changes ($^{Max}\Delta R/R_0$), C50 and Hill slope were determined for purified constructs labeled with indicated fluorophore substrates. Values are based on titrations performed at 37 °C. Shown are the mean and for $\Delta R/R_0$ also the standard deviations (n = 3 technical replicates).

**Table S 9| Summarizing characteristics of intensiometric NAD+ sensors.**

| Construct | Fluorophore | Max emission [nm] | C50 | Max ΔFI/$FI_0$ | Hill slope |
|---|---|---|---|---|---|
| ChemoG-NAD | SiR | 666 nm | 21.0 µM | 28.0 ±1.9 % | 0.89 |
| ChemoD-NAD | SiR | 666 nm | 32.7 µM | 161.1 ±5.0 % | 0.84 |
| ChemoD-NAD | CPY | 628 nm | 36.8 µM | 104.7 ±1.2 % | 0.83 |
| ChemoD-NAD | $JF_{635}$ | 662 nm | 47.5 µM | 226.6 ±4.3 % | 0.59 |

Maximum fluorescence intensity changes ($^{Max}\Delta FI/FI_0$), C50 and Hill slopes were determined for purified constructs labeled with the indicated fluorophores. Values are based on titrations performed at 37 °C. Shown are the means and for $\Delta F/F_0$ also the standard deviations (n = 3 technical replicates).

**Table S 2| Summarizing characteristics of fluorescence lifetime-based NAD+ sensors.**

| Construct | Fluorophore | Max emission [nm] | C50 | MaxΔτ | Hill slope |
|---|---|---|---|---|---|
| ChemoG-NAD | SiR | 666 nm | 14.2 µM | 0.53 ±0.03 ns | 1.34 |
| ChemoD-NAD | SiR | 666 nm | 22.4 µM | 1.16 ±0.01 ns | 0.99 |
| ChemoD-NAD | CPY | 628 nm | 44.6 µM | 1.18 ±0.01 ns | 0.91 |
| ChemoD-NAD | $JF_{635}$ | 662 nm | 32.3 µM | 0.77 ±0.01 ns | 0.68 |

Maximum intensity-weighted average fluorescence lifetime changes (MaxΔτ), C50 and Hill slopes were determined for purified constructs labeled with the indicated fluorophores. Values are based on titrations performed at 37 °C. Shown are the means and for MaxΔτ also the standard deviations (n = 3 technical replicates).

**Table S 3| ChemoG FRET pairs recommended for the development of ChemoG FRET biosensors.**

| Construct | Interface mutations | | Addgene# |
|---|---|---|---|
| | EGFP | HaloTag7 | |
| ChemoG1 | - | - | 193799 |
| ChemoG2 | A206K | - | 193800 |
| ChemoG3 | A206K | L271E | 193801 |
| ChemoG3.1 | A206K-T225R | - | 193802 |
| ChemoG3.2 | A206K-T225R | L271E | 193803 |
| ChemoG5 | A206K-T225R | L271E-E143R-E147R | 193805 |

#### Extended notes

##### Extended note 1 – Development of ChemoG biosensors.

***Generation of sensor variants*.** Certain ChemoG interface mutations increase FRET to a larger extent than others. For example, the interface mutation T225R$^{EGFP}$ usually leads to a stronger FRET increase than the interface mutation L271E$^{HT7}$. This feature revealed useful to fine-tune the dynamic range of ChemoG-based sensors. For the generation of new sensors (**Fig. S18**), we recommend to try a palette of ChemoG FRET pairs with different interface mutations (**Table S11**, available on Addgene). The sensing domain can be derived from an existing biosensor as *e.g.* ChemoG-CaM that was derived from YC 3.6[7] or a new sensing domain, preferentially exhibiting a large conformational change. To create ChemoG sensor variants, the sensing domain should be cloned between the EGFP and HaloTag7 variants (*i.e.* ChemoG FRET pairs). Using ChemoG-encoding plasmids and DNA encoding the sensing domain of interest, 6 plasmids encoding sensor variants can simply be obtained through PCR and molecular cloning (*e.g.* by Gibson assembly[1]). We recommend to use single GGS linkers connecting the ChemoG FRET pairs with the sensing domain but these can also be further engineered in a second step if necessary. The linkers can be created during the design of the primers used for the PCR amplification of the fragments. We deposited plasmids encoding ChemoG variants for protein production in *E. coli*. In case the sensor variants should be tested in mammalian cells, the vector backbone should first be exchanged.

***Testing sensor variants in vitro or in cells*.** Two options are available:

- produce the sensor variants in *E. coli*, purify them and test them *in vitro*, or
- express and test the sensor variants in mammalian cells (require extra sub-cloning, see above).

For the first option, the purified sensor variants should be labeled with an orange/red fluorophore substrate. We recommend SiR-halo or a rhodamine substrate with similar spectral properties to minimize direct excitation of the synthetic fluorophore. The labeled sensors should then be titrated with different concentrations of an analyte of interest (AOI). The sensor variant with the largest dynamic range can be identified from fluorescence emission spectra. An ideal sensor exhibits low FRET in absence and high FRET in presence of the AOI (or *vice versa*), showing a large peak inversion in each emission channel. Some noticeable fluorescence should remain in both channels for precise measurements.

For the second option, mammalian cells should be transfected with plasmids encoding the sensor variants. The transfected cells should be labeled with cell-permeable fluorophore substrates. As previously, we recommend SiR-halo. Labeled cells can subsequently be treated with reagents known to act on the biological activity of interest (*e.g.* AOI concentration change). Via fluorescence microscopy or flow cytometry, the fluorescence profile of treated and untreated cells can be compared to identify sensor variants with the largest dynamic range. Sensors presenting noticeable fluorescence signal in both channels in presence and absence of treatment should be chosen in order to ensure precise measurement. Technical details on how to conduct the different experiments can be found in the method section of the manuscript.

**Extended note 2 – Tuning the spectral properties of the optimized ChemoG sensor.**

The spectral properties of the optimized ChemoG sensor can be tuned by exchanging the FRET donor EGFP with other fluorescent proteins and/or by using different fluorophore substrates as FRET acceptor (**Fig. S19**). For exchanging the FRET donor, EGFP is substituted with an alternative fluorescent protein (*e.g.* EBFP2 = ChemoB) with the same interface mutations (*e.g.* $EGFP^{T225R}$ → $EBFP2^{T225R}$) via molecular cloning. As FRET donor, we recommend EGFP-derived fluorescent proteins such as EBFP2, mCerulean3 or Venus to ensure a good transferability of the interface mutations. We recommend to use FP constructs we deposited on Addgene to ensure that the adequate FP mutations are used. For red fluorescent proteins, we recommend using mRuby2 without additional mutations for biosensor design. The FRET acceptor can be readily chosen by simply labeling the ChemoX sensors with different rhodamine-based HaloTag substrates (*e.g.* $JF_{525}$, CPY or $JF_{669}$). The ChemoX sensors performance can be evaluated as explained in **Extended note 1** and in the methods.

**Extended note 3 – Tuning the readout mode of the optimized ChemoG sensor.**

The readout of ChemoG FRET sensors can be tuned by small modifications (**Fig. S20**). Single channel fluorescence intensity and fluorescence lifetime-based ChemoD sensors are obtained by substituting EGFP with its non-fluorescent variant ShadowG[40] carrying the same interface mutation(s). Additionally, the fluorescence quenching mutation P174W should be introduced into HaloTag7. For intensiometric sensors, we recommend labeling with $JF_{635}$ while for fluorescence lifetime imaging, CPY worked best in our hands so far. The performance of the sensors can be evaluated analogously as explained in **Extended note 1** and in the methods.

To convert ChemoG FRET sensors into a bioluminescent ChemoL sensor, a circularly permuted variant of NanoLuc is fused to the N-terminus of EGFP. We recommend labeling the sensor with rhodamine fluorophore substrates whose spectral properties are compatible with the available equipment. In our case, CPY was the most red-shifted fluorophore compatible with our plate reader but we foresee no conceptual hurdle in using any rhodamine fluorophore substrate proven functional for FRET biosensing. The sensors performance can be evaluated analogously as explained in **Extended note 1** and in the methods. It should be noted, that the expression of ChemoL sensors in mammalian cells revealed substantially lower than the corresponding ChemoG FRET sensors. While this does not affect the luminescent readout, it is not advisable to use ChemoL sensors for FRET applications even if this is conceptually possible.
